# Structure-enabled enzyme function prediction unveils elusive terpenoid biosynthesis in archaea

**DOI:** 10.1101/2024.01.29.577750

**Authors:** Raman Samusevich, Safa Mert Akmeşe, Téo Hebra, Roman Bushuiev, Martin Engst, Jonáš Kulhánek, Anton Bushuiev, Joshua D. Smith, Tereza Čalounová, Helena Smrčková, Marina Molineris, Renana Schwartz, Adrian Svoboda, Ratthachat Chatpatanasiri, Sotirios C. Kampranis, Dan Thomas Major, Josef Sivic, Tomáš Pluskal

## Abstract

The exponential growth of uncharacterized enzyme sequences in genomic repositories demands novel tools for functional annotation. Here, we combined alignment-driven structural domain analysis with protein language models to create EnzymeExplorer, a machine-learning pipeline for enzyme function prediction. We applied this approach to terpene synthases (TPSs), which present an ideal model case because they catalyze complex carbocationic rearrangements with unpredictable product outcomes. We detected new structural domains and achieved significantly higher average precision than existing methods for function prediction. By analyzing the UniRef90 database, we identified TPSs overlooked by existing computational methods. Remarkably, we discovered and experimentally confirmed three archaeal TPSs, expanding the known taxonomic distribution of TPS catalysis to a new domain of life. Further in silico screening of archaeal proteomes revealed that terpene biosynthesis is widespread across Archaea. Our approach offers a powerful framework for characterizing enzyme “dark matter” in the rapidly expanding genomic and metagenomic datasets.

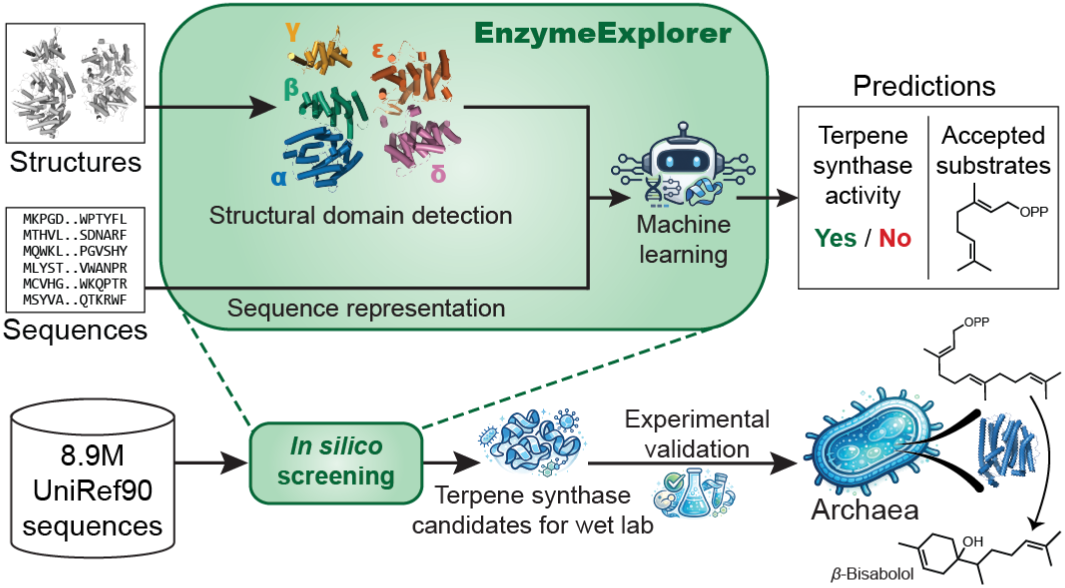

## Main text

The number of protein sequences in genomic repositories is increasing exponentially, making experimental characterization of all discovered proteins unfeasible. Bioinformatic signature scanning tools such as InterProScan^1^ can provide partial sequence annotations, including conserved domains, sequence motifs, and binding sites. However, for enzymes, these annotations typically offer only broad functional classifications (e.g., “*O*-methyltransferase”) without revealing additional biochemical context, such as substrates or reaction products. Moreover, over 40 million protein sequences in the UniProt database^2^ currently lack InterPro annotations^3^ entirely. This unannotated “dark matter” of protein sequence space likely harbors millions of uncharacterized enzymes, presenting both a barrier to biological research and an untapped resource for biotechnology^4^. While current state-of-the-art methods for functional annotation of enzymes^5,6^ perform well at identifying previously reported enzymatic activities, it is difficult for them to annotate protein sequences distant from training data. Thus, enhanced, contextually-rich predictions for uncharacterized enzyme functions are needed to facilitate efficient exploration of enzyme diversity within the uncharted protein sequence space.

Computational prediction of enzyme functions without recognizable sequence signatures remains a formidable scientific challenge^7^. To address this challenge, we developed a new predictive approach that leverages protein structure prediction to provide informative features for machine learning. Structural information can help overcome the sparsity of sequence space, as structure is known to evolve slower than sequence^8^, and catalytically similar enzymes with low sequence similarity share common folds^9^. Thus, we set out to build a predictive pipeline that integrates structure-based and language-based representations of protein sequences. To implement our approach, we focused on a model enzyme family that exemplifies the current challenges in sequence-based function prediction.

Terpene synthases (TPSs) are widely distributed enzymes responsible for producing the core hydrocarbon scaffolds of terpenoids, the largest class of natural products^10^. With over 100,000 known molecular structures, terpenoids include widely used flavors, fragrances, and first-line medicines, such as the antimalarial drug artemisinin and the chemotherapy agent paclitaxel^11,12^. TPSs catalyze complex cyclization reactions on isoprenoid diphosphate substrates composed of C_5_ isoprene units (e.g., geranyl pyrophosphate C_10_ or farnesyl pyrophosphate C_15_, see overview in **Fig. S1**), often yielding multiple products with fused rings and intricate stereochemistry^10^. The complex reaction mechanism of a TPS involves a cascade of carbocationic rearrangements in which active-site residues shape the substrate into a catalytically productive conformation and stabilize the highly reactive carbocation intermediates, thereby directing the reaction toward specific products^10,13^. TPS active sites are mostly lined by chemically “inert” residues and substitution of even a single residue has been shown to disrupt the precise choreography of the carbocationic interconversion to result in synthases with drastically different product profiles^14–16^. These minor substitutions, only changing the active site contour by a few angstroms, have profound outcomes that are difficult to predict by sequence information alone. This mechanistic complexity positions TPSs as an ideal model enzyme class for applying machine learning approaches^17^.

Here, we present a novel machine learning pipeline that combines alignment-driven structural analyses with a protein language model to predict TPS enzymatic activity and substrate preference. We demonstrate that this approach outperforms existing tools developed for enzyme detection and classification, and generalizes well to distant homologs. As a result, we are able to identify previously undetected structural domain subtypes and to discover and experimentally characterize several new functional TPSs overlooked by current annotation methods. Remarkably, we reveal that TPSs are widespread in archaea and functionally characterize the first TPSs ever described from this domain of life.

### Structural analysis reveals fine differences in domain configurations

The goal of predictive discovery of family-specific enzymatic activity is to utilize a small dataset of previously biochemically characterized enzymes from that family to detect the activity in larger datasets of uncharacterized protein sequences. The quality of the training dataset is critical for the effectiveness of the prediction. We used MARTS-DB^18^ as the primary dataset for TPSs, which includes 1,256 experimentally characterized TPS enzymes that catalyze 3,852 corresponding reactions (**Supplementary File 1**). This dataset provides robust coverage of monoterpene synthases (monoTPSs), sesquiterpene synthases (sesquiTPSs) and diterpene synthases (diTPSs; as defined in **Fig. S1**), but includes relatively fewer examples of less studied TPS classes, such as sesterTPSs or tetraTPSs (**Figs. S2a**-**b**). It also includes 181 examples of isoprenyl diphosphate synthases (IDS). Because IDSs are close relatives of TPSs^10^ we utilized them as “hard negatives” (enzymes that share significant structural and sequence similarity with TPSs, yet do not catalyze TPS reactions).

To construct our predictive pipeline, we set out to develop structure-based representations of the input protein sequence and combine them with language-based representations (**Figs. 1a, 2a**). Since enzymes in general comprise one or more structural domains^19^ and the organization of these domains contributes to determining enzymatic function, we studied the domain configurations of the enzymes in MARTS-DB to provide informative features for machine learning. TPS enzymes feature a modular protein architecture, composed of structural domains known as α, β, and γ^10,17^. Thus, for the structure-based component of our pipeline, we leveraged AlphaFold 3^20^ and ESMFold^21^ to predict the protein structures, and designed an approach that segments the predicted structures into structural domains (**Fig. S3**). This process generates a structural domain configuration of each input protein sequence.

**Fig. 1.**
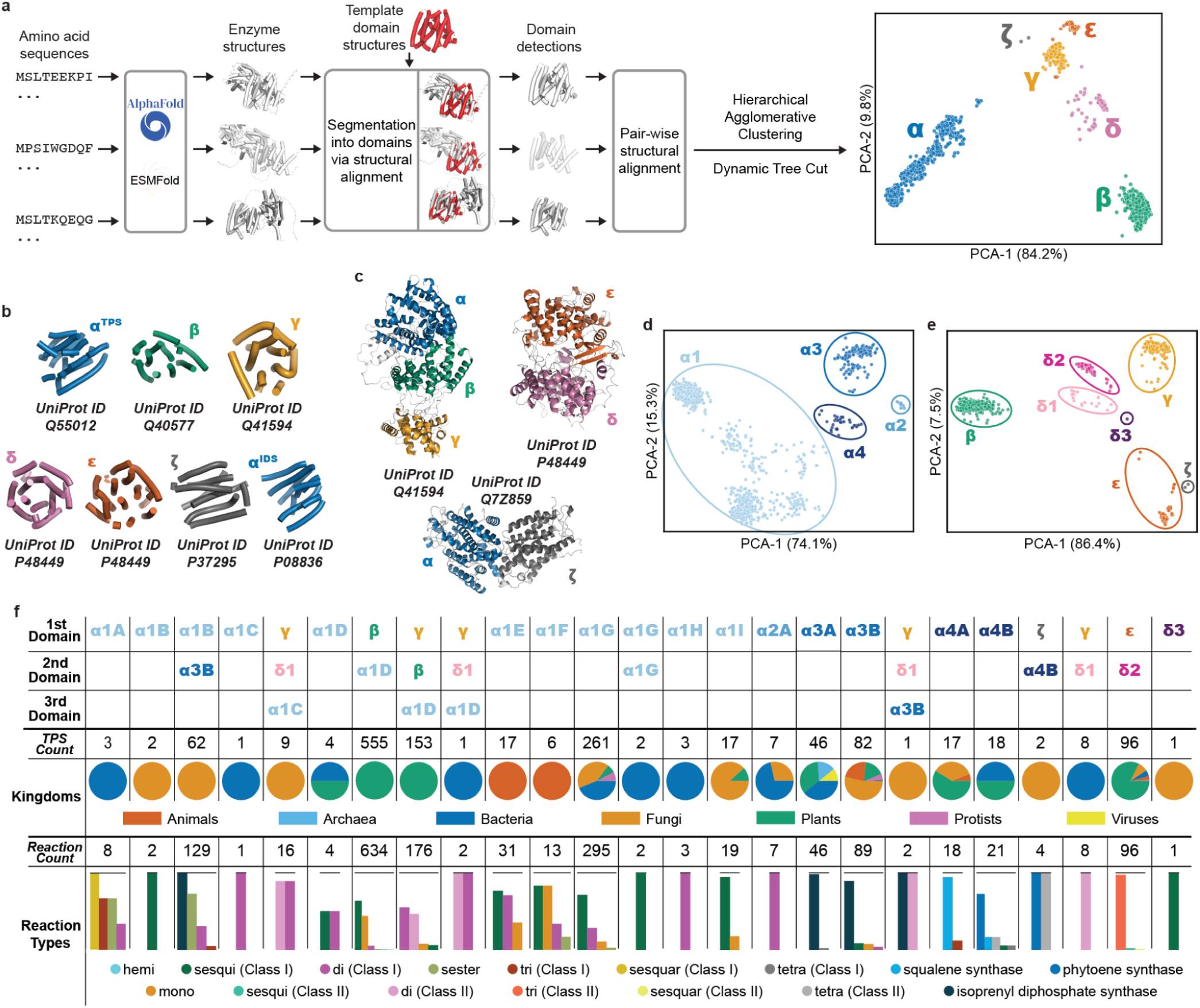
Discovery and analysis of structural domain types and subtypes using a segmentation-clustering pipeline. **a**, Overview of our structure-based workflow. Predicted protein structures are segmented into family-specific domains via structural alignment. These detected domains are then clustered, revealing distinct structural domain groups. **b**, Representative α, β, γ, δ, ε, and ζ TPS domains. The α domain is represented by both a TPS and an IDS to illustrate the structural variation among α domains (α^TPS^ vs α^IDS^). Experimental crystal structures of α^34,35^, β^36^, γ^37^, δ^38^ and ε^38^ domains obtained from the RCSB PDB^39^ are shown, while an AlphaFold 3^20^ (AF3)-predicted structure of ζ domain is shown, for which no experimental structure is available. **c**, The highlighted structures of typical α, β, γ domains in the context of the diTPS taxadiene synthase (UniProt ID Q41594, PDB ID 3P5R^37^), the typical δ and ε domains in the context of a triTPS lanosterol synthase (UniProt ID P48449, PDB ID 1W6J^38^), and the typical α and ζ domains in the context of a bifunctional lycopene cyclase/phytoene synthase (UniProt ID Q7Z859, AF3 predicted structure). **d**, Principal component analysis (PCA) of clustering of all detected α domains. **e**, PCA of clustering of all detected non-α domains, highlighting the three distinct subtypes of the δ domain. **f**, Structural domain configurations of 1,374 validated natural TPS/IDS enzymes in MARTS-DB^18^, including the taxonomy distribution and TPS reaction types for each domain configuration. Please note that if two different enzymes counted in the ***TPS Count*** row catalyze the same reaction (substrate/product pair), these are counted independently as two reactions in the ***Reaction Count*** row. ***Reaction Types*** are represented in bar plots, solid lines at the top stand for the total number of reactions of the corresponding domain configuration (***Reaction Count***).

**Fig. 2.**
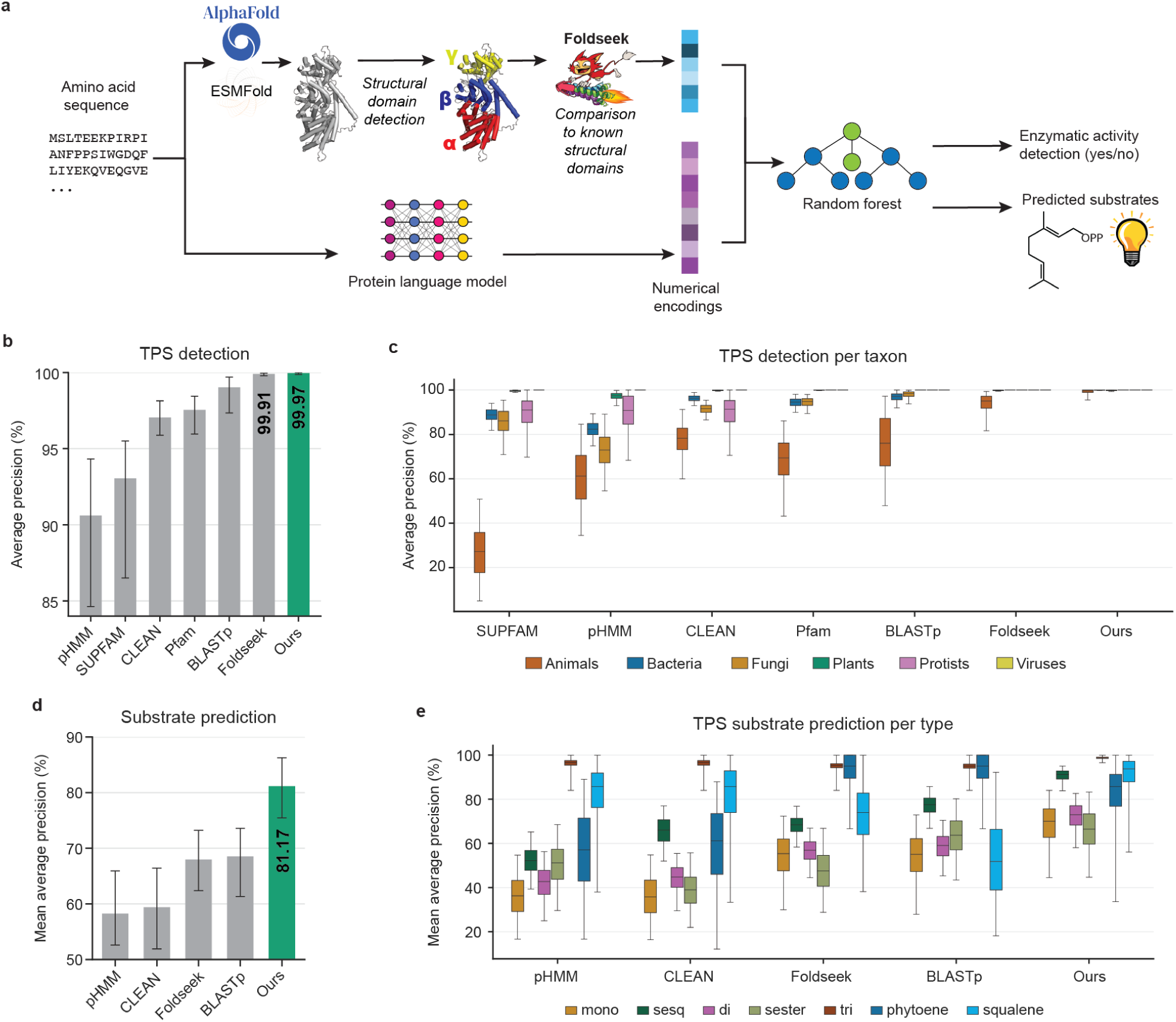
Accurate enzyme function prediction using EnzymeExplorer. **a**, Overview of our predictive pipeline. Predicted protein structures are segmented into enzyme-specific domains, which are compared to domains of the same type within previously characterized enzymes from the same family. Similarities to characterized domains, quantified as TM-scores, are recorded as features for downstream analysis. Protein sequences are also processed by a protein language model to extract embeddings, which are retrieved by mean pooling of the final layer’s per-residue embeddings. Finally, the concatenated TM-score features and embeddings are fed into random forest classifiers to predict the presence of enzymatic activity and classify substrates. **b**, *In silico* evaluation of TPS detection performance was measured using average precision. Individual precision-recall curves are in **Fig. S7**. Each method is optimized/retrained on the EnzymeExplorer dataset. **c**, Taxon-specific performance of our approach for TPS detection compared to other methods. **d**-**e**, Substrate prediction performance, highlighting the efficacy of our approach compared to existing methods across all types, including underrepresented minority TPS types like sesterTPS. Additional comparisons using other metrics are provided in **Fig. S7**. In all figures, the absolute performance values are calculated using the pooled out-of-fold predictions and error bars represent the 95% bias-corrected accelerated (BCa) confidence intervals^51^ from a block-bootstrapping with 1000 iterations.

We applied this approach to our dataset of 1,374 characterized TPS/IDS enzymes and constructed the unweighted pair group method with arithmetic mean^22,23^ (UPGMA) dendrogram from pairwise structural-similarity TM-scores calculated for all detected domains (**Fig. S4a**). UPGMA is a hierarchical agglomerative clustering (average-linkage) algorithm, widely used in computational biology for phylogenetics^24,25^ and biological classification analyses^26–29^. This analysis recovered the established α, β and γ structural domain types and remarkably separated additional clusters that we label δ, ε and ζ (**Fig. 1a**, PCA analysis on the right). δ and ε correspond to lanosterol-synthase-like β/γ regions, whereas ζ is a provisional group represented by two predicted structures (**Fig. 1c)**. All six domain types are structurally distinct both when they are aligned separately (**Figs. 1b**) and when they are compared in the context of the full-length TPS proteins (**Fig. 1c**). Visual inspection of the UPGMA dendrogram **(Fig S4a)** revealed distinct, well-separated branches within the major structural domain classes, suggesting the presence of finer-scale structural subtypes. However, these branches showed heterogeneous heights and nested subclades, and applying a single global cut height would either over-merge or over-split structurally coherent groups. We therefore applied the Dynamic Tree Cut algorithm^30^ to identify the structural domain subtypes. Unlike cutting the dendrogram at a single fixed height, Dynamic Tree Cut adapts to the local shape and density of individual branches, allowing clusters with different levels of compactness and separation to be detected, which ultimately revealed distinct subtypes of these main structural domain types (**Figs. S4a-b**). Specifically, the α domain structures can be further separated into 14 distinct subtypes, which we designated α1A to α4B (**Figs. 1f, S4a**), while the newly identified δ domain structures contain three distinct subtypes, designated δ1, δ2, and δ3.

We further investigated the ζ domain due to its marginally distinct structure in comparison to the rest of the major domain types (**Fig. S4a**). Even though there are only two structural predictions of TPSs containing this domain type (UniProt ID Q7Z859 and P37295), the evidence shows that these are related to class II tetraTPS reactions (**Fig. 1f**), specifically carotenoid biosynthesis^31,32^. Notably, this structural domain has not been reported in CATH-DB^33^.

Overall, we were able to distinguish 25 existing TPS domain configurations (**Fig. 1f** and **Supplementary File 2**), further supported by phylogenetic analysis (**Fig. S6**). The domain configurations provide insights into the functional diversity of TPS structures: for example, all known tetraTPSs have the α4B domain, while the α3 domain types are associated with the IDS reactions. Moreover, the β domain type can be found only in plants. In addition, the δ and ε domains are primarily associated with class II triTPSs, being the only structural domains synthesizing lanosterol or cycloartenol, the precursors of steroidal compounds.

### Predictive model for enzyme functional annotation

The key challenge in predicting TPS enzymatic activities lies in designing a predictor that (i) generalizes well to unseen protein sequences, especially with limited training data as some TPS classes have only a handful of characterized sequences, and (ii) can distinguish fine-grained structural differences between the different TPS classes. To address this challenge, we have developed a pipeline that combines alignment-driven structural analysis with a protein language model (**Fig. 2a**). Relying solely on sequence-based evolutionary embeddings often fails to capture the precise 3D geometric constraints required for complex carbocationic cascades, while pure structural models may miss deep evolutionary context. Therefore, we engineered a combined feature space. In detail, the similarity of detected structural domains to each recognized TPS domain template (**Fig. 1b**) is analyzed using the Foldseek^40^ algorithm. This set of similarities, quantified as TM-scores, is then encoded into a numerical vector representing the structural information. This representation enables encoding fine-grained structural information specific to the given enzyme class (here TPS). In the language-based component, we employed the pretrained large protein language model Ankh-large^41^, which provided an additional numerical vector encoding the sequence information. Leveraging a strong protein language model supports generalization to previously unseen sequences by providing evolutionary information^41–43^. For each protein, we computed a single sequence embedding by mean-pooling the final hidden layer across residue tokens, and this embedding was used as the sequence vector representation. We used the concatenated structural and sequence vector representations of the input protein as an input feature for a random forest classifier^44^ that predicts the presence or absence of TPS enzymatic activity and the substrate of the enzyme (**Fig. 2a**, right side). Given the extreme sparsity of data for certain TPS classes, a random forest was deliberately chosen over high-capacity neural network architectures to mitigate overfitting and robustly handle the high-dimensional concatenated features. The random forest classifier was trained in a supervised multi-label framework using our curated dataset of 3,852 TPS reactions as positives, 333 IDS reactions as hard negatives (**Supplementary File 1**), and 10,000 additional negative examples (non-TPS protein sequences), which were sourced from Swiss-Prot^45^ by extensive hard negative mining. The complete dataset was divided into five folds for cross-validation with a 50% maximum sequence identity and 80% minimum coverage using MMSeqs2^46^ (**Supplementary File 4).** Finally, we appended a calibration module on top of the random forest classifier to yield the probability estimates of the predictions in deployment (**Tables S7, S8**).

Having established our pipeline, we investigated its performance in predicting enzyme function by comparing it to well-established profile hidden Markov models (pHMMs) in the Pfam^47^ and SUPFAM^48^ databases as well as to TPS-specific pHMMs retrained^49^ on our own TPS dataset (**Supplementary File 1**). As additional baselines, we used the protein sequence search algorithm BLASTp^50^, the protein structure search algorithm Foldseek^40^, and the state-of-the-art EC number predictor CLEAN^6^. Each of these baselines were optimized and fine-tuned separately for each prediction task using the 5-fold cross-validation dataset (**Supplementary File 4**). Compared to all these methods, our approach consistently achieved the highest average precision for TPS detection across diverse taxa (**Figs. 2b-c**) and the most accurate multi-label substrate predictions across different TPS types (**Figs. 2d-e**), including underrepresented TPS classes such as sesterTPSs. We also performed pooled out-of-fold block-bootstrapping to prove the statistical significance of performance differences between EnzymeExplorer and the rest of the methods, showing that EnzymeExplorer performed significantly better than the baseline methods for both TPS detection and substrate prediction tasks (**Fig. S7**). The detailed methodology, baselines, validation scheme, evaluation metrics, and ablation studies supporting these results are outlined in the **Materials and Methods** and in **Supplementary Information** (**Figs. S7, S8, S9**).

### Experimental validation of model predictions

To validate the performance of our machine learning approach for the discovery and characterization of novel TPSs, we experimentally characterized the enzymatic activity of selected candidate synthases. To functionally characterize these enzymes, we heterologously expressed the corresponding genes in the budding yeast *Saccharomyces cerevisiae* strain JWY501^52^. This strain has been engineered for elevated production of the sesquiTPS and diTPS substrates farnesyl diphosphate (FPP) and geranylgeranyl diphosphate (GGPP), respectively.

In the first validation experiment, we mined uncharacterized protein sequences that carry PFAM or SUPFAM signatures characteristic for TPSs (**Supplementary File 3**). We populated the phylogenetic tree of these sequences together with the previously characterized TPSs in MARTS-DB^18^ and used it to calculate the phylogenetic distances between uncharacterized sequences and the characterized TPSs by summing the branch lengths. Finally, we selected nine sequences as experimental candidates that were phylogenetically most distant from characterized TPSs in MARTS-DB (**Fig. S10a, Table S1**). Using gas chromatography mass spectrometry (GC-MS) and liquid chromatography mass spectrometry (LC-MS) we confirmed that out of the nine tested enzymes, six exhibited production of either sesquiterpene or diterpene compounds. As the final step, we ran EnzymeExplorer for these nine candidates, which was able to predict the substrates of four out of six experimentally verified candidates correctly (**Table S1**), confirming that our algorithm can provide reliable predictions on phylogenetically distant biological sequences (**Figs. S11-S14**).

To further assess our method’s ability to facilitate novel discoveries in unexplored parts of protein sequence space, we aimed to identify “*impossible to discover”* TPSs, focusing on cases where existing bioinformatic methods would either completely miss these proteins or yield low-precision predictions. We applied EnzymeExplorer to sequences from the UniRef90^53^ database that have a predicted structure in AlphaFold Protein Structure Database^54^ and lack any known InterPro signatures^3^, including any “domain of unknown function” (**Fig. S10b**, screening results available on Zenodo). For experimental validation, we filtered the predictions with >95% predicted TPS probability and selected enzymes originating from organisms where terpenoid biosynthesis has been less studied or unreported, such as bacteria, viruses, and archaea. To this end, we sampled 11 sequences from these taxa with high phylogenetic distance to the known TPSs in MARTS-DB (**Fig. S10c**). Following the heterologous expression of 11 candidates in *S. cerevisiae*, we confirmed that seven candidates exhibited TPS activity (**Figs. S15-S18, S23a-b**). Among these highly unconventional protein sequences, we identified an unusual viral TPS (A0A2P0VN22), which cannot be detected by both sequence and structure alignment methods (**Table S6**). Notably, we identified three active TPS enzymes in archaea (A0A0E3NXY0, A0A537EJD0, A0A5E4I9B1), a domain of life with no previous TPS reports until now^55^. The sequence alignment methods failed to identify these archaeal TPSs except A0A5E4I9B1 (**Table S6**), which could be identified by Pfam using the TPS-tuned bit-score threshold (see **Baselines** in **Performance evaluation of predictive models**). However, with the default InterProScan^1^ thresholds, Pfam fails to annotate this sequence as a TPS.

To further validate the finding of functional TPSs in archaea, we selected the most active of the identified TPSs, A0A5E4I9B1 (**Fig. S23b**), to structurally characterize its main product. GC–MS extracted-ion chromatograms (**Fig. S23b**) revealed seven peaks (8–14) that were absent from the control, consistent with the well-documented multiproduct behavior of TPSs^56,57^. Among these, peak 12 was the most abundant and considered as the main product. To enhance terpene production in yeast, we transitioned from genomic integration to plasmid-based expression of the A0A5E4I9B1 gene. This enabled the isolation of the main product and its identification as *β*-bisabolol by nuclear magnetic resonance (NMR) spectroscopy (**Figs. S19, S23c**), which aligns with the recently discovered plant TPS producing the same compound^58^. The remaining peaks (8–11, 13, 14) are minor products detected reproducibly across replicates; while not assigned individually here, their presence is consistent with the known tendency of TPSs to generate product mixtures^56,57^. We further validated the TPS activity of A0A5E4I9B1 through heterologous expression of the protein in *E. coli* and an *in vitro* assay with FPP, yielding *β*-bisabolol as the main product (**Figs. S26, S27**).

**Table 1.**
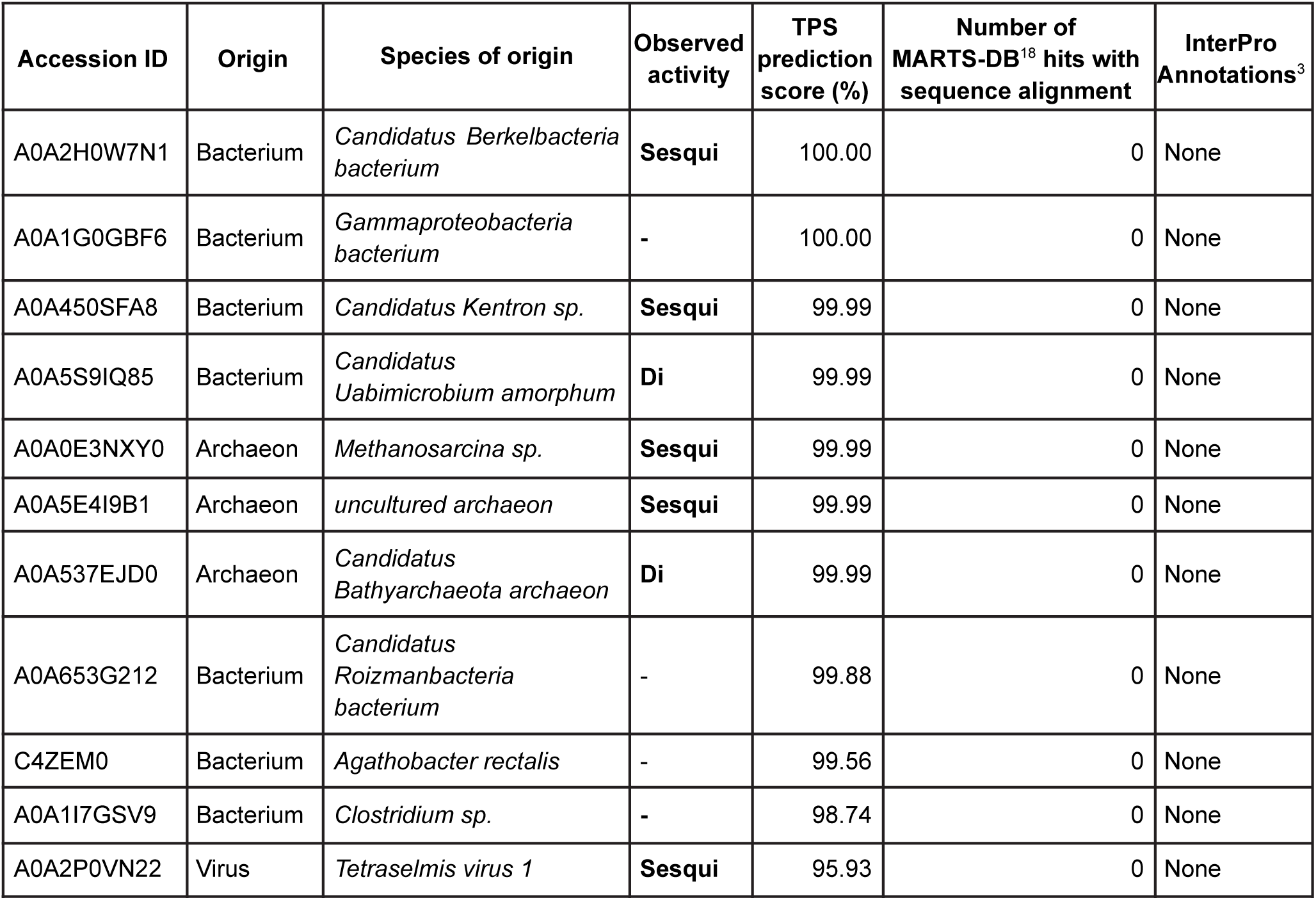
List of candidate TPS sequences expressed in yeast. Sequences were selected based on EnzymeExplorer predictions from UniRef90^53^ sequences that lack any InterPro annotations^3^. (**Fig. S10b**). Candidates are sorted by the TPS prediction scores in descending order, which represent the probability estimates by EnzymeExplorer. Additionally, the number of sequence alignment hits at MARTS-DB^18^ for each candidate is given, as well as the InterPro annotations assigned to them. For sequence alignment, MMSeqs2^46^ is used with 40% minimum coverage and no further limitation, to get as many alignments as possible.

### Structural analysis of the archaeal terpene synthases

All three archaeal terpene synthases are composed of a single α-domain, which are structurally similar to each other (**Fig. S25c**) and to the α1-domains (**Fig. 3a**), however, they do not show any significant similarity to the finer-scale domain subtypes (**Fig. S25a-b**), and sit right at the border of the α1 domain cluster (**Fig. 3a)**. Notably, the α1D is the closest domain subtype to the archaeal α-domains, and these are predominantly found in plant TPSs (**Fig. 1f**).

**Fig. 3.**
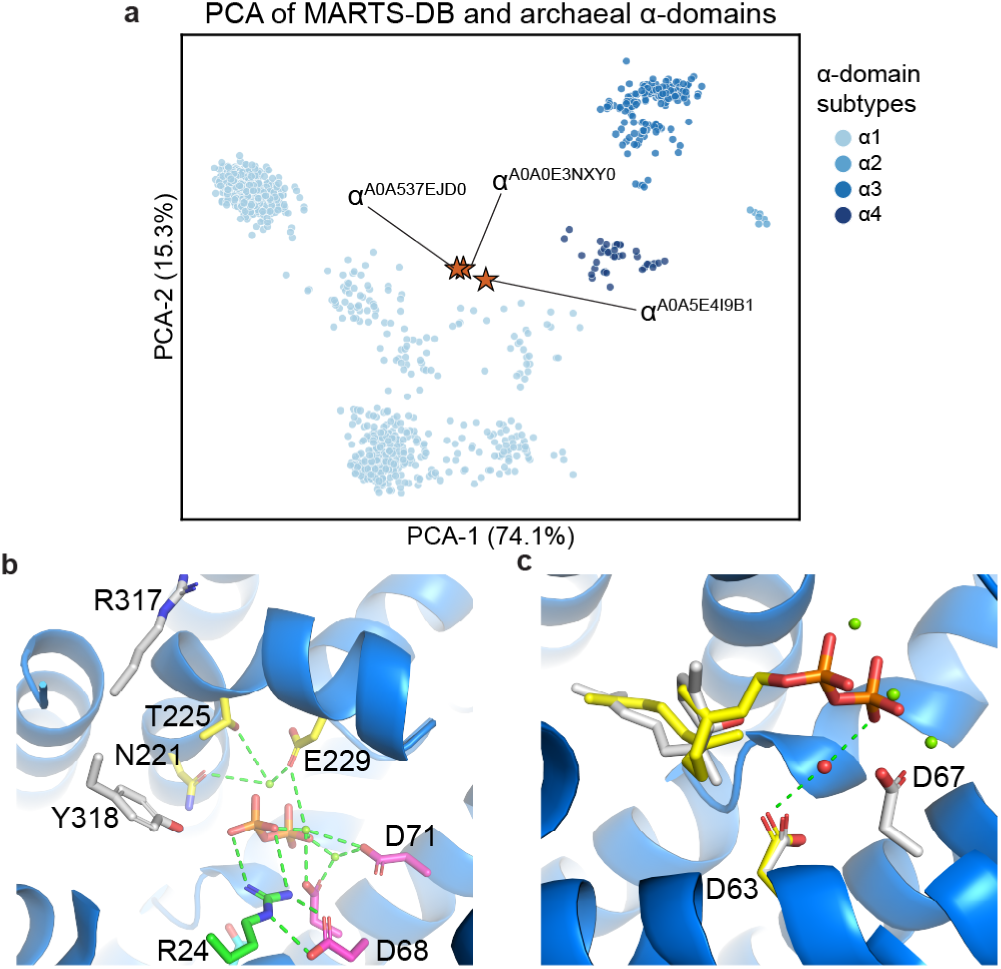
Structural analysis of the archaeal TPSs. **a**, Combined PCA of MARTS-DB α-domains and archaeal α-domains. The archaeal α-domains are illustrated by stars, sitting at the border of the α1 domain subtype in their own small cluster. **b**, Refined TPS model of A0A5E4I9B1. Conserved interactions (≤ 4 Å) marked with green dashed lines are detailed in **Table S2**. **c**, Superposition of the lowest energy pose of the substrate FPP and matching low energy pose of *β*-bisabolol in A0A5E4I9B1 obtained from EnzyDock docking. Position of the catalytic water molecule is shown near the diphosphate moiety.

Although the amino acid sequences of the detected archaeal TPSs have no typical PFAM or SUPFAM domains, we were able to identify several known motifs (**Fig. S20**). Due to the lack of crystal structures, we generated an AlphaFold 3^20^ model of A0A5E4I9B1, including three Mg^2+^ metal ions and docked pyrophosphate in it (**Fig. S21**). The AlphaFold 3 model correctly assigns the ^67^**D**DXX**D** motif in helix D^10^, wherein D67 and D71 coordinate Mg_A_ and Mg_C_ metal ions in bi- and monodentate modes, respectively (bold letters indicate metal binding residues; naming of ions in **Fig. S22**) (**Fig. 3a**). D68 coordinates R24, which is a conserved residue observed in plant TPSs towards the N-terminus sequence region^59^. An additional well-known metal binding motif, which binds Mg_B_ and Mg_C_, is (**N,D**)D(L,I,V)X(**S,T**)XXX**E** in helix H^10^, and in A0A5E4I9B1 this motif is ^221^**N**TLN**T**WPR**E**. Because the conserved TPS catalytic architecture is preserved in the AlphaFold 3 model of A0A5E4I9B1, a more in-depth mechanistic analysis is possible.

The active site contains an unusual D63 which protrudes into the active site (**Fig. 3c**). Considering that this residue is a full helical turn downstream from the highly conserved D67, this is unlikely an artefact of the structure model, and in fact a similar Glu residue is found in 2-Methylisoborneol Synthase^60^ (2-MIBS). A possible role of this Asp residue is to bind and deprotonate an active-site water, which is involved in forming *β*-bisabolol, although the analogous Glu residue in 2-MIBS seemingly does not play a catalytic role^60,61^. To assess the catalytic mechanism in A0A5E4I9B1, we performed reaction modeling. A water molecule possibly binds near the diphosphate moiety, which resembles a conserved water binding site in plant TPS^59^ (**Fig. 3c**). To model the reaction, we performed EnzyDock simulations, which allows rapid multistate docking that can provide mechanistic insight^59,62^. We docked the substrate FPP and various diastereomers of *β*-bisabolol using EnzyDock (**Fig. 3c**). These docking simulations suggest that the isoprenoid chain in FPP connects to the O1*α* oxygen of the diphosphate as observed for plant TPS^59^ and is bound in an extended form (**Table S3**), and the tail of *β*-bisabolol occupies the same binding pocket as in FPP (**Fig. 3c**). In our docking simulations, the aforementioned water molecule is within 4.2 Å of the C6 position in FPP and is close to the position of the hydroxyl group in *β*-bisabolol (4.1-4.5 Å) (**Table S2**).

### Terpenoid biosynthesis is widespread in archaea

The discovery of functional TPSs in archaea marks a significant advancement in our understanding of the distribution and evolutionary origins of TPSs, as these enzymes were thought to exist in archaea, but their discovery remained elusive^55^. To shed more light into this unique finding, we investigated the prevalence of TPSs across the phylogeny of archaea. Analysis of archaeal genomes from the Genome Taxonomy Database^63–65^ (GTDB) revealed TPS presence in 2,867 archaeal taxa at the 95% model-probability threshold and in 1,646 taxa at the 99.95% model-probability threshold (**Supplementary File 6**). Notably, some genomes, such as *Halorussus* caseinilyticus, encode up to five different TPSs. TPSs were detected across 15 of the 25 archaeal phyla at the 99.95% model-probability threshold, with the highest representation in *Halobacteriota*, *Thermoplasmatota*, and *Thermoproteota* (**Fig. 4**).

**Fig. 4.**
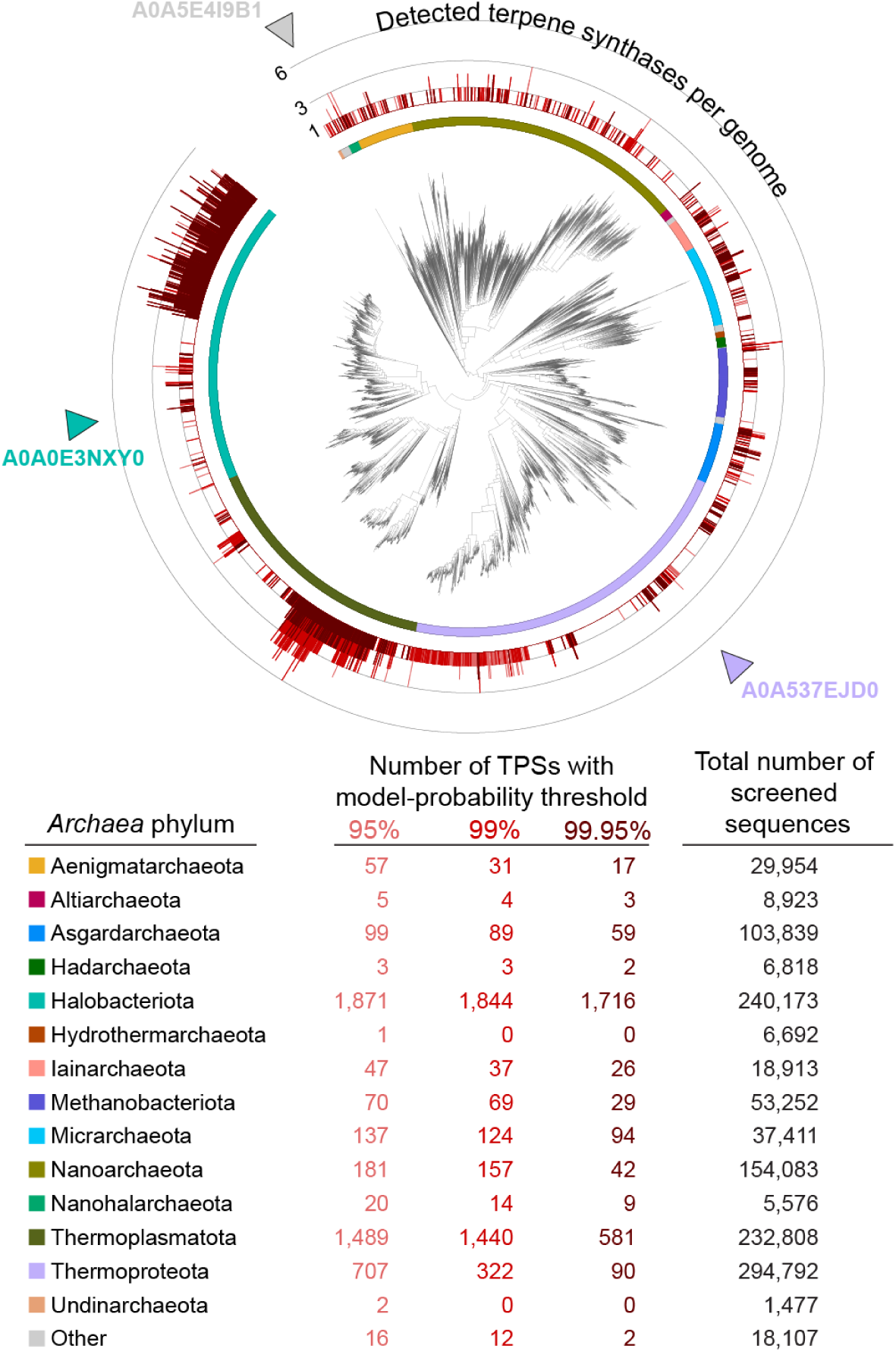
Evolutionary analysis of the putative archaeal TPSs. The Genome Taxonomy Database^65^ phylogenetic tree of archaea with the distribution of predicted terpene synthases depending on the probability threshold of the EnzymeExplorer model. The tree is annotated with the number of predicted TPS detections, based on the selected EnzymeExplorer detection threshold (>95% light red, >99% red, >99.95% dark red; higher value indicates higher precision for TPS activity prediction). The genomes of the experimentally verified archaeal TPSs (A0A0E3NXY0, A0A537EJD0, A0A5E4I9B1) are shown at the phylogenetic tree with triangles.

## Discussion

Compared to state-of-the-art methods for predicting enzyme function^6^, the EnzymeExplorer approach comprises several innovations. By focusing on a single enzyme class, we were able to incorporate class-specific biochemical knowledge on the characteristic structural domains into our machine learning workflow. To achieve that, we developed a structural alignment-based method for precise segmentation of protein structures into structural domains and an unsupervised clustering approach for the identification of subtypes within the established domain types. This annotation of structural domains supports functional predictions, and is therefore important for building trust in the computational annotations, especially in the absence of other recognizable protein signatures. Domain structures tend to be more conserved than sequences, and are linked to enzymatic reaction mechanisms^8,19^.

By combining the structural analyses with a protein language model, we developed a machine-learning approach that significantly outperforms existing methods for enzyme sequence annotation, particularly for sequences from minority classes for which only few examples exist (e.g., sesterTPS). The enhanced performance of our predictive model allowed us to examine the “dark matter” protein sequences that lack any of the ∼85,000 existing InterPro signatures^3^. Existing annotated enzyme datasets are relatively small and sparse, and machine learning methods have not yet demonstrated the capability to discover entirely novel enzymatic sequences within such unexplored regions of protein sequence space. In contrast, EnzymeExplorer successfully predicted seven such enzymes, which we experimentally validated. Furthermore, our automatic annotation of the refined structural domain subtypes provided additional contextual evidence and interpretable family-specific insights for the discoveries.

The same structure-aware pipeline is applicable to the enzyme families whose substrate specificity is encoded in a conserved structural template but remains poorly predictable from sequence alone due to the limitation of standard homology screens below the ∼30% identity twilight zone^66^ that protein language model representations are already known to bypass^21,41,67^. Radical-SAM enzymes are a natural next target^68^ both for prediction of family membership and for the finer question of which C-, N-, S- or O-centred atom the radical is directed against. Modular non-ribosomal peptide synthetases and Type-I polyketide synthases pose the same problem at the domain scale: adenylation and acyltransferase substrate identity remains poorly predictable from sequence signatures alone^69,70^. Iterative Type-II PKS^71^, glycosyltransferases^72^, and enzymatic halogenases^73^ complete a natural target list for structure-aware enzyme function prediction.

The broad generalization capabilities of EnzymeExplorer in annotating the unexplored protein sequence space led to the discovery of the first archaeal TPSs. While it has been well established that cellular membranes of archaea are composed of terpenoid-based phospholipids^55^, and several isoprenyl diphosphate synthase genes have been previously found in archaea^74,75^, no functional terpene cyclization has been observed in archaea until now. This absence of evidence even contributed to the hypothesis that archaea might lack secondary metabolite production^55^. Using NMR spectroscopy, we identified the major product of one archaeal TPS as the sesquiterpene alcohol *β*-bisabolol. *β*-bisabolol has been previously reported in various plant species^76^ and recently the first plant *β*-bisabolol synthase has been discovered^58^. Our method has thus not only uncovered the first non-plant *β*-bisabolol synthase but also demonstrated its potential to discover elusive enzyme sequences from underexplored organisms or metagenomic datasets.

## Acknowledgments

We thank Peter G Mikhael, Itamar Chinn, Chi Zhang, Jitka Štáfková, Petr Kouba, and Regina Barzilay for their constant friendly support and inspiring, insightful discussions about this work.

## Funding

This work was supported by the Ministry of Education, Youth and Sports of the Czech Republic through the e-INFRA CZ (ID:90140). T.H. was supported by the European Union’s Horizon Europe research and innovation program under the Marie Skłodowska-Curie grant agreement No. 101130799 and the IOCB Postdoctoral Fellowship program. T.P. was supported by the Czech Science Foundation (GA CR) grant 21-11563M and by the European Union’s Horizon Europe program (ERC, TerpenCode, 101170268 and Marie Skłodowska-Curie Actions, ModBioTerp, 101168583). J.S. was supported by the European Union’s Horizon Europe program (ERC, FRONTIER, 101097822 and ELIAS, 101120237). Views and opinions expressed are, however, those of the authors only and do not necessarily reflect those of the European Union or the European Research Council. Neither the European Union nor the granting authority can be held responsible for them.

## Author contributions

T.P. conceptualized the project. R.Sam. conducted the initial computational research phase, which included exploratory data analysis; development of the initial structural-domain segmentation, clustering, and predictive-modeling workflows; biological interpretation of the domain analyses; screening of UniRef90 and UniProt; automated mining of characterized TPSs; and preparation of the original manuscript draft. Following the initial review, S.M.A. substantially revised, extended, and reworked the computational analyses, codebase, results, and manuscript, with contributions from R.Sam. and input from all authors. T.H. performed metabolic engineering, LC-MS, and GC-MS analyses; processed laboratory experimental data and wrote corresponding parts of the manuscript. T.H. and J.D.S. conducted compound purification and NMR analyses. T.H. and A.S performed the protein expression and *in vitro* assays. S.C.K. wrote the manuscript and provided critical feedback to the project. T.Č. mined putative TPSs for model fine-tuning. H.S. and M.M. performed metabolic engineering. R.B., A.B., and J.K. performed fine-tuning of protein language models. R.Ch. performed error analysis. R.Ch., R.B., and A.B. performed exploratory data analysis. D.T.M and R.Sch. performed TPS structural model refinement and docking. J.S. and T.P. supervised the project.

## Code and data availability

The source code developed for this study is available on GitHub under the MIT license (https://github.com/pluskal-lab/EnzymeExplorer).

The predictions of EnzymeExplorer for UniRef90 clusters with sequences longer than 230 amino acid residues and without InterPro annotations are available on Zenodo under CC by 4.0 (DOI: 10.5281/zenodo.21559271).

## Notes

https://doi.org/10.5281/ZENODO.10359045

### Competing Interest Statement

The authors have declared no competing interest.

### Summary of Updates

We substantially improved both the EnzymeExplorer pipeline and the manuscript. The revised version provides greater clarity, rigor, and transparency, particularly with respect to the computational validation of the method. We have also strengthened the experimental validation of the selected enzyme sequences through in vitro biochemical assays.

## References

1. Jones, P. et al. InterProScan 5: genome-scale protein function classification. Bioinformatics 30, 1236–1240 (2014).

2. UniProt Consortium. UniProt: The universal protein knowledgebase in 2025. Nucleic Acids Res. 53, D609–D617 (2025).

3. Blum, M. et al. InterPro: the protein sequence classification resource in 2025. Nucleic Acids Res. 53, D444–D456 (2025).

4. Buller, R. et al. From nature to industry: Harnessing enzymes for biocatalysis. Science 382, eadh8615 (2023).

5. Kim, G. B. et al. Functional annotation of enzyme-encoding genes using deep learning with transformer layers. Nat. Commun. 14, 7370 (2023).

6. Yu, T. et al. Enzyme function prediction using contrastive learning. Science 379, 1358–1363 (2023).

7. de Crécy-Lagard, V., Dias, R., Friedberg, I., Yuan, Y. & Swairjo, M. A. Limitations of current machine-learning models in predicting enzymatic functions for uncharacterized proteins. bioRxiv (2024) doi:10.1101/2024.07.01.601547.

8. Durairaj, J. et al. Integrating structure-based machine learning and co-evolution to investigate specificity in plant sesquiterpene synthases. PLoS Comput. Biol. 17, e1008197 (2021).

9. Gao, Y., Honzatko, R. B. & Peters, R. J. Terpenoid synthase structures: a so far incomplete view of complex catalysis. Nat. Prod. Rep. 29, 1153–1175 (2012).

10. Christianson, D. W. Structural and Chemical Biology of Terpenoid Cyclases. Chem. Rev. 117, 11570–11648 (2017).

11. White, N. J. Qinghaosu (artemisinin): the price of success. Science 320, 330–334 (2008).

12. Köksal, M., Jin, Y., Coates, R. M., Croteau, R. & Christianson, D. W. Taxadiene synthase structure and evolution of modular architecture in terpene biosynthesis. Nature 469, 116–120 (2010).

13. Raz, K., Levi, S., Gupta, P. K. & Major, D. T. Enzymatic control of product distribution in terpene synthases: insights from multiscale simulations. Curr. Opin. Biotechnol. 65, 248–258 (2020).

14. Yoshikuni, Y., Ferrin, T. E. & Keasling, J. D. Designed divergent evolution of enzyme function. Nature 440, 1078–1082 (2006).

15. Greenhagen, B. T., O’Maille, P. E., Noel, J. P. & Chappell, J. Identifying and manipulating structural determinates linking catalytic specificities in terpene synthases. Proc. Natl. Acad. Sci. U. S. A. 103, 9826–9831 (2006).

16. Kampranis, S. C. et al. Rational conversion of substrate and product specificity in a Salvia monoterpene synthase: structural insights into the evolution of terpene synthase function. Plant Cell 19, 1994–2005 (2007).

17. Whitehead, J. N., Leferink, N. G. H., Johannissen, L. O., Hay, S. & Scrutton, N. S. Decoding Catalysis by Terpene Synthases. ACS Catal. 13, 12774–12802 (2023).

18. Engst, M., et al. MARTS-DB: a database of mechanisms and reactions of terpene synthases. BMC Bioinformatics 27, 10 (2025).

19. Holliday, G. L., Fischer, J. D., Mitchell, J. B. O. & Thornton, J. M. Characterizing the complexity of enzymes on the basis of their mechanisms and structures with a bio-computational analysis: The complexity of enzymes. FEBS J. 278, 3835–3845 (2011).

20. Abramson, J. et al. Accurate structure prediction of biomolecular interactions with AlphaFold 3. Nature 630, 493–500 (2024).

21. Lin, Z. et al. Evolutionary-scale prediction of atomic-level protein structure with a language model. Science 379, 1123–1130 (2023).

22. Murtagh, F. & Contreras, P. Algorithms for hierarchical clustering: an overview. Wiley Interdiscip. Rev. Data Min. Knowl. Discov. 2, 86–97 (2012).

23. Loewenstein, Y., Portugaly, E., Fromer, M. & Linial, M. Efficient algorithms for accurate hierarchical clustering of huge datasets: tackling the entire protein space. Bioinformatics 24, i41–9 (2008).

24. Kumar, S., Stecher, G., Li, M., Knyaz, C. & Tamura, K. MEGA X: Molecular Evolutionary Genetics Analysis across computing platforms. Mol. Biol. Evol. 35, 1547–1549 (2018).

25. Katoh, K., Rozewicki, J. & Yamada, K. D. MAFFT online service: multiple sequence alignment, interactive sequence choice and visualization. Brief. Bioinform. 20, 1160–1166 (2019).

26. Kaplan, N., Friedlich, M., Fromer, M. & Linial, M. A functional hierarchical organization of the protein sequence space. BMC Bioinformatics 5, 196 (2004).

27. Mi, H., Guo, N., Kejariwal, A. & Thomas, P. D. PANTHER version 6: protein sequence and function evolution data with expanded representation of biological pathways. Nucleic Acids Res. 35, D247–52 (2007).

28. Frech, C. & Chen, N. Genome-wide comparative gene family classification. PLoS One 5, e13409 (2010).

29. Ma, Q., Chirn, G.-W., Cai, R., Szustakowski, J. D. & Nirmala, N. R. Clustering protein sequences with a novel metric transformed from sequence similarity scores and sequence alignments with neural networks. BMC Bioinformatics 6, 242 (2005).

30. Langfelder, P., Zhang, B. & Horvath, S. Defining clusters from a hierarchical cluster tree: the Dynamic Tree Cut package for R. Bioinformatics 24, 719–720 (2008).

31. Verdoes, J. C., Krubasik, K. P., Sandmann, G. & van Ooyen, A. J. Isolation and functional characterisation of a novel type of carotenoid biosynthetic gene from Xanthophyllomyces dendrorhous. Mol. Gen. Genet. 262, 453–461 (1999).

32. Sandmann, G., Zhu, C., Krubasik, P. & Fraser, P. D. The biotechnological potential of the al-2 gene from Neurospora [correction of Neurospra] crassa for the production of monocyclic keto hydroxy carotenoids. Biochim. Biophys. Acta 1761, 1085–1092 (2006).

33. Knudsen, M. & Wiuf, C. The CATH database. Hum. Genomics 4, 207–212 (2010).

34. Lesburg, C. A. & Christianson, D. W. PENTALENENE SYNTHASE. Preprint at 10.2210/pdb1ps1/pdb (1998).

35. Tarshis, L. C., Proteau, P., Poulter, C. D. & Sacchettini, J. C. STRUCTURE OF FARNESYL PYROPHOSPHATE SYNTHETASE. Preprint at 10.2210/pdb1ubw/pdb (1997).

36. Starks, C. M., Back, K., Chappell, J. & Noel, J. P. 5-EPI-ARISTOLOCHENE SYNTHASE FROM NICOTIANA TABACUM WITH SUBSTRATE ANALOG FARNESYL HYDROXYPHOSPHONATE. Preprint at 10.2210/pdb5eat/pdb (1997).

37. Koksal, M. & Christianson, D. W. Crystal Structure of Taxadiene Synthase from Pacific Yew (Taxus brevifolia) in complex with Mg2+ and 2-fluorogeranylgeranyl diphosphate. Preprint at 10.2210/pdb3p5r/pdb (2010).

38. Thoma, R. et al. Structure of human OSC in complex with Ro 48-8071. Preprint at 10.2210/pdb1w6j/pdb (2004).

39. Berman, H. M. et al. The Protein Data Bank. Nucleic Acids Res. 28, 235–242 (2000).

40. Barrio-Hernandez, I. et al. Clustering predicted structures at the scale of the known protein universe. Nature (2023) doi:10.1038/s41586-023-06510-w.

41. Elnaggar, A., et al. Ankh: Optimized protein language model unlocks general-purpose modelling. *arXiv [cs.LG]* (2023) doi:10.48550/arXiv.2301.06568.

42. Bepler, T. & Berger, B. Learning the protein language: Evolution, structure, and function. Cell Syst. 12, 654–669.e3 (2021).

43. Tule, S., Foley, G. & Bodén, M. Do protein language models learn phylogeny? Brief. Bioinform. 26, bbaf047 (2024).

44. Breiman, L. Random Forests. Mach. Learn. 45, 5–32 (2001).

45. Boeckmann, B. et al. The SWISS-PROT protein knowledgebase and its supplement TrEMBL in 2003. Nucleic Acids Res. 31, 365–370 (2003).

46. Steinegger, M. & Söding, J. MMseqs2 enables sensitive protein sequence searching for the analysis of massive data sets. Nat. Biotechnol. 35, 1026–1028 (2017).

47. Bateman, A. et al. The Pfam protein families database. Nucleic Acids Res. 32, D138–41 (2004).

48. Pandit, S. B. et al. SUPFAM: a database of sequence superfamilies of protein domains. BMC Bioinformatics 5, 28 (2004).

49. Priya, P., Yadav, A., Chand, J. & Yadav, G. Terzyme: a tool for identification and analysis of the plant terpenome. Plant Methods 14, 4 (2018).

50. Altschul, S. F. et al. Gapped BLAST and PSI-BLAST: a new generation of protein database search programs. Nucleic Acids Res. 25, 3389–3402 (1997).

51. Efron, B. Better bootstrap confidence intervals. J. Am. Stat. Assoc. 82, 171 (1987).

52. Wong, J. et al. High-titer production of lathyrane diterpenoids from sugar by engineered Saccharomyces cerevisiae. Metab. Eng. 45, 142–148 (2018).

53. Suzek, B. E. et al. UniRef clusters: a comprehensive and scalable alternative for improving sequence similarity searches. Bioinformatics 31, 926–932 (2015).

54. Varadi, M. et al. AlphaFold Protein Structure Database: massively expanding the structural coverage of protein-sequence space with high-accuracy models. Nucleic Acids Res. 50, D439–D444 (2022).

55. Hoshino, Y. & Villanueva, L. Four billion years of microbial terpenome evolution. FEMS Microbiol. Rev. 47, (2023).

56. Vattekkatte, A., Garms, S., Brandt, W. & Boland, W. Enhanced structural diversity in terpenoid biosynthesis: enzymes, substrates and cofactors. Org. Biomol. Chem. 16, 348–362 (2018).

57. Steele, C. L., Crock, J., Bohlmann, J. & Croteau, R. Sesquiterpene synthases from grand fir (Abies grandis). Comparison of constitutive and wound-induced activities, and cDNA isolation, characterization, and bacterial expression of delta-selinene synthase and gamma-humulene synthase. J. Biol. Chem. 273, 2078–2089 (1998).

58. Pu, W.-T. et al. Functional characterization of a sesquiterpene synthase involved in stereospecific biosynthesis of β-bisabolol. J. Agric. Food Chem. 73, 32775–32785 (2025).

59. Schwartz, R., Zev, S. & Major, D. T. Differential substrate sensing in terpene synthases from plants and microorganisms: Insight from structural, bioinformatic, and EnzyDock analyses. Angew. Chem. Int. Ed Engl. 63, e202400743 (2024).

60. Köksal, M., Chou, W. K. W., Cane, D. E. & Christianson, D. W. Structure of 2-methylisoborneol synthase from Streptomyces coelicolor and implications for the cyclization of a noncanonical C-methylated monoterpenoid substrate. Biochemistry 51, 3011–3020 (2012).

61. Gu, B. & Dickschat, J. S. Functional characterisation of highly conserved and structurally prominent residues of 2-methylisoborneol synthase. Chem. Eur. J. 29, e202300775 (2023).

62. Das, S. et al. EnzyDock: Protein-ligand docking of multiple reactive states along a reaction coordinate in enzymes. J. Chem. Theory Comput. 15, 5116–5134 (2019).

63. Parks, D. H. et al. GTDB: an ongoing census of bacterial and archaeal diversity through a phylogenetically consistent, rank normalized and complete genome-based taxonomy. Nucleic Acids Res. 50, D785–D794 (2022).

64. Parks, D. H. et al. GTDB release 10: a complete and systematic taxonomy for 715 230 bacterial and 17 245 archaeal genomes. Nucleic Acids Res. 54, D743–D754 (2026).

65. Rinke, C. et al. A standardized archaeal taxonomy for the Genome Taxonomy Database. Nat. Microbiol. 6, 946–959 (2021).

66. Rost, B. Twilight zone of protein sequence alignments. Protein Eng. 12, 85–94 (1999).

67. Rives, A. et al. Biological structure and function emerge from scaling unsupervised learning to 250 million protein sequences. Proc. Natl. Acad. Sci. U. S. A. 118, (2021).

68. Broderick, J. B., Duffus, B. R., Duschene, K. S. & Shepard, E. M. Radical S-adenosylmethionine enzymes. Chem. Rev. 114, 4229–4317 (2014).

69. Süssmuth, R. D. & Mainz, A. Nonribosomal peptide synthesis-principles and prospects. Angew. Chem. Int. Ed Engl. 56, 3770–3821 (2017).

70. Khosla, C., Tang, Y., Chen, A. Y., Schnarr, N. A. & Cane, D. E. Structure and mechanism of the 6-deoxyerythronolide B synthase. Annu. Rev. Biochem. 76, 195–221 (2007).

71. Hertweck, C., Luzhetskyy, A., Rebets, Y. & Bechthold, A. Type II polyketide synthases: gaining a deeper insight into enzymatic teamwork. Nat. Prod. Rep. 24, 162–190 (2007).

72. Drula, E. et al. The carbohydrate-active enzyme database: functions and literature. Nucleic Acids Res. 50, D571–D577 (2022).

73. Andorfer, M. C. & Lewis, J. C. Understanding and improving the activity of flavin-dependent halogenases via random and targeted Mutagenesis. Annu. Rev. Biochem. 87, 159–185 (2018).

74. Tachibana, A. et al. Novel prenyltransferase gene encoding farnesylgeranyl diphosphate synthase from a hyperthermophilic archaeon, Aeropyrum pernix. Molecularevolution with alteration in product specificity. Eur. J. Biochem. 267, 321–328 (2000).

75. Petrova, T. E. et al. Structural characterization of geranylgeranyl pyrophosphate synthase GACE1337 from the hyperthermophilic archaeon Geoglobus acetivorans. Extremophiles 22, 877–888 (2018).

76. Egbuta, M. A., McIntosh, S., Waters, D. L. E., Vancov, T. & Liu, L. In vitro anti-inflammatory activity of essential oil and β-bisabolol derived from Cotton Gin trash. Molecules 27, 526 (2022).

