## Supplementary Information for "Structure-enabled enzyme function prediction unveils elusive terpenoid biosynthesis in archaea"

### Materials and Methods

#### Dataset construction

MARTS-DB[18] is a manually curated database of experimentally verified TPSs, and some additional IDSs as well. It provides the data for the sequence and the reactions catalyzed by these enzymes. We used the snapshot version of MARTS-DB from the 22nd of February 2026 as the initial raw data. After a deduplication of sequences, we finalized a dataset with 1,374 sequences (1,256 TPSs and 181 IDSs, with 63 bifunctional TPS&IDSs), and 4,185 reactions (**Supplementary File 1**). We applied AlphaFold 3[20] and ESMFold[21] to receive the structural predictions of each sequence and picked the structure with highest pLDDT score for each sequence. The final dataset of sequences and structures are then used for the structural domain segmentation and annotation.

For the predictive pipeline, we enriched the dataset with 10,000 additional negatives (enzymes that do not catalyze TPS reactions). These negatives were taken from Swiss-Prot[45] via separate pipelines for hard negatives (non-TPS enzymes that are sequentially/structurally/biochemically similar to TPSs) and easy negatives (non-TPS enzymes that have no similarity to TPSs), predicted 3D structures of which were already provided in AlphaFold Protein Structure Database[54] (AFDB). We preprocessed Swiss-Prot[45] (Uniprot version 2026_02) by removing all putative TPSs by RHEA IDs[77] (RHEA Knowledgebase release 140), EC numbers[77] (RHEA Knowledgebase release 140), GO annotations[78,79] (release 2026-01-23), and Pfam & SUPFAM annotations[47,48]. We then reduced the redundancy across the remaining data by 95% sequence identity using MMSeqs2[46]. The first batch of hard negatives were fetched using RHEA knowledgebase (RHEA Knowledgebase release 140) to mine for the enzymes that accept the TPS substrates found in MARTS-DB. We used MMSeqs2 to fetch the second batch of hard negatives that are sequentially similar to the MARTS-DB sequences. Combining these two batches of hard negatives resulted in 5,353 negative samples. We topped these off to 10,000 negatives by adding the easy negatives from the rest of the negative data. To this end, we clustered these negatives by 50% sequence identity using MMSeqs2, and randomly picked 4,647 cluster representatives as easy negatives.

To gather a dataset of putative TPS-like sequences, we mined large-scale protein sequence resources, including BFD[80], UniParc[81], Mgnify[82], 1KP[83], Phytozome[84] and NCBI Transcriptome Shotgun Assembly (TSA) databases. Protein sequences were selected based on matches to the same TPS-specific Pfam and SUPFAM domain combinations. This mining effort resulted in approximately 80 thousand TPS-like sequences (**Supplementary File 3**).

#### Structural domain segmentation and annotation

We implemented a two-step methodology to segment predicted protein structures into enzymatic domains and to annotate them with refined structural subtypes. The first step employs an alignment-based approach to partition predicted structures into family-specific enzymatic domains using established structural templates (**Fig. S3**). Next, an unsupervised clustering method based on structural similarity metrics (TM-score) identifies subtypes within these domains, leveraging characterized enzymes as reference examples.

**Structural domain segmentation**

We used structural domains of well-characterized, canonical TPSs from prior studies[10] as templates for TPS-specific domain definitions. These reference enzymes were chosen because they have high-resolution crystal structures with established domain boundaries and catalytic motifs, they span the major TPS architectures relevant to our targets (e.g. single-domain class I α; plant αβγ; sterol-type βγ) as well as the IDS architecture (single-domain IDS α), and they are widely cited standards in the TPS structural literature, ensuring reproducibility against community benchmarks. Specifically, the fold of pentalenene synthase[85] served as the template for the α domain because it is a prototypical class I bacterial sesquiterpene cyclase with clearly resolved **DD**xx**D** and **NSE**/**DTE** metal-binding motifs. In addition to this, we used a farnesyl pyrophosphate synthase[86] as the secondary template for the α domain since it is a very well established α domain with IDS-like fold[10]. To define the β domain, we aligned the α-domain template to the characterized structure of 5-epi-aristolochene synthase[87], a canonical plant TPS, and used the unaligned remainder as the β-domain standard, leveraging its well-annotated αβ architecture. Similarly, after aligning both α and β templates to taxadiene synthase[12], a representative αβγ diterpene cyclase, the γ domain standard was defined as the remaining unaligned region. In addition, domains from lanosterol synthase with βγ architecture[88] were included as alternative β and γ templates to capture sterol-type β/γ folds that are structurally related to plant TPS β/γ domains. This template panel thus anchors our segmentation in diverse TPS examples that collectively cover the domain organizations encountered in TPS families. After the initial run of domain segmentation, we encountered a novel domain structure in two sequences (UniProt ID Q7Z859 and P37295) with a very poor alignment score to these established domain types. This led to the definition of an extra domain template denoted with ζ from the AlphaFold 3[20] predicted structure of P37295. Moreover, due to structural uniqueness of the alternative β and γ templates, which was shown by the domain subtype identification **(Fig S4)**, they are renamed as δ and ε domains, respectively. All TPS-specific structural domain templates are given in **Fig 1b**.

To segment predicted structures we aligned them to a curated panel of seven family-specific templates using sequence-independent alignment tools PyMOL v3.1.6[89] and US-align v20241108[90]. Templates are pre-filtered to their canonical secondary-structure residues to remove loop-driven noise[91]. Each template carries its own detection cutoffs, tuned by inspection of positive and negative controls: the required TM-score ranges from 0.35 for the α template to 0.65 for the more compact ε and ζ folds, and the minimum number of aligned residues from 60 to 100. The US-align algorithm outputs partial residue mappings between the template and query structure. Hence, we developed an approach (applied as a post-processing step) to pair all residues in the domain template to corresponding residues in the query structure. For each unmapped residue in the template, the nearest mapped residue pair was used to derive local sequence shifts, extending mappings to include all residues in the template. Detections are then confidence-filtered against the per-residue pLDDT, keeping residues with pLDDT ≥ 70. Segmentation proceeds iteratively for multiple rounds until no alignment for the templates are found. Within a round we align every template against every query, keep the detection that meets both the TM-score and length thresholds with the highest TM-score, and resolve competing detections on the same structure with a disjoint-domain selector based on TM-score of each detection. Residues newly claimed by the accepted detection are removed from the query's residue pool, and the next round rescans the remainder. Between rounds, unassigned helices and sheets close to a detected domain are attached to that domain.

**Domain subtype identification**

After segmenting all protein structures in MARTS-DB into domains, we performed pairwise comparisons of the detected domains using US-align[90] to compute TM-scores to measure structural similarity. An unweighted pair group method with arithmetic mean (UPGMA) was then employed to organize the domains into hierarchical clusters and to build a dendrogram of structural domains **(Fig. S4a)**. UPGMA is also referred to as average-linkage hierarchical agglomerative clustering, an algorithm widely used in bioinformatics for phylogenetics[24,25] and biological classification analyses[26–29]. SciPy[92] (v1.13.0) Python library was used for the implementation of UPGMA. Because the UPGMA dendrogram exhibited heterogeneous branch heights and nested substructures, applying a single global distance threshold failed to consistently isolate structurally coherent groups. Therefore, we identified subtype boundaries using the Dynamic Tree Cut[30] algorithm, which adaptively partitions UPGMA dendrograms by leveraging local branch topology rather than a fixed cut height. This approach allows clusters of varying sizes and densities to be detected while preserving nested hierarchical organization. To objectively determine the optimal hyperparameters (minClusterSize=3 and deepSplit=0), we conducted a systematic combinatorial sweep. These final parameters were selected because they maximized cluster reproducibility and compactness while minimizing the fraction of unstable, transitional domains across the resulting subtypes **(Fig. S5)**. The clades determined by Dynamic Tree Cut were defined as domain subtypes, and organized manually based on the dendrogram hierarchy. The δ3 and ζ domains were initially not detected by Dynamic Tree Cut due to the low number of members, and were assigned manually as a postprocessing step. Finally, we analysed each domain subtype cluster w.r.t. their reproducibility, stability, and compactness **(Figs. S4c-e, S24)**.

### Protein language model fine-tuning

Using the mined TPS-like sequences (**Supplementary File 3**, see **Dataset construction**), we performed an end-to-end fine-tuning of the ESM-1b model[67], where the original checkpoint (ESM-1v) was trained on the UniRef90 dataset[53]. We adopted the original BERT-like masking strategy and selected 15% of the input protein sequence tokens for masking. Of these tokens, 80% were replaced with the <mask> token, 10% were replaced with a random token, and the remaining 10% were left unchanged. Unlike conventional token-wise masking, we applied masking to contiguous subsequences of randomly sampled lengths ranging from 2 to 6 tokens. We limited the input length to 1024 tokens, and for sequences exceeding this length, we randomly sampled a subsequence of the desired length. The model was trained for 100,000 steps with Adam optimizer with inverse square root learning rate scheduler, with an initial learning rate of 1e-4 and 16,000 warm-up steps.

Using the same TPS-like sequences, we performed an end-to-end fine-tuning of the Ankh-base[41] model, an encoder-decoder (T5-based[93]) protein language model originally pre-trained on UniRef50[53]. Consistent with Ankh's original pre-training objective, we adopted its span-denoising masking strategy rather than the BERT-like scheme used for ESM-1v: 20% of the input protein sequence tokens were randomly selected and each masked token was replaced with a unique sentinel token, while the decoder was trained to reconstruct the corresponding original residues from the sentinel-delimited target sequence. We limited the input length to 512 tokens. The model was fine-tuned for 30 epochs (approximately 67,000 steps) with a global batch size of 32 using the AdamW optimizer with a linearly decaying learning rate scheduler, an initial learning rate of 5e-5, and no warm-up steps. 10% of the sequences were held out for validation.

We used both of the fine-tuned models in an ablation study of different PLMs **(Fig. S9)**.

#### Predictive modeling

We extracted numerical representations of input sequences from the Ankh-large protein language model (PLM) and the predicted structure. Predicted protein structures were segmented into family-specific domains (see **Structural Domain Segmentation and Annotation**). We computed each detected domain's structural similarity (TM-score) against all known domains of the same type in the training dataset using Foldseek[94]. The TM-scores served as a structural similarity profile for each domain. We concatenated the structural profile with PLM embeddings to obtain an input vector for predictive models. Using this input vector, we trained a multi-label random forest classifier to detect family-specific enzymatic activity and predict substrates. For the case of TPSs, we picked eight canonical substrates which are the most common ones found in nature and supported by sufficient amount of data in the dataset for a 5-fold cross-validation split with 50% maximum sequence identity threshold (GPP, FPP, GGPP, GFPP, copalyl-PP, 2,3-epoxysqualene, 2xFPP, 2xGGPP). The stereochemistry was ignored while assigning substrate classes to the TPSs in MARTS-DB. The pipeline was implemented in Python 3.10.0. We used the random forest[44] implementation from the scikit-learn library (v1.7.2). Feature extraction based on PLM representations was implemented using PyTorch.

Hyperparameter tuning was performed on training data via nested-cross-validation within the training folds. Additionally, we implemented a hyperparameter tuning approach to optimize model performance. Hyperparameter optimization was conducted using the scikit-optimize library (v0.10.2), leveraging Gaussian Processes (gp_minimize function).

#### Performance evaluation of predictive models

We evaluated the performance of predictive models using the final dataset (**Supplementary File 4**) described in **Dataset construction** section**.** For each evaluated model, we conducted multiple experiments where part of the data was held out as a test set. We employed 5-fold cross-validation, in which one fold served as a hold-out dataset, untouched during model training and hyperparameter tuning, while the remaining four folds were used for training.

##### Stratified Group K-Fold Splitting

To ensure that models generalize to proteins distant from the training set, we used a stratified group k-fold. Groups were defined based on an MMSeqs2[46] sequence clustering with 50% maximum sequence identity and 80% minimum coverage.

The default implementation of stratified group k-fold in scikit-learn produced folds with unstable class proportions. To address this, we iterated over random seeds and selected the fold split that minimized the Jensen-Shannon divergence between the class distributions in each fold and the overall dataset. This ensured more balanced class distributions across folds.

**Evaluation Metrics**

Model performance was evaluated using average precision (mAP) as the primary metric. AP summarizes the performance of score-based classifiers across different detection thresholds and is well-suited for imbalanced datasets[95]. Additionally, we computed the area under the ROC curve (AUC-ROC) as a secondary evaluation metric **(Fig. S7g-h)**. For substrate prediction metrics, we used macro-averaging due to the dataset imbalance (mean AP).

**Uncertainty Quantification and Statistical Significance**

For every classifier and every target class we produced out-of-fold (OOF) predictions by 5-fold cross-validation and concatenated the predictions from the five held-out folds into a single evaluation pool. The point estimates of each performance metrics (AP, mean AP, and AUC-ROC) were computed on this original, un-resampled pool.

To quantify uncertainty we performed block-bootstrapping. To this end, we clustered the dataset with 50% sequence similarity and 80% coverage (same as **Stratified Group K-Fold Splitting**), and assigned a group id to each sequence based on their cluster. We then drew n = 1,000 bootstrap resamples of the pooled OOF groups with replacement with the same number of groups as the original pool. A single set of resampled group indices was shared across every method and every class within a draw. Resamples in which any requested class had zero positives or zero negatives were discarded and redrawn; this kept every metric defined without biasing the sampling distribution. The metric was recomputed on each resample, yielding a 1,000-point bootstrap distribution per (method, class, metric). Two-sided 95% confidence intervals (CI) on each (method, class, metric) were computed with the bias-corrected accelerated bootstrap (BCa) method[51] (**Figs. 2b-e, S7c-d, S7g-h, S8, S9**). Because the resampled group indices were shared across methods within a draw, the bootstrap was a proper paired design at the group level. For every pair of methods and every draw we recorded the per-draw difference (e.g. $AP_{EnzymeExplorer}$ - $AP_{Foldseek}$). The paired construction removed shared sampling variance between A and B and produced tighter, less noisy delta distributions than would have been obtained from two independent bootstraps, which were then shown with forest plots (**Figs. S7c,g, S8a,b,d,e, S9a,b**). Each "delta" forest plot showed one 95% CI box per method pair, with the box spanning the inner 25–75% percentile range of the 1,000 paired Δ values, the whiskers extended to the BCa CI limits, and the median as a horizontal line inside the box. A box lying entirely above zero indicates that method A outperforms method B at the 95% confidence level in the paired sense; a box that overlaps zero indicates the pairwise difference is not significant at that level. Using these analyses, we were able to show that EnzymeExplorer outperforms all baselines with statistical significance (**Figs**. **2b,d, S7c**)

**Post-hoc Transformation of Random Forest Prediction Scores for Calibration**

The raw scores produced by EnzymeExplorer for each prediction class were calibrated by a post-hoc transformation into probability estimates without retraining the base model. We used temperature scaling[96] for all prediction tasks except for two substrate classes (GPP and FPP) where a logit-input Platt rescaler[97] was used due to better data availability and lower risk of overfitting (**Table S5**). To this end, LogisticRegression from scikit-learn (v1.5.1) was utilized. For the deployment, we fitted one global calibration per each task using the pooled out-of-fold predictions, which produces the final calibrated probabilities of the model. The calibration procedure did not have any effect on the model training or the ranking of predictions. It was implemented to obtain interpretable prediction scores from the deployment pipeline for downstream candidate selection. Even though the calibration was applied for each prediction task separately, only the calibrated TPS prediction probabilities were used in candidate selection procedures. This was because the improvements on log-loss and Brier scores after calibration were relatively higher for the TPS detection task (**Table S8**). We used 95% calibrated TPS prediction probability as the lowest model-probability threshold as at this threshold, the classifier achieves precision 0.999 and recall 0.987 on the held-out folds (**Table S7**). For the higher model-probability thresholds (used in section **Terpenoid biosynthesis is widespread in archaea**), 99% and 99.95% are selected based on the predicted probabilities of the experimentally validated archaeal candidates (**Table 1**).

#### Baselines

We compared the performance of our approach against several well-established baseline models: Pfam[47] and SUPFAM[48] domain-based annotation tools for identifying conserved protein families and functional domains, a state-of-the-art deep-learning-based tool Foldseek[94] designed for protein search within the structure space, a state-of-the-art deep-learning model CLEAN[6] for enzyme annotation based on Enzyme Commission (EC) number prediction, profile Hidden Markov Models (pHMMs) trained using the HMMER[98] library on our curated dataset following the procedure outlined in the Terzyme paper[49], a sequence homology search tool BLASTp[50]. We used official implementations for all baselines.

The alignment-based methods (such as Pfam, SUPFAM, Foldseek, BLAST, and pHMM) were evaluated based on calculation of a prediction score using the number of correct hits over all significant alignments. To this end, for each method we determined the best performing E-value and bit-score thresholds regarding each prediction task (independently for TPS detection and each substrate class). For example, TPS detection for BLAST is evaluated by querying each cross-validation test fold to the corresponding train fold with the E-value threshold $10^{-5}$ (which was determined by sweeping multiple E-value thresholds from 1 to $10^{-200}$), and yielding the rate of TPSs over all hits as the prediction score. Even though this maximized the performance of each baseline method unfairly to EnzymeExplorer, we were still able to show that EnzymeExplorer outperformed all baseline methods. Moreover, we retrained CLEAN by enriching its dataset with the TPSs in MARTS-DB, so that a fair comparison can be made between the models. This retraining improved CLEAN's performance significantly, nevertheless, it was outperformed by Pfam, BLAST, Foldseek, and EnzymeExplorer (**Figs. 2b-e**). The prediction scores of CLEAN were calculated via translating the predicted EC numbers to RHEA[77] reactions. The ratio of TPS reactions among the predicted EC numbers was returned as the TPS detection score, and for the substrate prediction tasks, the ratio of TPS reactions accepting the corresponding substrate was returned as the substrate prediction score, analogously.

### Ablation study

To evaluate the choice of PLMs, contribution of input features, and classifier algorithms to the overall predictive performance, we conducted three ablation studies.

We evaluated six different PLM models in TPS detection and substrate prediction tasks to choose the best model for our pipeline: Ankh-base[41], Ankh-large[41], ESM-1v[99], ESM-2[21] (esm2_t36_3B_UR50D), fine-tuned Ankh-base (see **Protein language model fine-tuning)**, and fine-tuned ESM-1v (see **Protein language model fine-tuning)**. The evaluations showed that both ESM-2 and Ankh-large models were standing out for TPS detection, but for substrate prediction, Ankh-large performs slightly better than the rest (**Fig. S9**). Surprisingly, the PLM ablation showed that fine-tuning does not improve the performance of the EnzymeExplorer.

We also assessed the role of protein language model (PLM) embeddings, structural domain features, and their combination in predicting TPS functions and substrate specificities. Models were trained using either PLM embeddings only, or structural domain features only, or a concatenation of PLM embeddings and structural features. Results demonstrated that combining PLM embeddings with structural domain features consistently achieved the highest performance for both TPS detection and substrate prediction, highlighting the complementary strengths of these feature types (**Fig. S8**).

Next, we evaluated the performance of different supervised learning algorithms for TPS detection and substrate prediction. The classifiers tested included random forest, logistic regression, and feed-forward neural network as a multilayer perceptron. For all prediction tasks, random forest outperformed other classifiers (**Fig. S8**).

### Selecting sequences for experimental validation

#### Mining and Prioritizing Phylogenetically Distant TPS Sequences with Pfam/SUPFAM domains

We screened large-scale protein sequence databases BFD, UniParc, MGnify, 1KP, Phytozome, and NCBI Transcriptome Shotgun Assembly (TSA), encompassing 5.1 billion sequences. Protein sequences were filtered based on TPS-specific Pfam domains (e.g., PF01397, PF03936) and SUPFAM domain combinations (sf48239 and sf48576), resulting in a reduced set of ~200,000 TPS-like sequences. Then, an MSA of the filtered putative and characterized TPS sequences was created using MAFFT v7.526[100]. The MSA was trimmed using trimAl to remove poorly aligned regions. A phylogenetic tree was constructed using FastTree2[101]. For each uncharacterized TPS-like sequence, the phylogenetic distance to the closest characterized TPS sequence was calculated as the sum of branch lengths in the tree. Nine sequences with the highest phylogenetic distance from known TPSs were selected as high-priority candidates, assuming they represent novel functions or types (**Table S1**).

#### Mining TPS-like Sequences from UniRef90 lacking InterPro signatures

We scanned UniRef90[53] (UniProt[2] release 2022_05), containing $19.9 \times 10^7$ representative protein sequences. We have excluded sequences already annotated with InterPro signatures[3] to focus on uncharacterized sequences. We further discarded sequences shorter than 230 amino acids to remove truncated or incomplete proteins unlikely to encode catalytically competent TPSs. This conservative threshold was chosen because all experimentally verified TPSs in our curated training set are longer than 260 amino acids (**Fig. S2b**), and known crystal structures of bacterial single-domain TPSs (e.g., pentalenene synthase 337 aa[85]; epi-isozizaene synthase 361 aa[102]) confirm that the canonical α-domain fold and catalytic DDxxD/NSE motifs cannot be accommodated in shorter sequences[10]. After this filtering, $8.9 \times 10^6$ sequences remained for screening. Sequences were filtered using our TPS detection pipeline, EnzymeExplorer. The existing predicted structures in the AlphaFold Protein Structure Database[54] (AFDB) are used for the protein structure input of the pipeline, and in case the predicted structure was missing in AFDB, the sequence was skipped. With the threshold of 95% model-predicted TPS activity probability, the screening resulted in 1,056 TPS-like sequences (**Supplementary File 5**). We populated the phylogenetic tree of these TPS-like sequences with the experimentally verified TPSs in MARTS-DB. Selection for experimental validation was driven by two criteria: maximal phylogenetic distance from known TPSs and origin from historically understudied taxa (bacteria, viruses, and archaea). By applying these criteria to the phylogenetic tree, we identified several highly distant, novel clades. We then sampled 11 sequences from across these specific clades (**Fig. S10c**) to experimentally validate the model's predictive reach (**Table 1**). We evaluated each baseline method for the TPS activity detection of these sequences. Except Pfam, Foldseek, and CLEAN, each method failed to predict TPS activity for all candidates (**Table S6**). Pfam could predict only two out of seven experimentally verified candidates. Foldseek failed to predict the TPS activity for the viral sequence (UniProt ID A0A2P0VN22). Even though CLEAN was able to predict TPS activity for each experimentally verified TPS candidate, the 5-fold cross-validation showed that CLEAN's average precision is significantly lower than BLASTp, Foldseek, and EnzymeExplorer (**Figs. 2b-e, Fig. S7**). The lack of precision makes CLEAN unfavourable for screening large sequence databases for "dark putative" TPSs and selection of experimental candidates.

#### Yeast strains and cultivation

Chemicals used for media preparation were purchased from either Sigma-Aldrich, Duchefa Biochemie, Lach:ner or Penta chemicals. Solvent for metabolite extract preparation, LC- and GC-MS were purchased from Fisher Chemical and were of LC-MS grade.

The list of all strains used in the study is available in **Table S4**. *Saccharomyces cerevisiae* JWY501 derivative strains were used to express terpene synthases and produce terpenes.[52] Selective medium (SCE) used to grow transformants contained 1.92 g/L Yeast Synthetic Drop-out Medium Supplements without Uracil (Sigma-Aldrich, Y1501), 6.7 g/L Yeast nitrogen base without amino acid (Sigma-Aldrich, Y0626), and 20 g/L glucose.

DH10$\beta$ electrocompetent *Escherichia coli* cells were used for all cloning experiments. Transformed cells were selected on Lysogeny Broth (LB) with the appropriate antibiotics (ampicillin or chloramphenicol). SOC was used for recovery after electroporation.

For maintenance *S. cerevisiae* strains were cultivated in solid selective medium at 30 °C. For general preculture, *S. cerevisiae* strains were cultivated in a liquid selective medium (SCE) at 30 °C, 200 RPM in an orbital shaker. For the production run, *S. cerevisiae* strains were cultivated in selective medium SCE in 24 deep well plates (CR1426, Enzyscreen) sealed with AeraSeal (Excel Scientific), at 30 °C, 800 RPM on an Eppendorf ThermoMixer C (Eppendorf).

For plasmid amplification, *E. coli* cultures were cultivated in LB medium in 24 deep well plates (CR1426, Enzyscreen) sealed with AeraSeal (Excel Scientific), at 37 °C, 800 RPM on an ThermoMixer C (Eppendorf).

**Yeast plasmids**

The list of all plasmids used in the study is available in **Table S5.** All pTP plasmids were generated using GoldenGate assembly based on the MoClo-YTK toolkit and overhangs [103]. Part plasmids use pYTK001 as a backbone. Terpene synthases DNA parts were synthesized by Twist Bioscience as gene fragments. Cassette plasmids for genomic integration use pYTK096 as a backbone.

**pECO_Trx construction**

The pECO_Trx vector was generated from the pHIS8-4b backbone[104] by polymerase chain reaction amplification and Gibson assembly with fragments encoding the $His_8$–Trx–TEV fusion tag and a green fluorescent protein dropout cassette carrying Golden Gate-compatible overhangs.

**Polymerase chain reactions**

For colony PCR and yeast genotyping we used Phire Green Hot Start II PCR Master Mix (Thermo Fisher Scientific) with the following conditions:

In 10 µL final volume, Phire Green Hot Start II PCR Master Mix 5 µL, 25 µM forward primer 0.2 µL, 25 µM reverse primer 0.2 µL, DNA template 4.6 µL. Reactions were conducted in ProFlex 3 ✕ 32-well PCR System thermocycler (Waltham, Massachusetts, United States). Thermocycling conditions used the following template: 98°C for 2 min as initial denaturation, for 30 cycles: 98 °C for 10 s, annealing temperature for 10 s, 72 °C for 10 s.kb-1 and a final extension 72 °C for 2 minutes. PCR products were separated on agarose gel (0.8 % w/v), 130 V, 30 min.

*E. coli* colonies were selected with a toothpick and spotted 4 times on selective media. The remaining bacteria were thus resuspended in 10 µL $ddH_2O$, and boiled for 10 minutes before being used as a DNA template.

Yeast genotyping was adapted from Lõoke et al. [105]. *S. cerevisiae* colonies were selected with a toothpick and resuspended in 100 µL 200 mM LiOAc, 1 % SDS solution, and boiled for 10 minutes. Then 300 µL of EtOH 96 % were added and the solution was a vortex and centrifuge 15 000 ✕ g for 3 minutes. The supernatant was discarded and the pellet washed with 500 µL EtOH 70% before centrifugation 15 000 ✕ g for 1 min. The supernatant was discarded and the pellet dried for 1 minute at room temperature. The precipitated DNA was dissolved in 100 µL $ddH_2O$ and cell debris spined down 15 000 ✕ g for 1 min.

**Golden Gate assembly reaction**

Part plasmids were generated in 10 µL reaction volume. T4 ligase buffer 1 µL, T4 ligase 0.5 µL (M0202L, New England Biolabs), BsmBI-v2 0.5 µL (R0739L, New England Biolabs), pYTK001 0.5 µL, DNA part 0.5 µL (20 fm), ddH2O up to 10 µL. Reactions were conducted in ProFlex 3 x 32-well PCR System thermocycler (Thermo Fisher Scientific). Thermocycling conditions used the following template: for 25 cycles, 42 °C for 2 min, 16 °C for 2 min, then 60 °C for 30 min and 80 °C for 10 min.

Cassette plasmids were generated in 10 µL reaction volume. T4 ligase buffer 1 µL, T4 ligase 0.5 µL (M0202L, New England Biolabs), BsaI-HFv2 0.5 µL (R3733L, New England Biolabs), DNA parts 0.5 µL (20 fm), ddH2O up to 10 µL. Reactions were conducted in ProFlex 3 ✕ 32-well PCR

System thermocycler (Thermo Fisher Scientific). Thermocycling conditions used the following template: for 25 cycles, 37 °C for 5 min, 16 °C for 5 min, then 60 °C for 30 min and 80 °C for 10 min.

#### *E. coli* transformation

Electrocompetent cuvettes were cooled down at 4°C 30 min before the experiment. Electrocompetent 20 µL of E. coli cells were thawed at 4°C 10 min before the experiment. Once thawed, 0.5 µL of plasmid was added to the cell with gentle mixing. The electroporator was set to 1700 V. For chloramphenicol, spectinomycin and kanamycin selective markers, cells recovered in 1 mL of SOC for 1 h at 37 °C, 200 RPM. Thus, cells were concentrated to 100 µL through centrifugation 5000 × g for 3 min and plated to their respective LB + selection marker. In case of ampicillin, cells were resuspended into 100 µL of SOC after electroporation and directly plated in LB + ampicillin. Cells were then grown overnight at 37 °C.

#### Yeast transformation

Budding yeast strains (**Table S4**) were generated by standard lithium acetate transformation protocol [106] and selected using auxotrophy. For genomic integration, 500-1500 ng of Plasmid were digested using NotI-HF in 10 µL final volume. For plasmid integration, 500-1500 ng of plasmid were used. Cells were precultured overnight in YPD medium, 30 °C, 200 RPM. Then, cells were diluted to OD 0.1 and cultivated in YPD medium, 30 °C, 200 RPM until they reached OD 0.5. For 5 mL of cells at OD 0.5: the cells were pelleted, 2500 × g, 5 min and washed in 5mL sterile sorbitol wash buffer (600 mM sorbitol, 100 mM $K_2HPO_4$). Cells were pelleted again 2500 × g, 5 min and resuspended in 1 mL sterile sorbitol wash buffer, and pelleted again 16 000 × g, 1 min and resuspended in 1 mL TE/LiAc. Then, cells were concentrated in 100 µL TE/LiAc and 10 µL of digested plasmid and 10 µL of Salmon sperm were added, mixed gently and incubated for 10 min at room temperature. Then, 260 µL of TE/LiAc/PEG (40 % w/v) was added and the cells were incubated for 45 min at 42 °C. Cells were then centrifuged 6000 × g for 1 min and washed in water before plating on selective medium. Cells were grown for 3-5 days at 30 °C.

#### Metabolite extraction

Cells were maintained in solid selective medium and preculture in 1 mL selective medium in 96 deep well plate (CR1496, Enzyscreen) sealed with AeraSeal (Excel Scientific), at 28 °C, 1500 RPM on an ThermoMixer C (Eppendorf) for 1 day. Cells were seeded at OD 0.05 in 2.2 mL in inducible selective medium SCE (10 % glucose, 90 % galactose) in 24 deep well plates (CR1426, Enzyscreen, Hamburg, Germany, Netherlands) sealed with AeraSeal (Excel Scientific), at 28 °C, 300 RPM for 48 hours. Then, 1 mL of ethyl acetate (E196-4, Fisher Scientific) was added to the culture medium and mixed for 1 hour at 28 °C, 250 RPM. Then 1 mL of ethyl acetate was added and thoroughly mixed by pipetting. The sample was collected in a 2 mL round bottom tube (Eppendorf) centrifuged for 5 mins, 14 100 × g and the organic phase was carefully collected and dried under N2 flow.

#### LC-MS analyses

For LC-MS analysis, samples were resuspended in 100 µL ethyl acetate. LC-MS analyses were performed using Vanquish Flex UHPLC System interfaced to an Orbitrap ID-X Tribrid mass spectrometer, equipped with heated electrospray ionization (Thermo Fisher Scientific). The LC conditions were as follows: column, Waters BEH (Ethylene Bridget Hybrid) C18 50 × 2.1 mm, 1.7 µm; mobile phase, (A) water with 0.1 % formic acid; (B) acetonitrile with 0.1% formic acid; flow rate, 350 µL.min-1; column oven temperature, 40 °C, injection volume, 1 µL, linear gradient of 5 to 100 % B over 5 min and isocratic at 100 % B for 2 min. Electrospray ionization was achieved in positive mode and mass spectrometer parameters were as follows: ion transfer tube temperature, 325 °C, auxiliary gas flow rate 10 L.min-1, vaporizer temperature 350 °C; sheath gas flow rate, 50 L.min-1; capillary voltage, 3000 V, MS resolution 60 000, quadrupole isolation, scan range from m/z 100-1000, RF Lens 45 %, maximum injection time 118 ms. LC-MS .raw data files were directly imported into mzmine 4.10.0 [107]. Extracted ion chromatograms for compounds of interest were generated using the raw data overview feature and exported as .pdf.

#### GC-MS analyses

For GC-MS analysis, samples were resuspended into 100 µL ethyl acetate. GC-MS analyses were performed using a 7890A gas chromatograph coupled with a 5975C mass spectrometer, equipped with electron ionization (EI) and quadrupole analyzer (Agilent Technologies). The samples (1 µl) were injected into a split/splitless inlet in split mode (split ratio 10:1). The injector temperature was 250 °C. A DB-1ms fused silica capillary column (30 m ✕ 250 µm; a film thickness of 0.25 µm, J&W Scientific) was used for separation. The carrier gas was helium at a constant flow rate of 1.0 ml/min. The temperature program was: 40 °C (1 min), then 5 °C.min-1 to 100 °C, followed by 15 °C.min-1 to 230 °C . The temperatures of the transfer line, ion source and quadrupole were 320 °C, 230 °C and 150 °C, respectively. EI spectra (70 eV) were recorded from 25 to 500 m/z. GC-MS .dx files were analyzed using OpenLab CDS 2.4. Extracted ion chromatograms for compounds of interest were exported as .csv files and built using the ggplot2 package in R. Figures were generated using Rstudio and the following packages: ggplot2, gridExtra, patchwork, dplyr, forcats, ggthemes, ggprism, DescTools, tidyverse, scales and Adobe Illustrator CS6. Peaks for sesquiterpene (m/z 204) and diterpene (m/z 272) were retrieved from extracted ion chromatograms using OpenLab CDS 2.4. The MS spectra from each peak were therefore queried to NIST EI search software, with NIST 2023 EI database. Both spectrum from the experimental data and its best match on the NIST EI database were exported as .MSPEC files. MSPEC files were converted to mgf files using a python script and imported into mzmine 4.10.0 and compared as a mirror plot [107].

#### Isolation of compound 12

Yeast strain JWY501[52] carrying plasmid TP0255µ was cultivated in 8 L of selective medium with 10 % glucose and 90 % galactose for autoinduction. After 3 days at 30 °C, 200 RPM, supernatant was gathered after 30 min centrifugation, 5000 ✕ g at 4 °C and extracted using cyclohexane at room temperature and the organic layer was dried *in vacuo*. Extract was first fractionated using normal phase flash chromatography with silica gel using n-hexane with increasing amounts of EtOAc as the mobile phase. Fractions containing terpenes were then further fractionated on normal phase semi preparative chromatography with Hexane:EthylAcetate solvent to yield pure terpene **12** (1 mg).

#### NMR spectroscopy

NMR spectra of compound **12**, 1H and 13C APT, have been acquired on a Bruker Avance III 600 MHz spectrometer equipped with 1.7 mm micro-cryoprobe, using CDCL3 as solvent.

#### *In vitro* assay of A0A5E4I9B1

The coding sequence corresponding to UniProt accession A0A5E4I9B1 was cloned into the pECO_Trx expression vector to produce an N-terminal $His_8$–thioredoxin fusion protein. Because soluble protein was not detected under the initial extraction condition, protein solubility was evaluated using ten conditions from the previously described buffer-screening workflow[108]. Buffer B5, containing 50 mM HEPES, pH 7.0, 50 mM NaCl and 100 mM urea, was used for subsequent protein production, extraction, purification and resuspension.

The recombinant protein was produced at a 10-ml culture scale. Cells were extracted in buffer B5, and the soluble fraction was purified by centrifugation-based nickel–nitrilotriacetic acid affinity chromatography according to the previously described protein-production and purification workflow. The purified protein was resuspended in buffer B5 and adjusted to 1 mg $ml^{-1}$.

Enzyme activity was assessed using farnesyl diphosphate as the substrate. Trisammonium farnesyl diphosphate was dissolved at 0.2 mg $ml^{-1}$ in 25 mM ammonium bicarbonate. Each 250-µl reaction contained 40 µl substrate solution, 40 µl enzyme solution and 170 µl reaction buffer. The reaction buffer contained 50 mM HEPES, pH 7.0, 50 mM NaCl, 100 mM urea, 10 mM $MgCl_2$ and 20% glycerol. Reactions were incubated overnight at 28 °C with shaking. Assays were performed in duplicate, and an equivalent reaction containing heat-inactivated enzyme was included as a negative control. Following incubation, reaction mixtures were extracted with pentane. The recovered pentane phase was transferred directly into gas chromatography vials without evaporation or concentration.

#### GC-TOF for *in vitro* assay

Pentane extracts were analyzed using a Pegasus BTX gas chromatography–time-of-flight mass spectrometry system (LECO). Separation was performed using an Rxi-5MS capillary column with a nominal length of 30 m, an internal diameter of 0.25 mm and a film thickness of 0.25 µm. The carrier-gas flow was maintained at 1.40 ml $min^{-1}$ throughout the analysis.

Samples were injected at 240 °C using split injection with a split ratio of 100:1. The septum purge flow was 3 ml $min^{-1}$, and the split flow was 140 ml $min^{-1}$. The oven was initially maintained at 40 °C for 1 min, increased to 90 °C at 15 °C $min^{-1}$, and then increased to 280 °C at 35 °C $min^{-1}$ and maintained for 3 min. The transfer line was maintained at 280 °C.

Mass spectra were acquired using electron ionization at 70 eV. The ion-source temperature was 250 °C, and the emission current was 1.0 mA. Terpene products were assigned by comparison of their chromatographic retention times and mass spectra with those obtained from the terpene-producing yeast positive reference.

### Taxonomic analysis of archaea

Assemblies of Archaea genomes were retrieved from the representative genomes in Genomic Taxanomy Database[64] (GTDB release 232). The sequences are filtered by the length 200-1200 amino acids. Then we eliminated confident negatives by using EnzymeExplorer with sequence-only features (TPS detection score > 0.1) since folding the complete dataset would be computationally too expensive. This preliminary filtering yielded 46,361 sequences. We used ESMFold[21] to fold these sequences, and screened them using our predictive pipeline.

### Multiple sequence alignment

Multiple sequence alignment was done using MAFFT v7.526[100] with default settings. Multiple sequence alignment was visualized using espript 3.0[109].

### Archaeal TPS structure analysis and ligand docking

To create a structure model of A0A5E4I9B1, we employed AlphaFold 3[20]. The model was generated using the A0A5E4I9B1 sequence and three $Mg^{2+}$ ions as cofactors. Attempts to include the diphosphate moiety during modelling resulted in poor models. In A0A5E4I9B1, the well-known $Mg_B$ and $Mg_C$ metal binding motif is $^{221}$**N**TLN**T**WPR**E**. In the original AlphaFold 3 model, the N/T/E residues of this motif do not interact with the $Mg_B$ and $Mg_C$ ions, and hence we refined the model. To this end, we first docked the diphosphate moiety to complete the polar region of the binding site prior to modelling the AlphaFold 3 model. The diphosphate moiety was docked into its standard binding pocket[10] using Nuclear Overhauser Effect (NOE) restraints between the $Mg^{2+}$ ions and the diphosphate oxygens, as implemented in EnzyDock[59,62]. EnzyDock is a CHARMM-based multistate, multiscale docking program for enzymes[110]. Subsequently, we refined the AlphaFold 3 model using a CHARMM-based minimization protocol designed to create an active holo-form of TPSs. The protein was fixed except for the helix hosting the $^{221}$**N**TLN**T**WPR**E** motif and neighboring loop regions (i.e., the flexible residues were 195-237) and the diphosphate-$(Mg^{2+})_3$ cluster were flexible. We applied a series of TPS-specific NOE restraints during stepwise minimization of the AlphaFold 3 model to form the conserved TPS interactions between A0A5E4I9B1 and the diphosphate-$(Mg^{2+})_3$ cluster. The protocol included 1,000 steps of steepest descent and 500 steps of adopted basis Newton-Raphson minimization. Hydration was included via the continuum GBSW model[111].

To model reaction states in A0A5E4I9B1, we employed EnzyDock. These docking simulations were performed using the above-mentioned refined AlphaFold 3 model, which includes the protein, the diphosphate moiety, and three $Mg^{2+}$ ions. The cationic isoprenoid tail of FPP and the main product β-bisabolol were docked into the active site. The substrate was docked as an (E,E)-farnesyl cation and a covalent bond was formed between its C1 position and either the O1α or O2α oxygens of the diphosphate on the fly during docking. Binding to O1α was favorable (**Table S3**), as has been observed for plant TPS[59]. *β*-bisabolol was docked in all four diastereomeric forms. When docking FPP, a water molecule was first docked in a position between D63 and the diphosphate moiety. The native protein, cofactors, and ligands were described by the CHARMM36m[112] and CGenFF force fields[113]. The nonbonded interactions of the enzyme-ligand complexes were treated using a grid potential generated around the centre of the active site of the protein for both van der Waals and electrostatic interactions. The centre of the grid was taken as the centre of the active site. In the present study, the grid was calculated with a grid spacing of 0.25

Å and dimensions of 15×15×15 $Å^3$. The size of the grid was chosen so that it encompasses the active site.

To compute the matching ligand states along the reaction path[114], we applied Pathfinder that comes with EnzyDock, and matches neighbouring reaction states (e.g., FPP → *β*-bisabolol) using RMSD criteria.

#### Use of artificial intelligence tools

ChatGPT (OpenAI; GPT-5.5) was used to assist with language editing and improving the clarity and readability of author-written text. Claude Code (Anthropic) was used as a coding assistant during software implementation, including assistance with code generation, refactoring, and debugging. All AI-generated suggestions and code were reviewed, modified where necessary, and validated by the authors. These tools were not used to generate experimental data, perform autonomous scientific analyses, interpret results, or formulate the scientific conclusions of the study. The authors take full responsibility for the final manuscript and software.

### Supplementary Figures

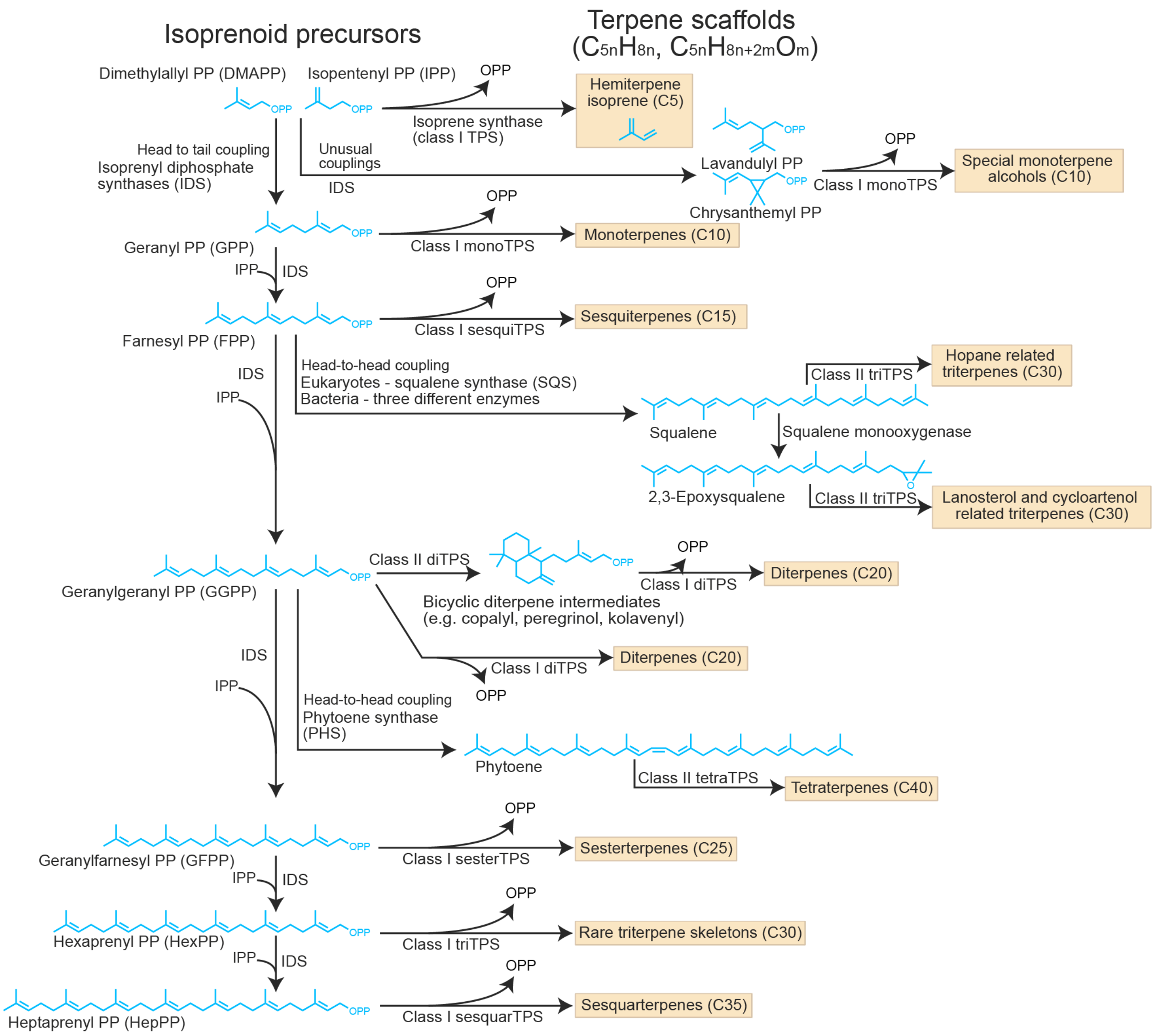


**Fig. S1 | Summary of TPS reaction types, substrates and product classes.** TPS substrates are synthesized by isoprenyl diphosphate synthases, enzymes that catalyze the sequential head-to-tail condensation of isoprene units (C5) in the form of isopentenyl diphosphate. By the number of isoprene units, terpenes are divided into types (hemi, mono, sesqui, di, etc.) where monoterpenes are composed of two isoprene units. Similar enzymes also catalyze other manners of condensations of isoprenyl diphosphates into products like squalene. The cyclization of those substrates into terpene products is guided by terpene synthases. Class I terpene synthases initiate reactions by abstracting the diphosphate group while class II terpene synthases initiate their reactions by protonation of a terminal alkene or of an epoxide group.

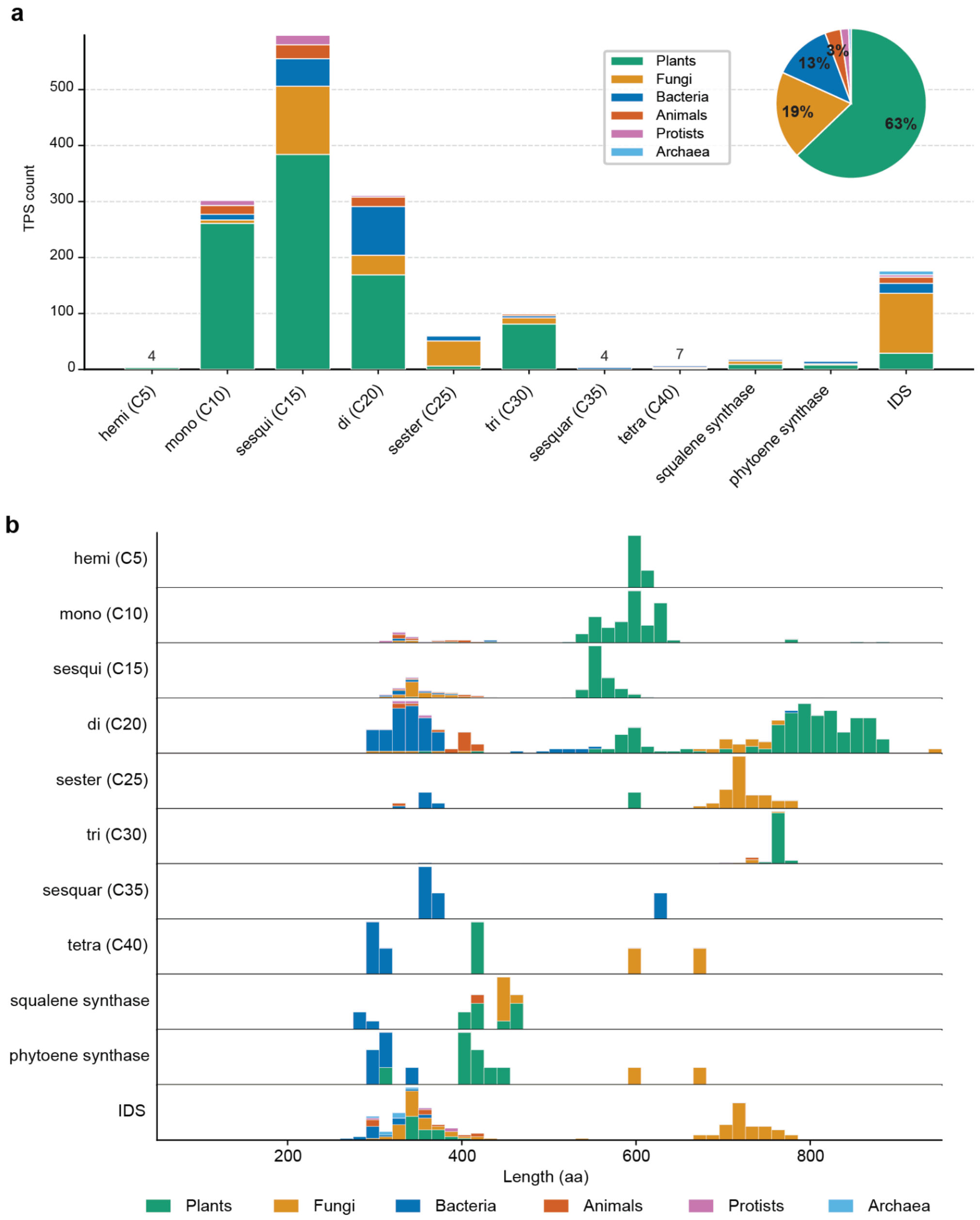


**Fig. S2 | Descriptive statistics of the curated TPS & IDS dataset. a**, Plot showing the number of reactions catalyzed by TPSs or IDSs in MARTS-DB (**Supplementary File 1**) divided into reaction types and taxonomic kingdoms of the source organisms. Note that in cases where a single enzyme catalyzes multiple types of reactions (e.g., sesquiTPS and diTPS), such an enzyme is counted multiple times in corresponding columns. **b**, The distribution of protein sequence length (amino acid residues) per TPS types and kingdoms.

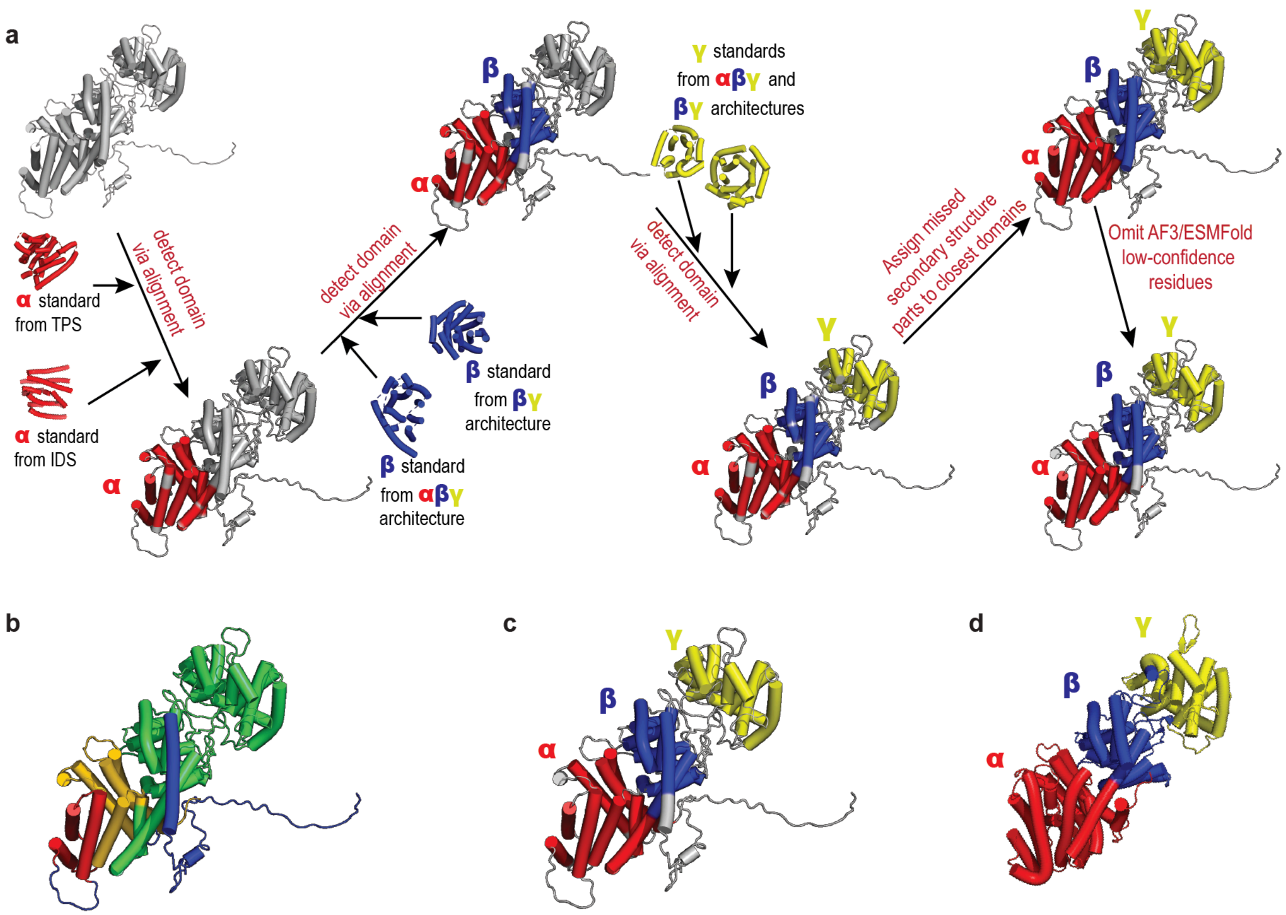


**Fig. S3 | The workflow for protein structure segmentation into domains. a**, A high-level overview of our pipeline for structure segmentation into domains. The pipeline uses alignment to known structural templates to partition predicted protein structures into distinct domains. **b**, An example of state-of-the-art (SOTA) general segmentation[115] failing to partition the AFDB structure of a TPS, demonstrated on a randomly selected TPS with UniProt accession B9GSM9. Moreover, the SOTA method does not assign domain types to the segmented regions, resulting in incomplete domain characterization. **c**, Segmentation result using our algorithm for the same UniProt accession B9GSM9. The TPS with an αβγ architecture is correctly segmented, with domains assigned to their corresponding types: α, β, and γ. **d**, Ground truth segmentation of a TPS with αβγ architecture (UniProt accession Q41594) into CATH domains: 3p5rA01, 3p5rA02, and 3p5rA03.

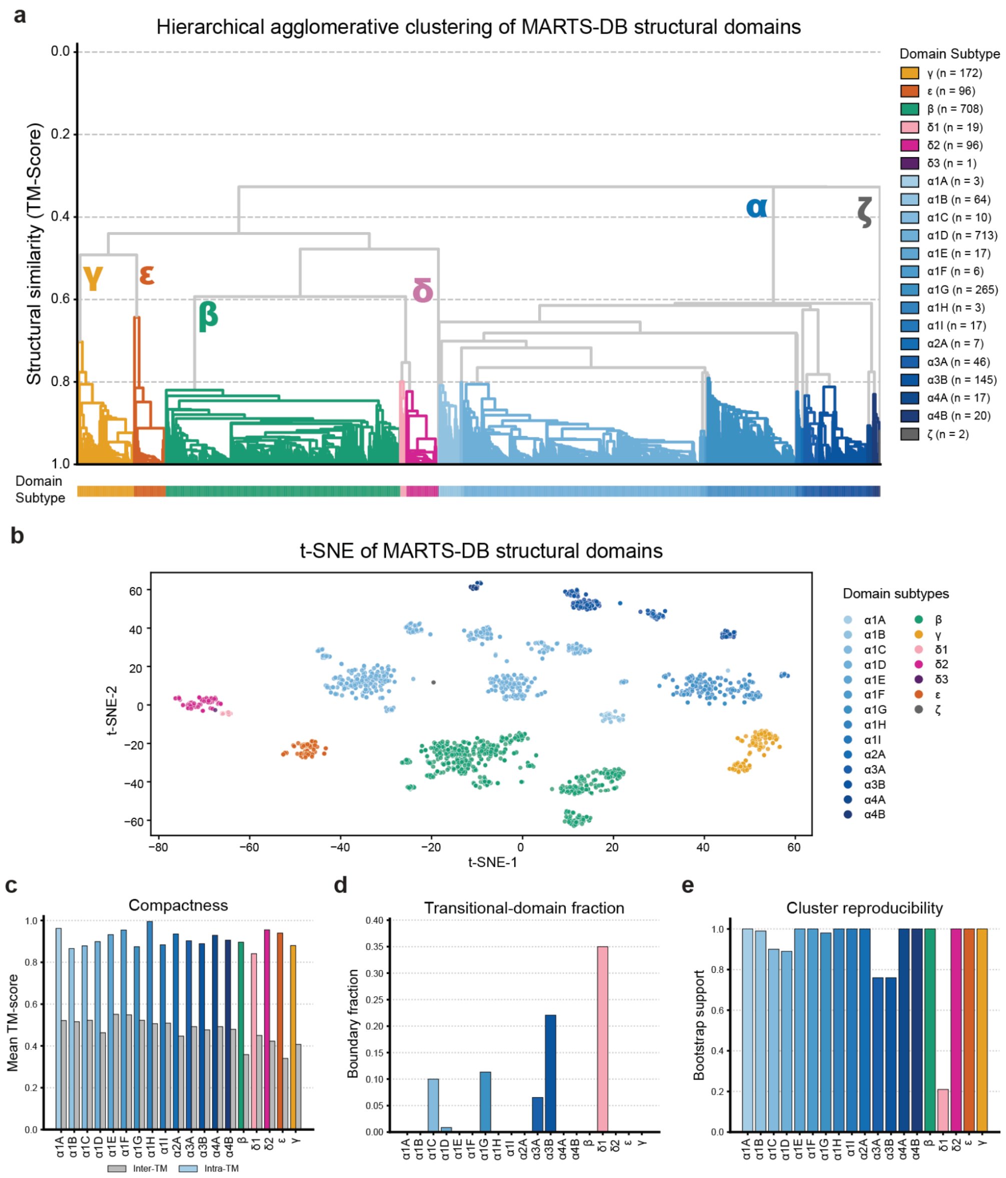


**Fig. S4 | Determination and evaluation of domain subtypes. a**, Dendrogram derived by unweighted pair group method with arithmetic mean (UPGMA) from pairwise structural-similarity TM-scores calculated for all detected domains in MARTS-DB. The subtype clades are determined by Dynamic Tree Cut[30] (min-cluster-size = 3, deep-split=0), and named manually as the final step. ζ and δ3 domains are missed by Dynamic Tree Cut due to small cluster size and labeled manually due to the significant structural dissimilarity. The dendrogram shows the structural distinction between the major domain types α, β, γ, δ, ε, and ζ. **b**, t-SNE visualization of all MARTS-DB domains with the distance metric of TM-Score from the pairwise structural alignments. The scatter plot supports the subtypes derived by the Dynamic Tree Cut algorithm. **c**, Compactness analysis for each domain subtype cluster showing mean intra (colored) and inter (gray) TM-scores for each subtype. **d**, Transitional fraction analysis for each cluster. A domain is flagged as transitional if TM-score to its own subtype's medoid is within 0.05 of the closest rival subtype's medoid. **e**, Cluster reproducibility analysis assessed by a subsampling-jackknife bootstrap for 100 iterations. For each iteration 80% of the domains are randomly sampled, and the dendrogram is built from scratch. Bootstrap support measures in how many of the iterations the surviving subtype members form a single monophyletic subtree in the bootstrap tree.

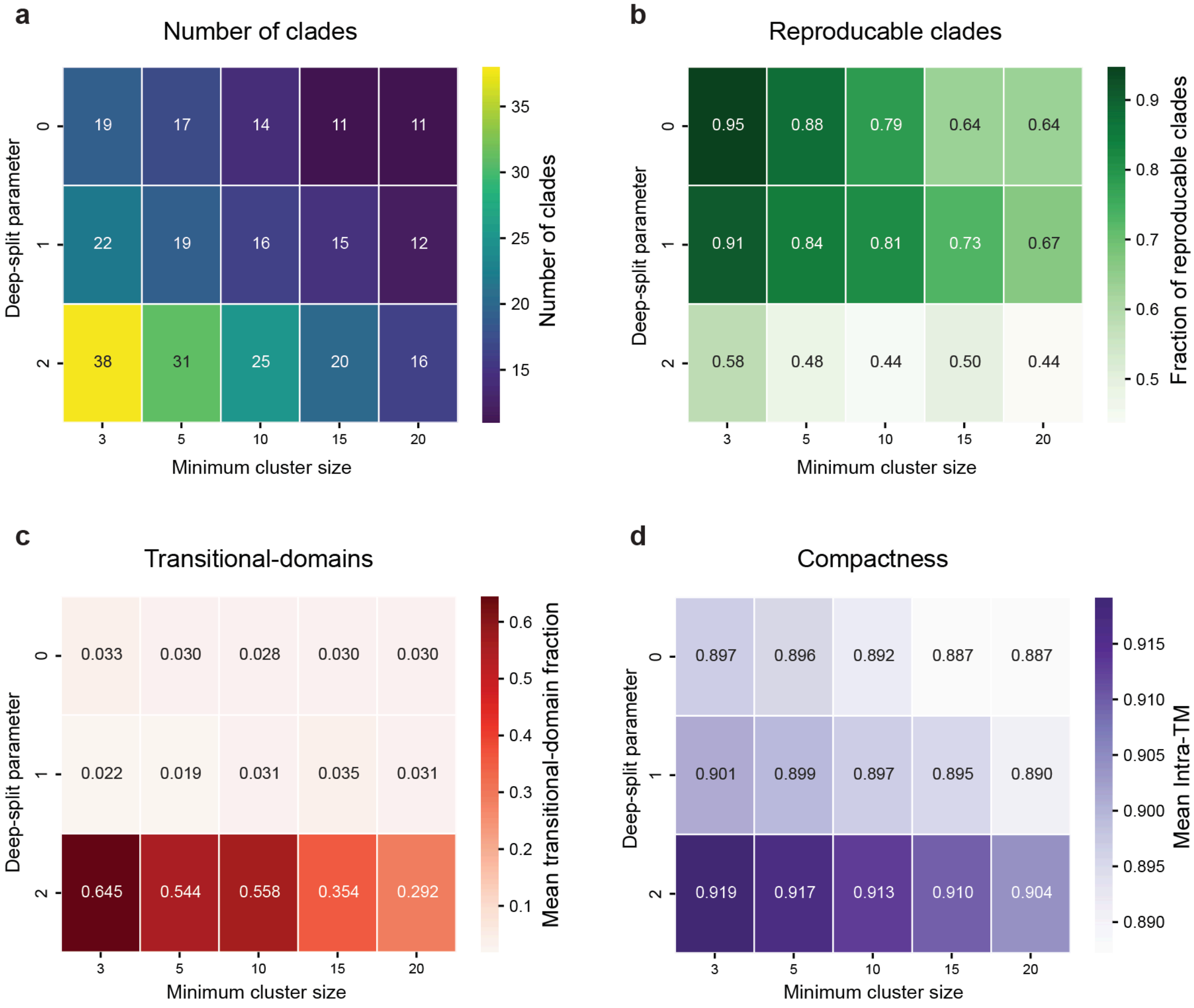


**Fig. S5 | Determination of Dynamic Tree Cut hyperparameters by a combinatorial sweep.** The figure shows the evaluation of Dynamic Tree Cut[30] algorithm with varying parameters applied on the raw dendrogram of MARTS-DB structural domains **(Fig. S4a)**. The sweep is applied by the deep-split parameters {0, 1, 2} and minimum cluster sizes {3, 5, 10, 15, 20}. Based on manual inspection of these evaluation metrics, the parameters 0 and 3 are chosen respectively. **a**, Number of final clades picked by Dynamic Tree Cut. **b**, Heatmap of the fraction of clades with the bootstrap support larger than 0.7. Bootstrap support for each clade is calculated separately for each application of Dynamic Tree Cut and by the same approach as in **Fig. S4e**: a subsampling-jackknife bootstrap for 100 iterations. For each iteration, 80% of the domains are randomly sampled, and the dendrogram is built from scratch. Bootstrap support measures for how many of the iterations the surviving subtype members form a single monophyletic subtree in the bootstrap tree. The parameters (0, 3) show highest reproducibility by far. **c**, Heatmap of the weighted average of transition-domain fraction across all determined clades. In case of deep-split parameter = 2, the transitional-domain fractions explode, making the subtype determination very unstable. **d**, Heatmap of weighted average of mean intra-TM across all determined clades. The deep-split parameter = 2 results with a better overall compactness, however much less stable clades as seen in **b** and **c**. The difference between deep-split parameters being 0 and 1 is minor and therefore interpreted as negligible.

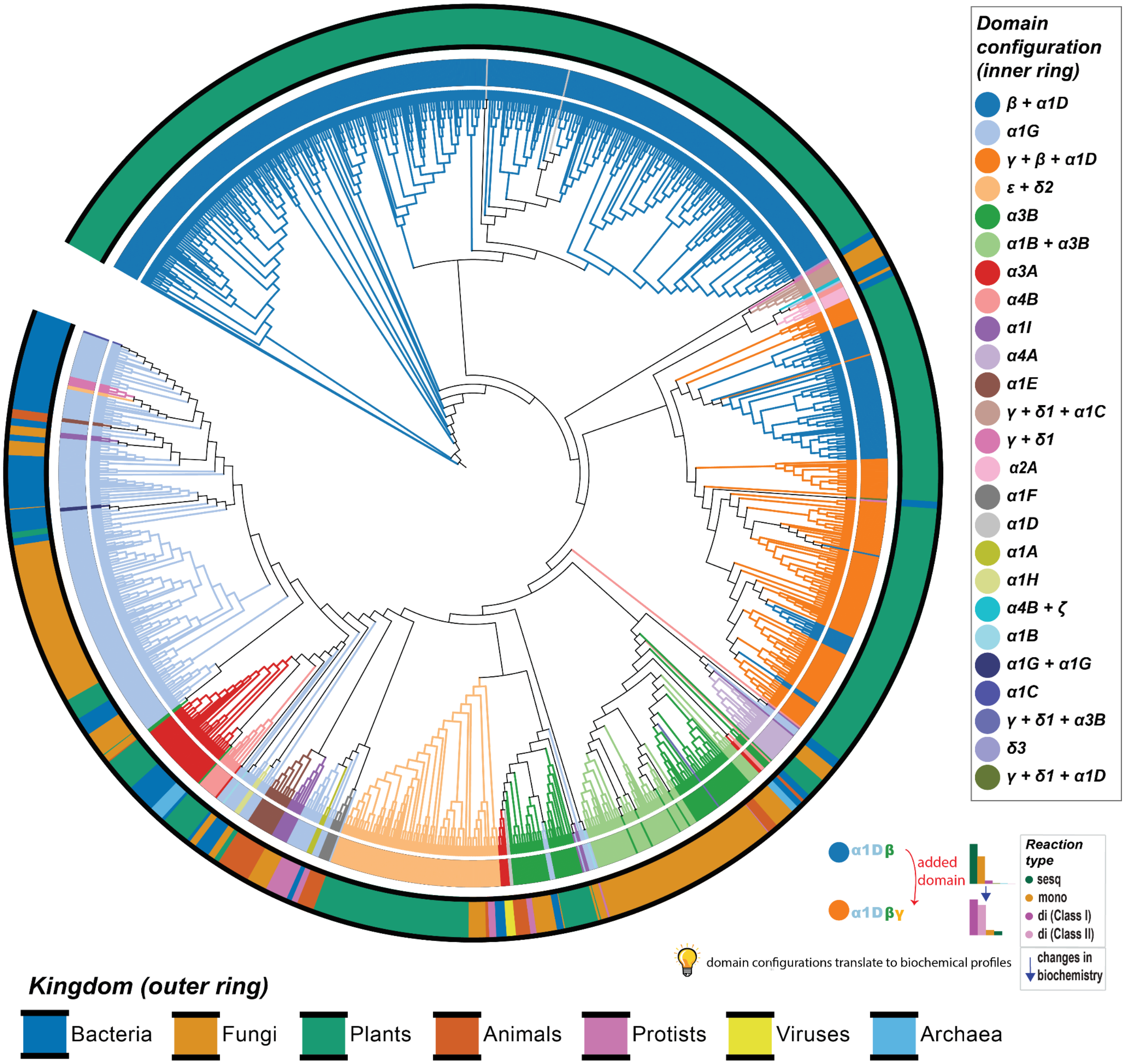


**Fig. S6 | Domain subtypes mapped onto the phylogenetic tree of characterized TPSs.**

The figure highlights the consistency of discovered domain configurations with the evolutionary relationships of corresponding enzymes. The inner color ring represents the identified domain configurations for the TPSs and IDSs in MARTS-DB (**Supplementary File 1**), while the outer color ring denotes the corresponding kingdoms. Comparison with **Fig. 1f** illustrates how changes in domain configurations align with modifications in the biochemical activity profiles of TPSs. This analysis emphasizes the evolutionary and functional significance of the discovered domain architectures. For instance, plant TPSs with the configuration α1D-β-γ are capable of diTPS synthesis, unlike those with the domain configuration α1D-β (see **Fig. S1** for the definitions of the TPS classes)

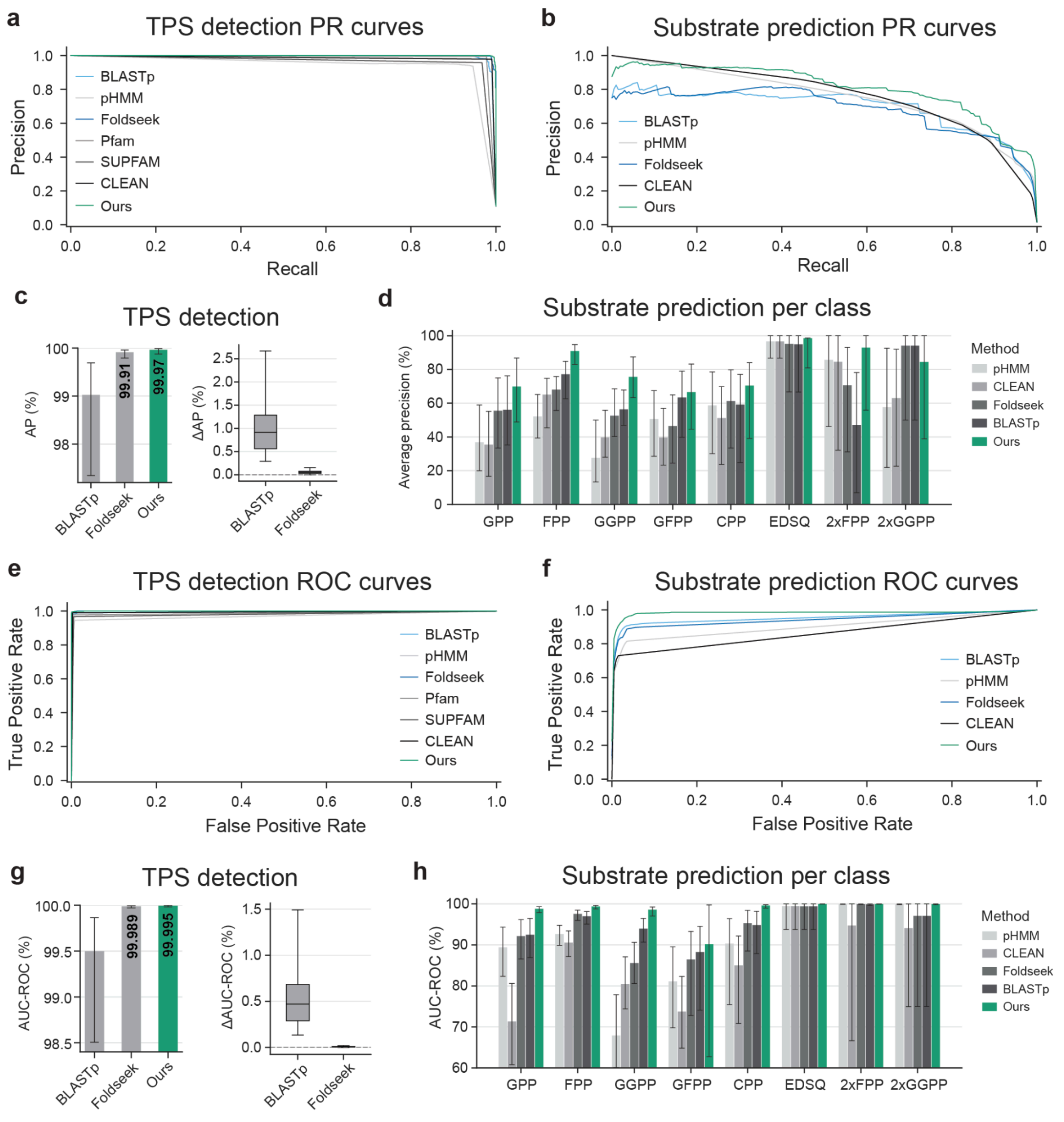


**Fig. S7 | Comprehensive evaluation of TPS detection and substrate prediction models. a**, Precision-recall (PR) curves for TPS detection methods. **b**, Macro-average PR curves for TPS substrate prediction methods. **c**, Average precision (AP) comparison for TPS detection of the top three performing methods. The bar plot describes APs with the error bars as BCa 95% confidence intervals (CI) calculated via pooled out-of-fold (OOF) block-bootstrapping. The forest plot describes the distribution of AP differences ($AP_{EnzymeExplorer}$ - $AP_{Baseline}$) derived by the pairwise block-bootstrapping. The boxes span the inner 25–75% percentile range of the bootstrap distribution. The whiskers represent BCa 95% CIs, and the medians are represented by the horizontal lines inside the box. **d**, APs for substrate prediction per substrate class of each substrate prediction method. Error bars represent the BCa 95% CIs calculated via pooled OOF block-bootstrapping. **e**, Receiver Operating Characteristic (ROC) curves for TPS detection methods. **f**, ROC curves for TPS substrate prediction methods. **g**, Area under the ROC curve (AUC-ROC) comparison for TPS detection by the top three performing methods, analogous to **c**. **h,** AUC-ROCs for substrate prediction per substrate class, analogous to **d**.

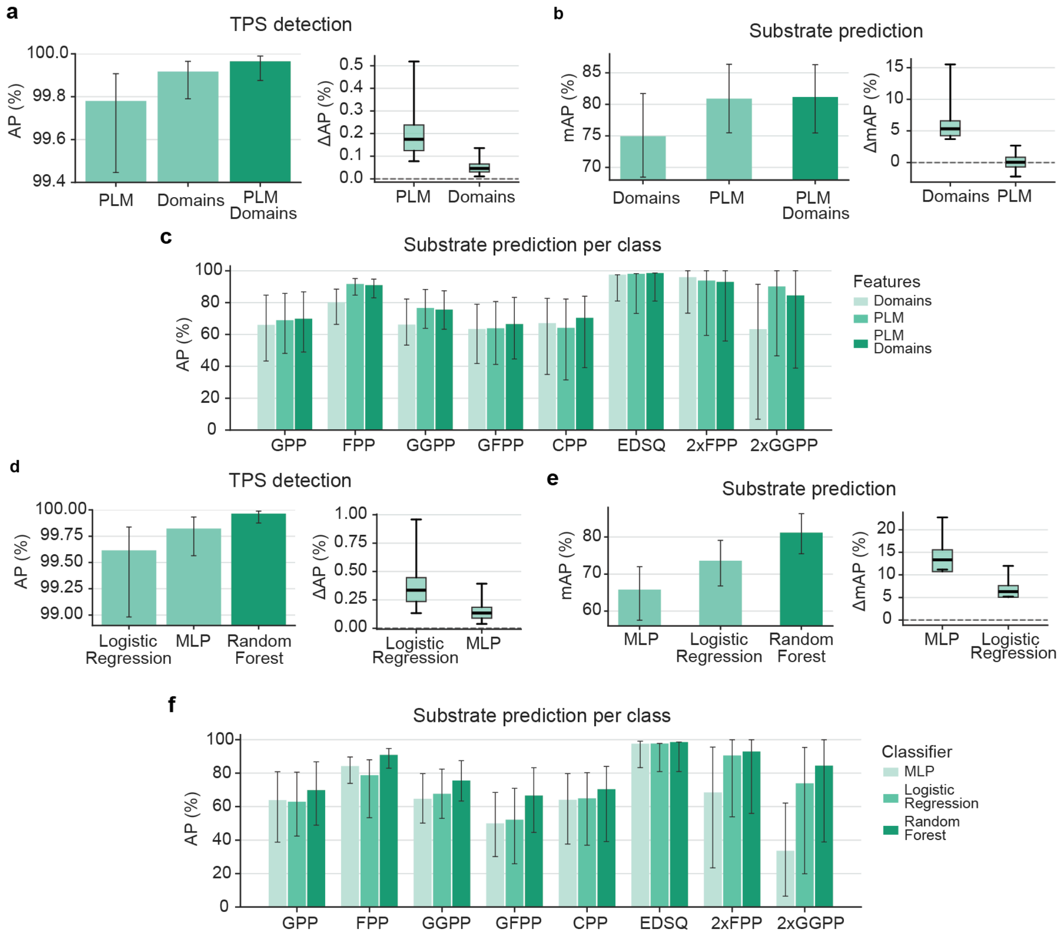


**Fig. S8 | Ablation study evaluating the impact of different input features on TPS detection and substrate prediction. a**, Average precision (AP) comparisons for TPS detection across different feature sets. The bar plot describes APs with the error bars as BCa 95% confidence intervals (CI) calculated via pooled out-of-fold (OOF) block-bootstrapping. The forest plot describes the distribution of AP differences ($AP_{PLM\&Domains}$ - $AP_{PLM}$) derived by the pairwise block-bootstrapping. The boxes span the inner 25–75% percentile range of the bootstrap distribution. The whiskers represent BCa 95% CIs, and the medians are represented by the horizontal lines inside the box. **b**, Mean average precision (mAP) comparisons for substrate prediction across different feature sets, analogous to **a**. **c**, APs for substrate prediction per substrate class. Error bars represent the BCa 95% CIs calculated via pooled OOF block-bootstrapping. **d**, AP comparisons for TPS detection across different classifiers. The bar plot describes the BCa 95% CI calculated via pooled OOF block-bootstrapping, analogous to **a**. **e**, mAP comparisons for substrate prediction across different classifiers, analogous to **a**. **c**, APs for substrate prediction per substrate class, analogous to **c**.

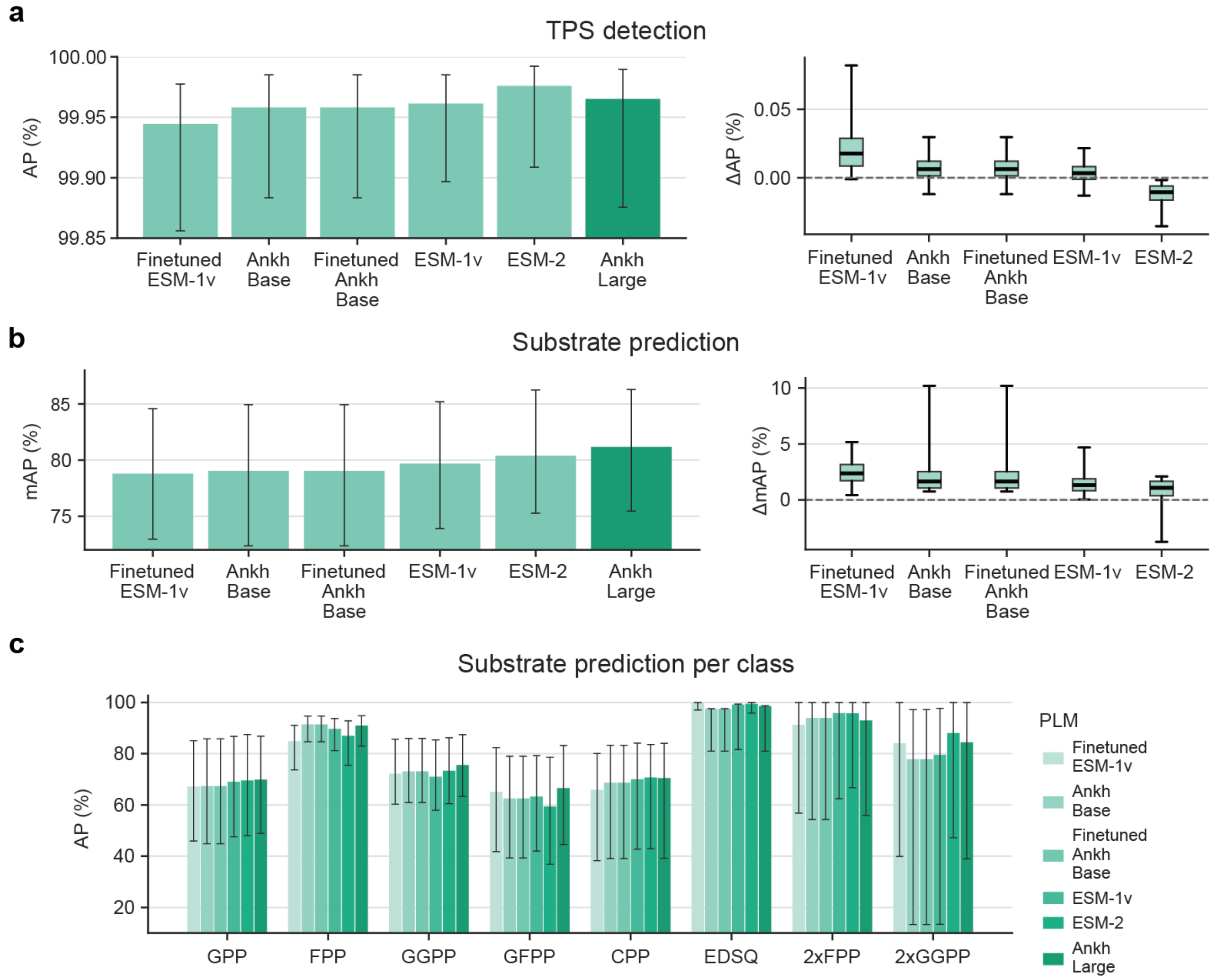


**Fig. S9 | Ablation study evaluating the impact of different protein language models (PLM) for TPS detection and substrate prediction.** For ESM-2 the esm2_t36_3B_UR50D model is used. **a**, Average precision (AP) comparisons for TPS detection across different PLMs. The bar plot describes APs with the error bars as BCa 95% confidence intervals (CI) calculated via pooled out-of-fold (OOF) block-bootstrapping. The forest plot describes the distribution of AP differences ($AP_{\text{Ankh-large}}$ - $AP_{\text{Ankh-base}}$) derived by the pairwise block-bootstrapping. The boxes span the inner 25–75% percentile range of the bootstrap distribution. The whiskers represent BCa 95% CIs, and the medians are represented by the horizontal lines inside the box. **b**, Mean average precision (mAP) comparisons for substrate prediction across different PLMs, analogous to **a** **c**, APs for substrate prediction per substrate class. Error bars represent the BCa 95% CIs calculated via pooled OOF block-bootstrapping. Ankh-large performs consistently better than ESM-2 for the majority substrate classes (GPP, FPP, GGPP).

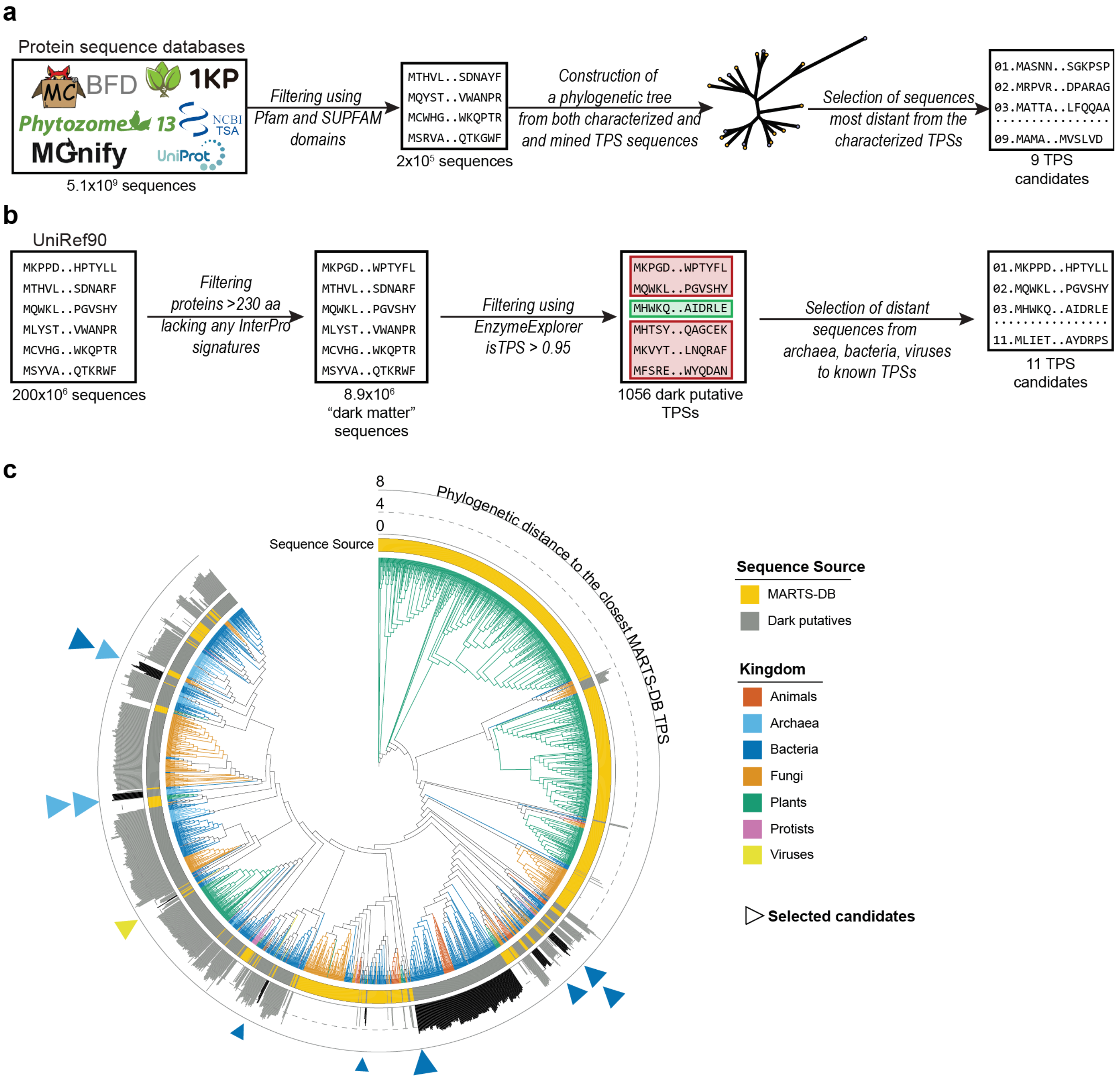


**Fig. S10 | Selection of candidate TPS sequences for experimental expression.** **a**, *In silico* screening workflow for selecting evolutionary distant uncharacterized TPS with TPS-specific Pfam/SUPFAM domains. **b**, *In silico* screening workflow for selecting TPS that lack any InterPro signatures[3]. **c**, Phylogenetic analysis for the candidate selection procedure in **b**. The figure shows the joint phylogenetic tree of dark putative TPSs and MARTS-DB TPSs. Clades are colored by the kingdom of respective sequences. The inner strip is colored with yellow for MARTS-DB TPSs, and gray for dark putative TPSs. The outer strip contains the phylogenetic distance bar plot for the dark putative TPSs, which is calculated by summation of branch lengths. The bars for manually selected clades of understudied kingdoms (viruses, archaea, and bacteria) are colored in black. The TPS candidates selected from these clades are shown with triangle markers, colored by the respective kingdoms and sizes scaled by their phylogenetic distance to the closest MARTS-DB TPS.

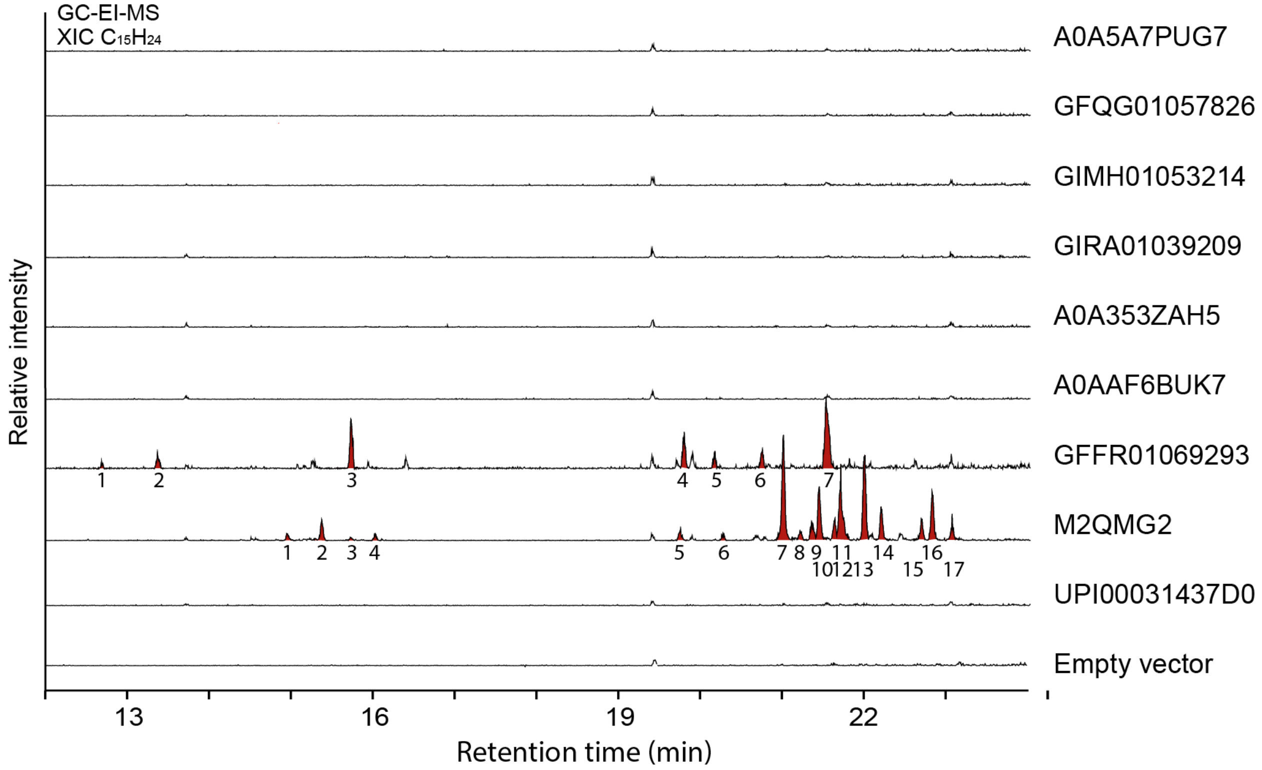


**Fig. S11 | GC-MS extracted ion chromatograms of the molecular mass of $C_{15}H_{24}$.** Results of yeast expressions of proteins shown in **Table S1** (protein accession IDs shown on the right side). Detected sesquiterpene ions are highlighted in red.

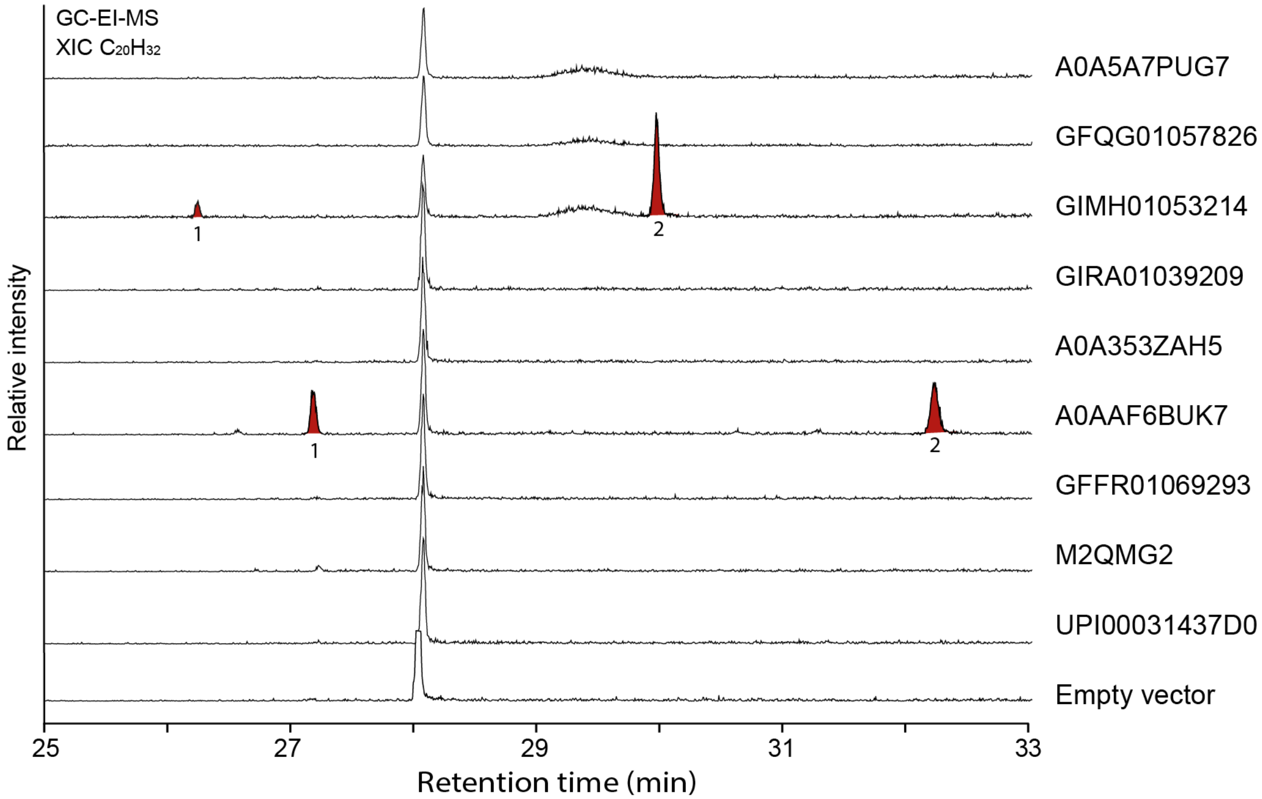


**Fig. S12 | GC-MS extracted ion chromatograms of the molecular mass of $C_{20}H_{32}$.** Results of yeast expressions of proteins shown in **Table S1** (protein accession IDs shown on the right side). Detected diterpene ions are highlighted in red.

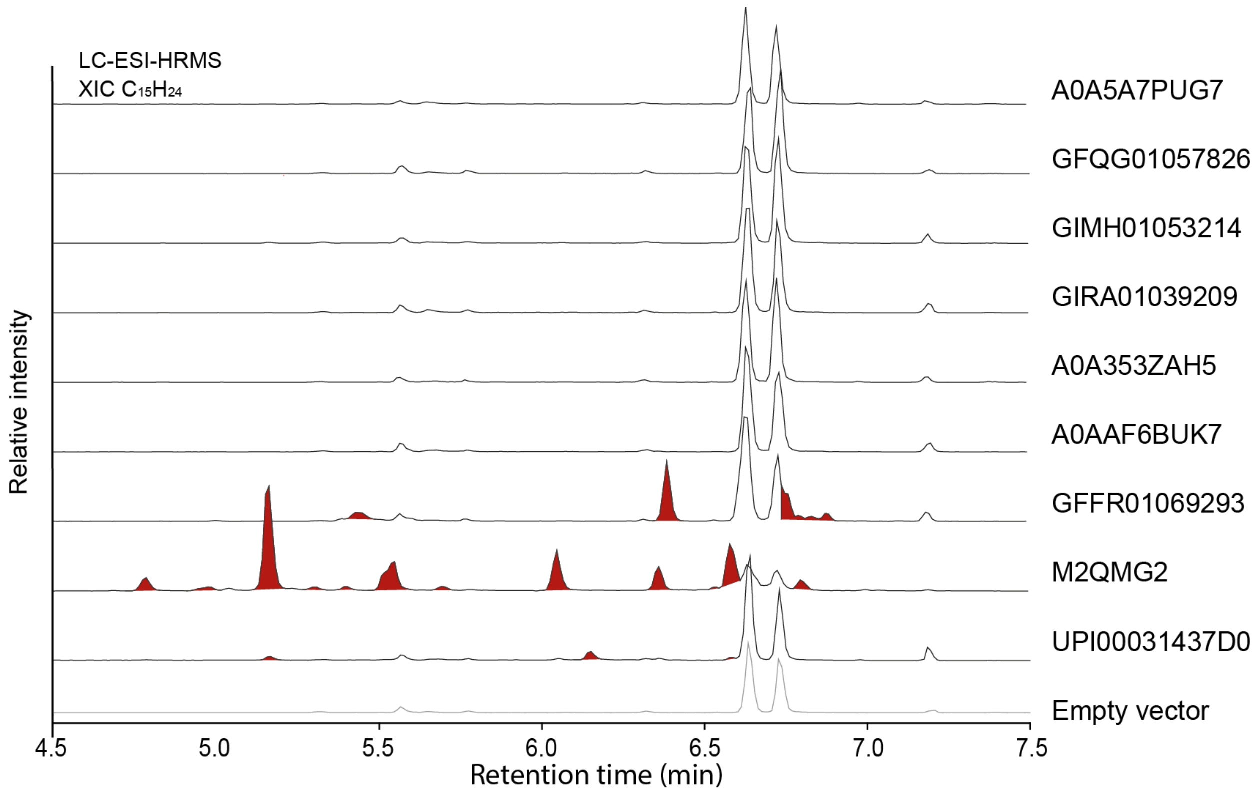


**Fig. S13 | LC-MS extracted ion chromatograms of the molecular mass of $[C_{15}H_{24}+H]^+$ ions at *m/z* 205.1951 (+/- 5 ppm).** Results of yeast expressions of proteins shown in **Table S1** (protein accession IDs shown on the right side). Detected sesquiterpene ions are highlighted in red.

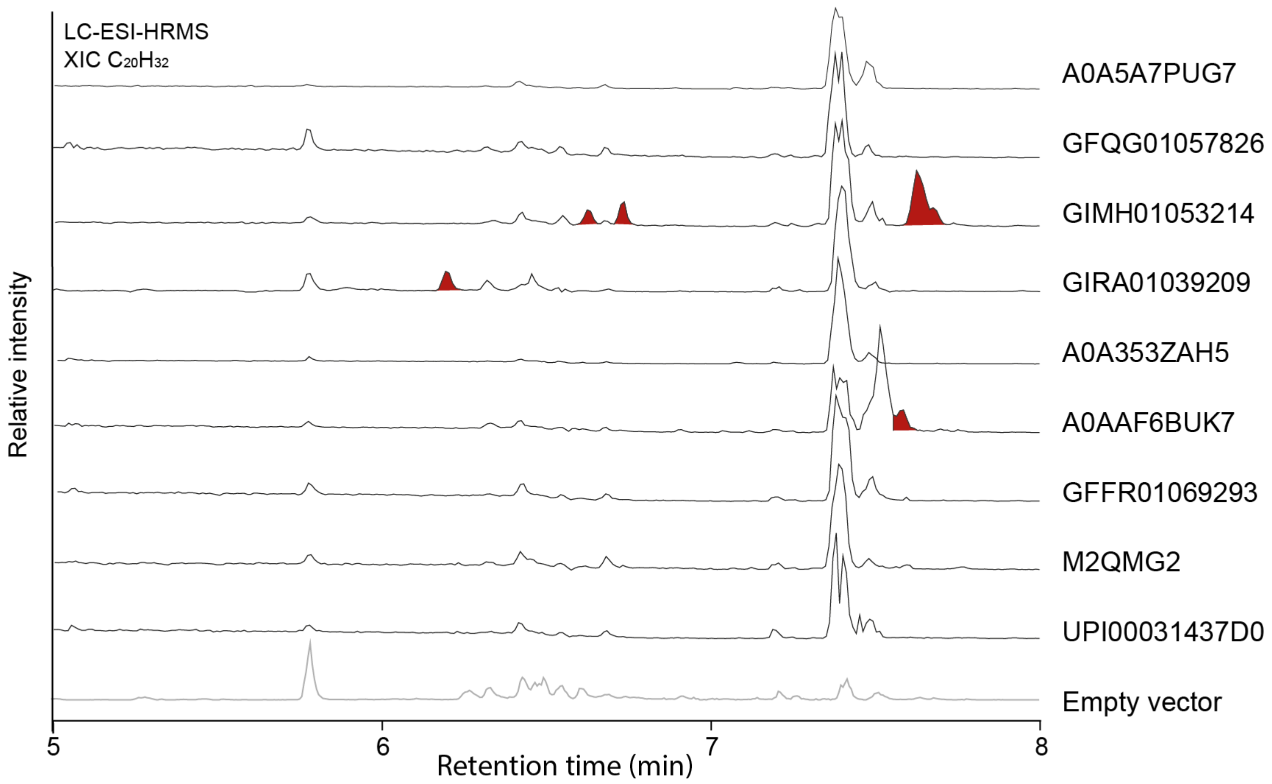


**Fig. S14 | LC-MS extracted ion chromatograms of the molecular mass of $[C_{20}H_{32}+H]^+$ ions at *m/z* 273.2577 (+/- 5 ppm).** Results of yeast expressions of proteins shown in **Table S1** (protein accession IDs shown on the right side). Detected diterpene ions are highlighted in red.

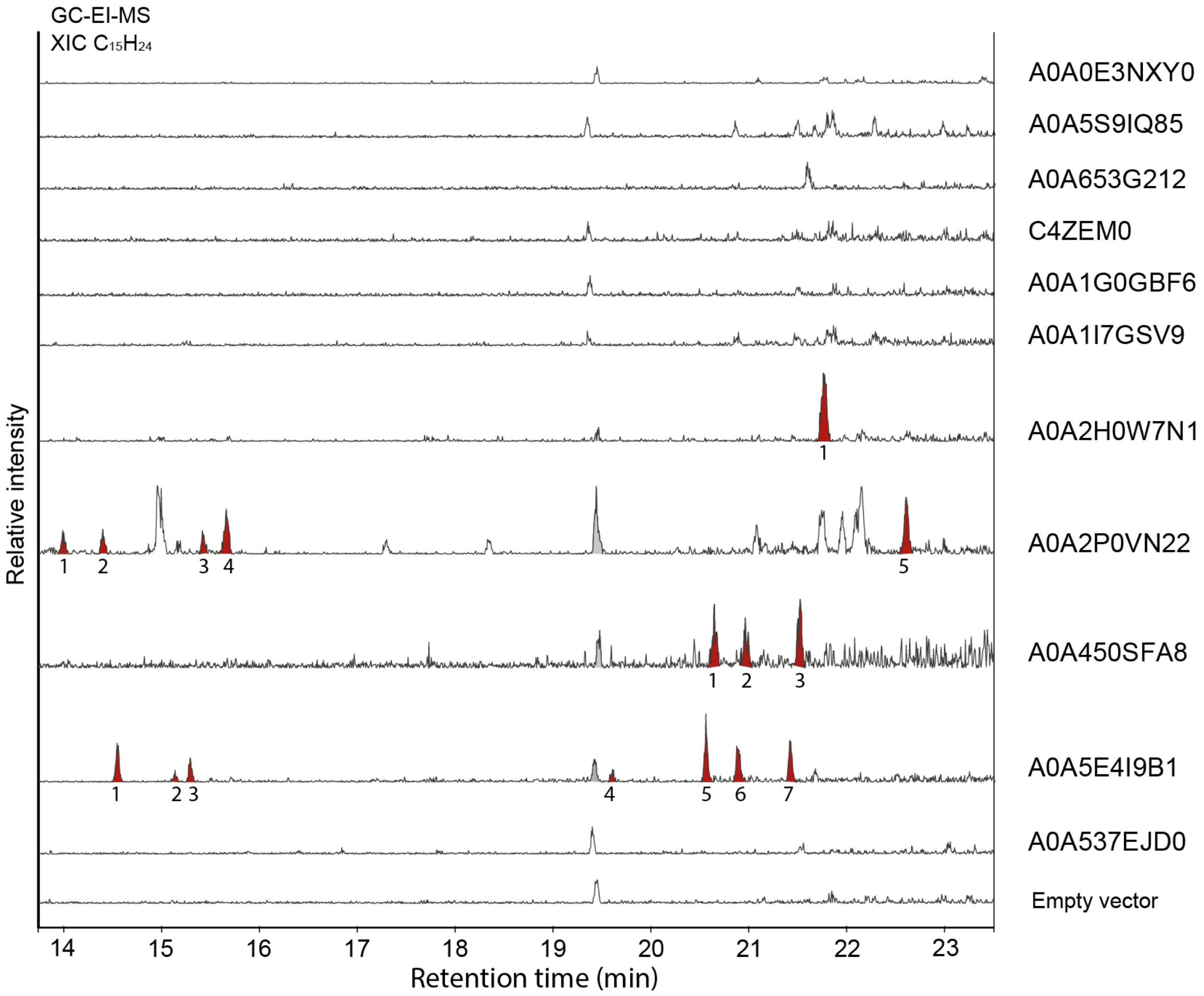


**Fig. S15 | GC-MS extracted ion chromatograms of the molecular mass of $C_{15}H_{24}$.** Results of yeast expressions of proteins shown in **Table 1** (protein accession IDs shown on the right side). Detected sesquiterpene ions are highlighted in red.

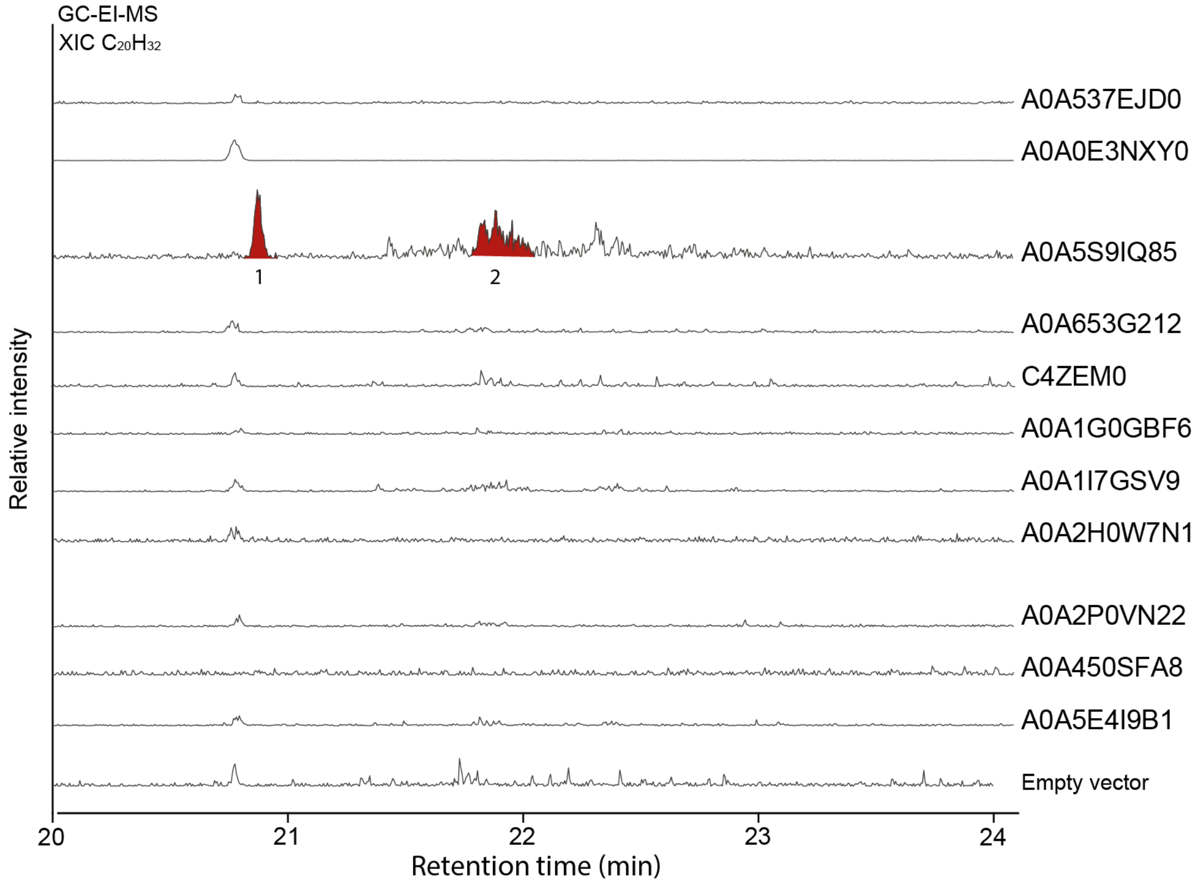


**Fig. S16 | GC-MS extracted ion chromatograms of the molecular mass of $C_{20}H_{32}$.** Results of yeast expressions of proteins shown in **Table 1** (protein accession IDs shown on the right side). Detected diterpene ions are highlighted in red.

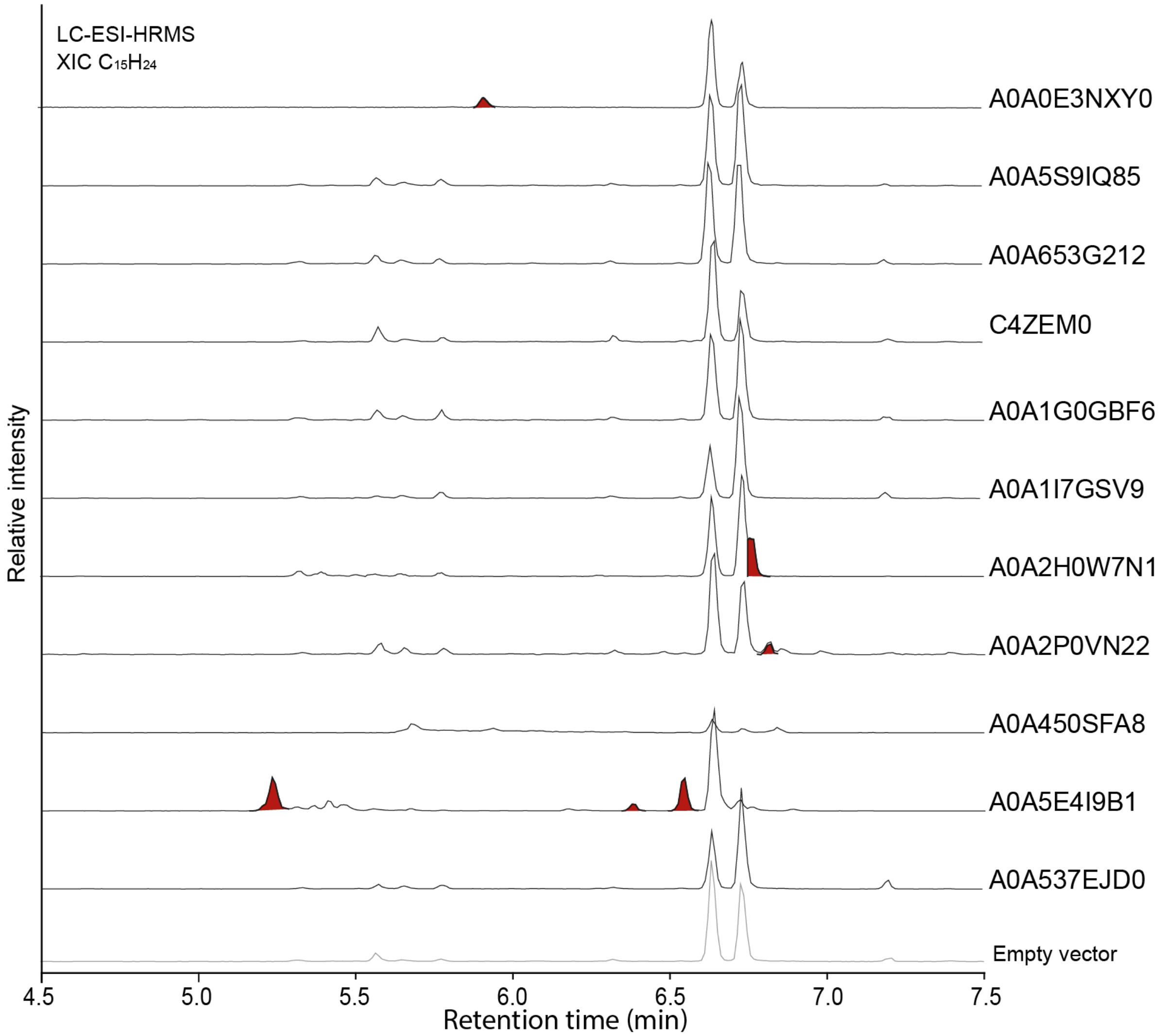


**Fig. S17 | LC-MS extracted ion chromatograms of the molecular mass of $[C_{15}H_{24}+H]^+$ ions at *m/z* 205.1951 (+/- 5 ppm).** Results of yeast expressions of proteins shown in **Table 1** (protein accession IDs shown on the right side). Detected sesquiterpene ions are highlighted in red.

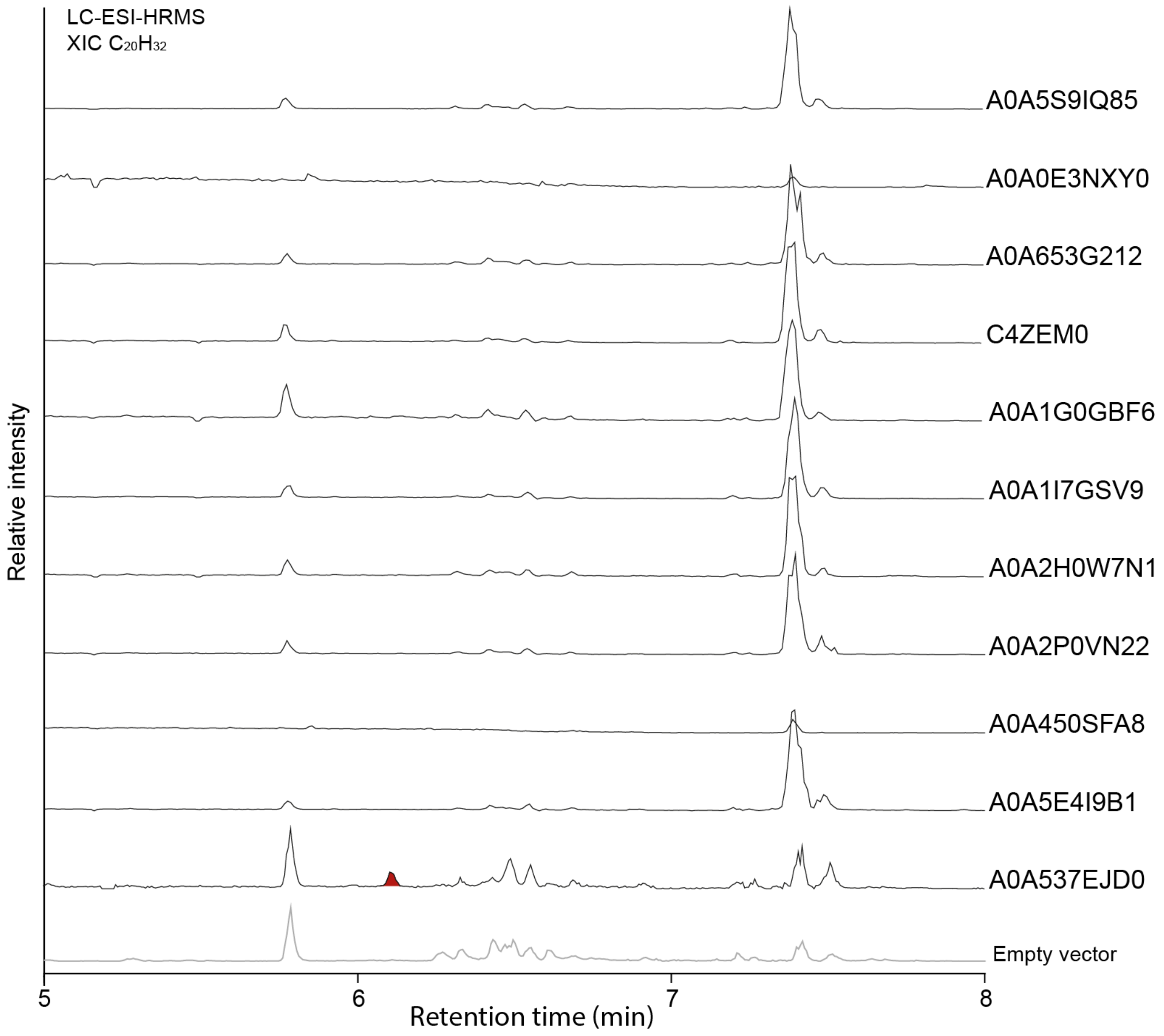


**Fig. S18 | LC-MS extracted ion chromatograms of the molecular mass of $[C_{20}H_{32}+H]^+$ ions at *m/z* 273.2577 (+/- 5 ppm).** Results of yeast expressions of proteins shown in **Table 1** (protein accession IDs shown on the right side). Detected diterpene ions are highlighted in red.

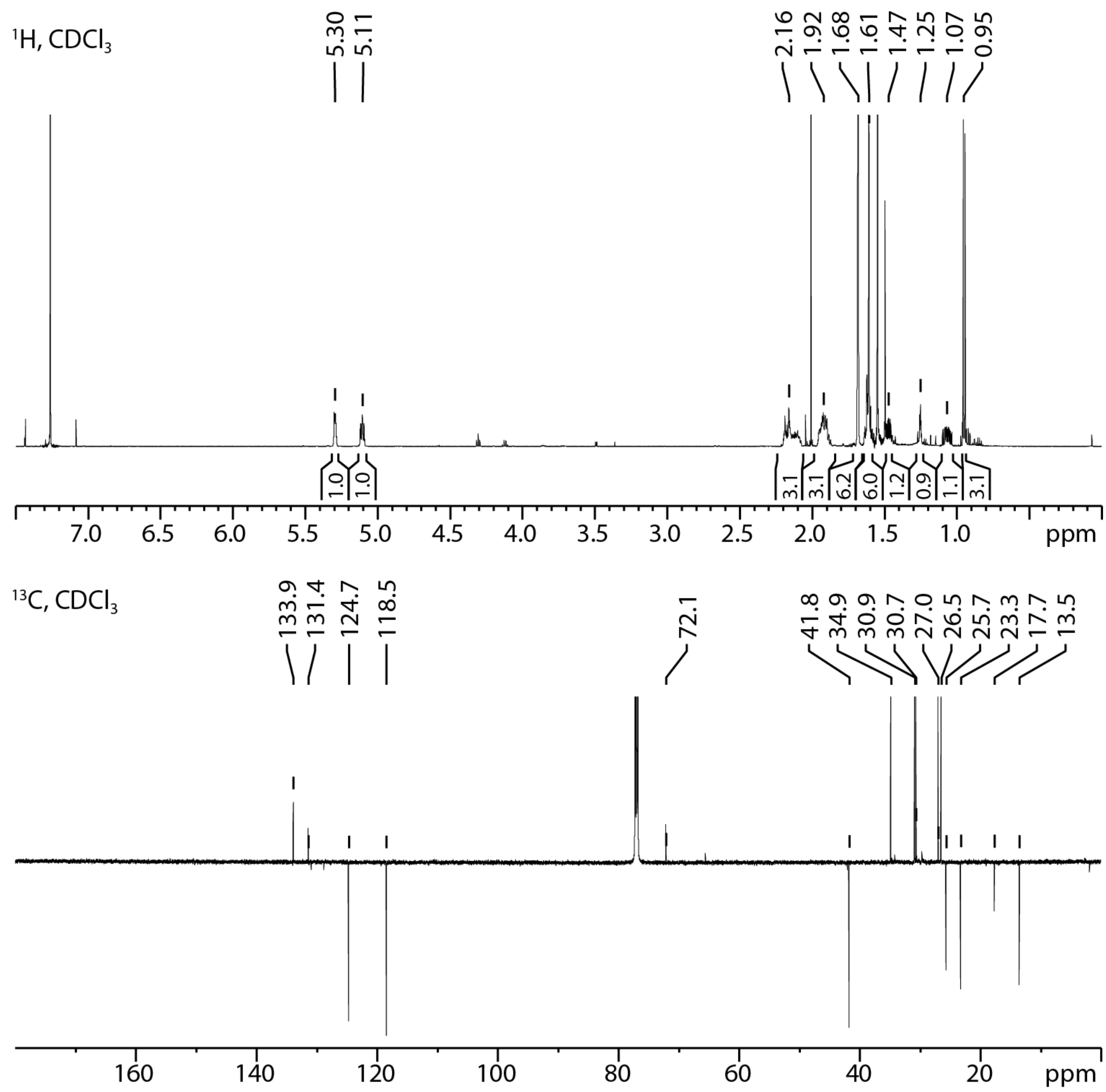


**Fig. S19 |** NMR spectra of isolated terpene **12** from the cultivation of yeast strain JWY501 expressing the archaeal enzyme A0A5E4I9B1.

**SMILES: CC1=CCC(C(C)CC/C=C(C)/C)(O)CC1**
**INCHIKEY:WTVHAMTYZJGJLJ-UHFFFAOYSA-N**

**δH**: 5.30 (1H, m), 5.11 (1H, m), 2.16 (3H, m), 1.92 (3H, m), 1.68 (6H, m), 1.61 (6H, m),1.47 (1H, m), 1.25 (1H, m), 1.07 (1H, m), 0.95 (3H, d).

**δC**: 133.9 (C), 131.4 (C), 124.7 (CH), 118.5 (CH), 72.1 (C), 41.8 (C), 34.9 (CH), 30.9 (CH2), 30.7 (CH2), 27.0 (CH2), 26.5 (CH2), 25.7 (CH3), 23.3 (CH3), 17.7 (CH3), 13.5 (CH3).

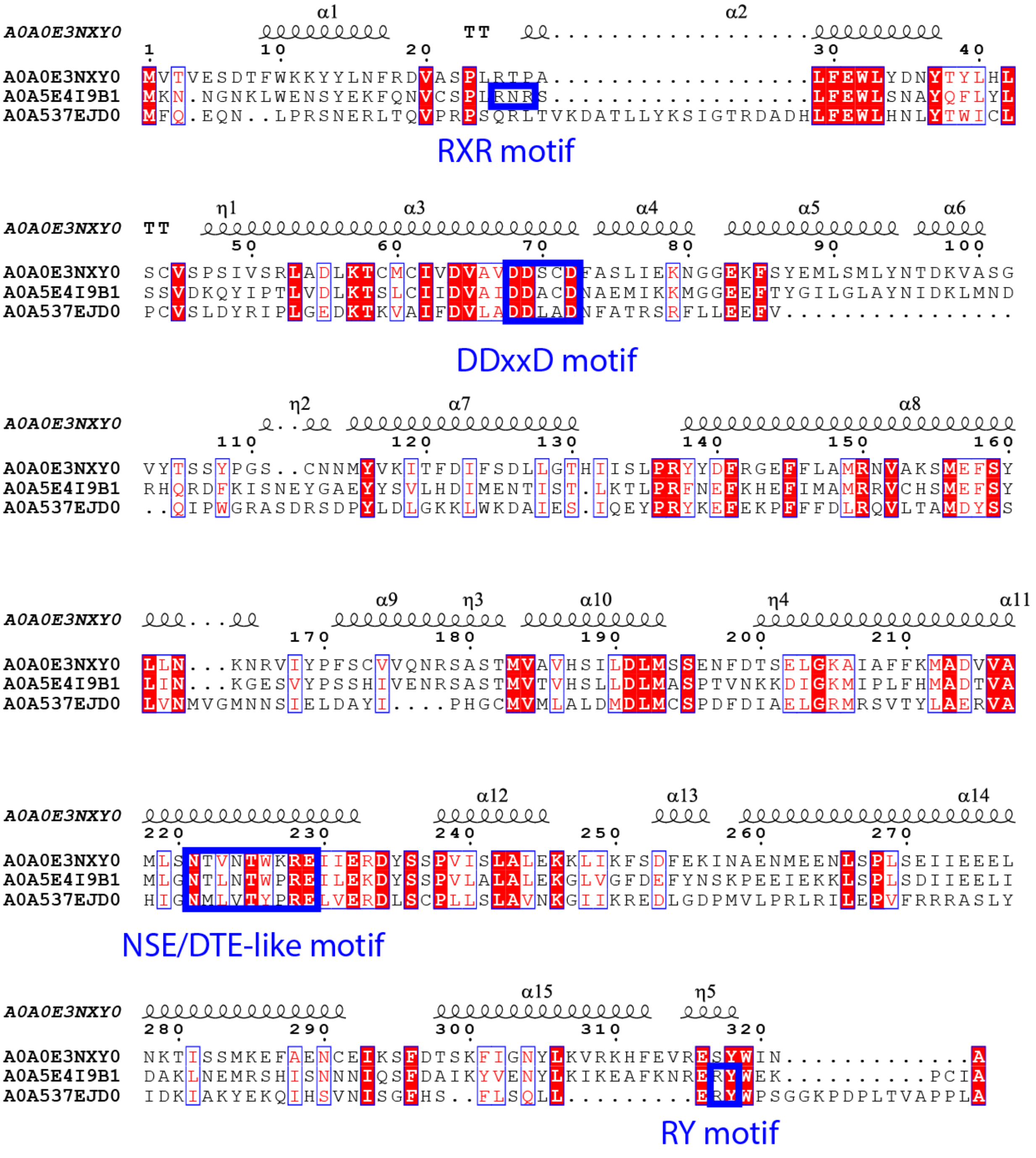


**Fig. S20 | Multiple sequence alignment of the three newly identified archaeal TPSs**. A0A5E4I9B1 and A0A0E3NXY0 were both identified as sesquiterpene synthases, while A0A537EJD0 is a diterpene synthase. Predicted secondary structure is shown for A0A0E3NXY0. Of the typical terpene synthase motifs, the aspartate rich motif DDXXD, typical for class I terpene synthases, is conserved across the three enzymes. A conserved sequence similar to the NSE/DTE motif can also be identified.

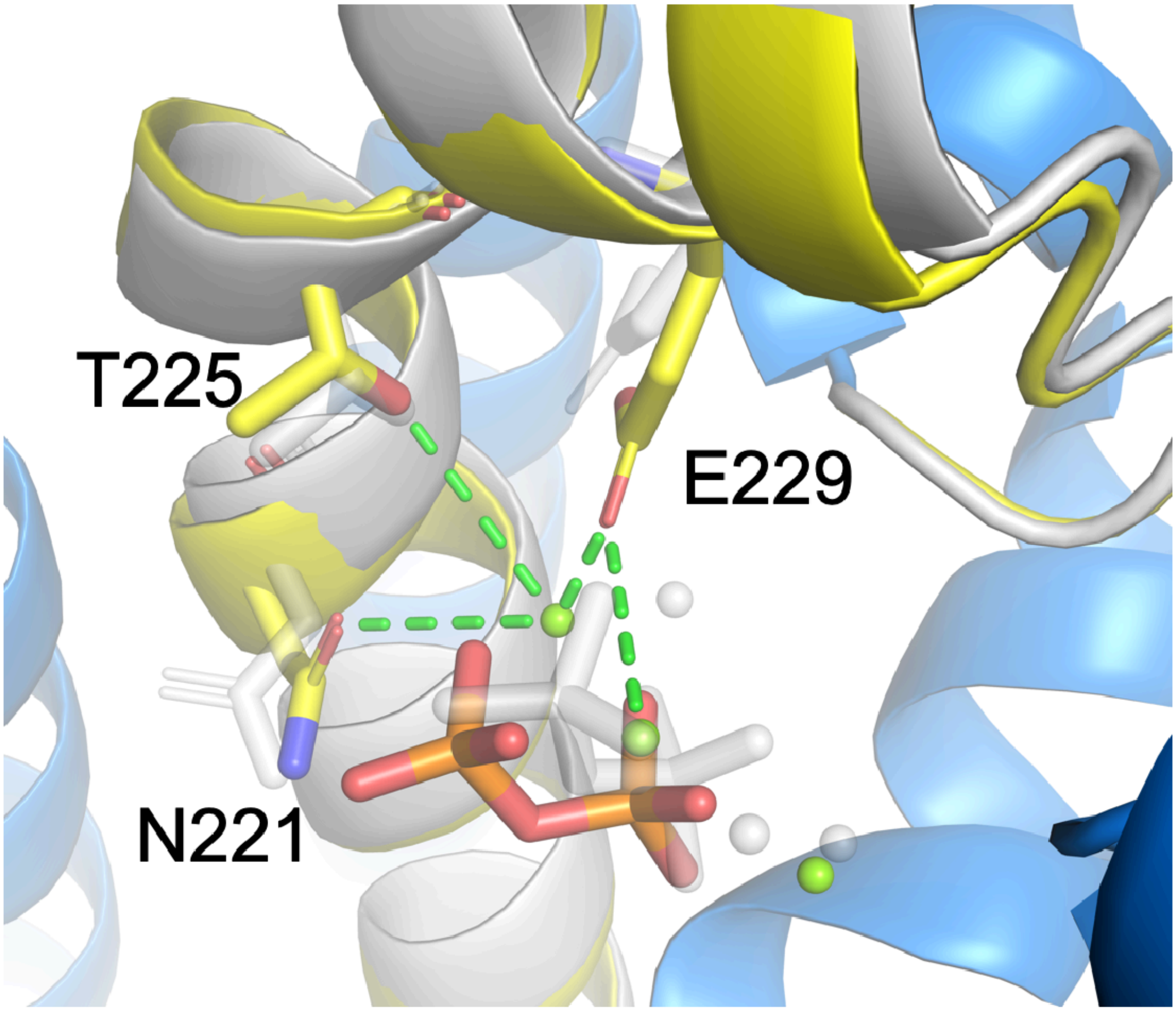


**Fig. S21 | The structure of the archaeal TPS A0A5E4I9B1 active site as predicted by AlphaFold**. The initial AlphaFold 3 model of A0A5E4I9B1 (grey) and activated model obtained from restrained energy minimization of the initial AlphaFold 3 model, which enforces TPS-specific interactions of conserved amino acid motifs (e.g., $^{221}$**N**TLN**T**WPR**E**) with the diphosphate and $Mg^{2+}$ cluster (yellow carbon color code). Blue color represents the region fixed during minimization.

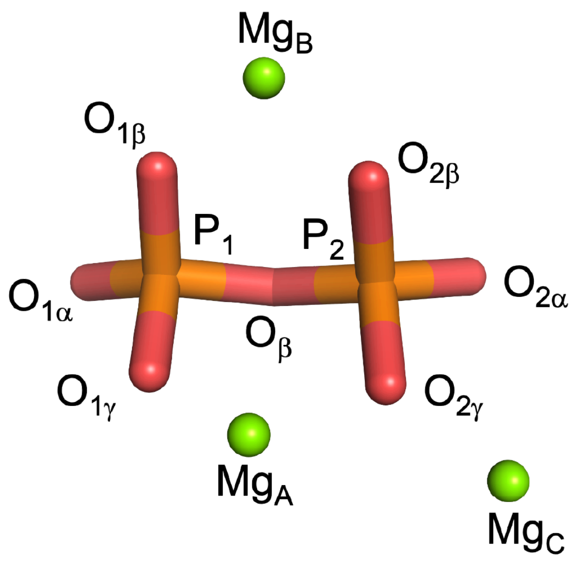


**Fig. S22 | Labeling of the diphosphate atoms and $Mg^{2+}$ ions in the TPS structure.**

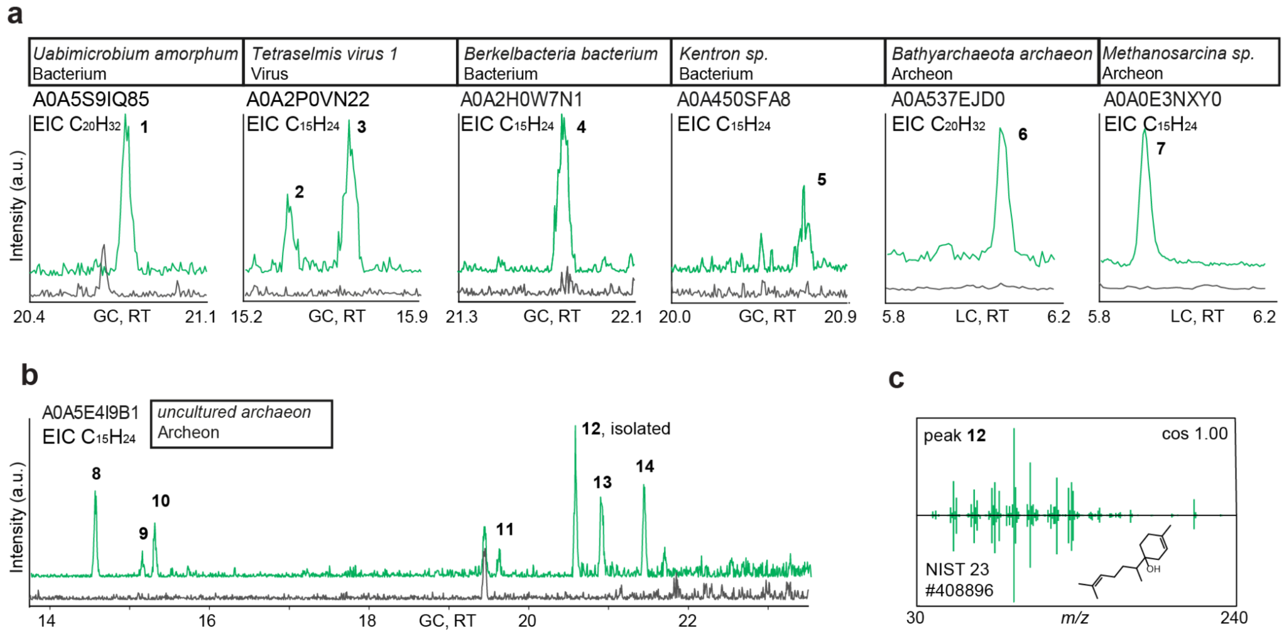


**Fig. S23 | Selected results of the experimental validation of EnzymeExplorer predictions for sequences that lack InterPro signatures**[3]**. a**, GC-MS and LC-MS extracted ion chromatograms (EICs) for sesquiterpene ($C_{15}H_{24}$) and diterpene ($C_{20}H_{32}$) molecular ions detected in the culture fluid of yeast strain JWY501 transformed with plasmids containing selected indicated enzymes (green). Empty vector control in JWY501 is shown in black. **b**, A GC-MS extracted ion chromatogram for sesquiterpene ($C_{15}H_{24}$) molecular ion detected in the culture fluid of yeast strain JWY501 transformed with plasmids containing selected indicated enzymes (green). Empty vector control in JWY501 is shown in black. **c**, The GC-MS fragmentation spectrum of peak 12 (top) shown as a mirror plot against the closest NIST23 EI library match (bottom). Matching peaks are in green; non-matching peaks in grey; cosine similarity score indicated (cos).

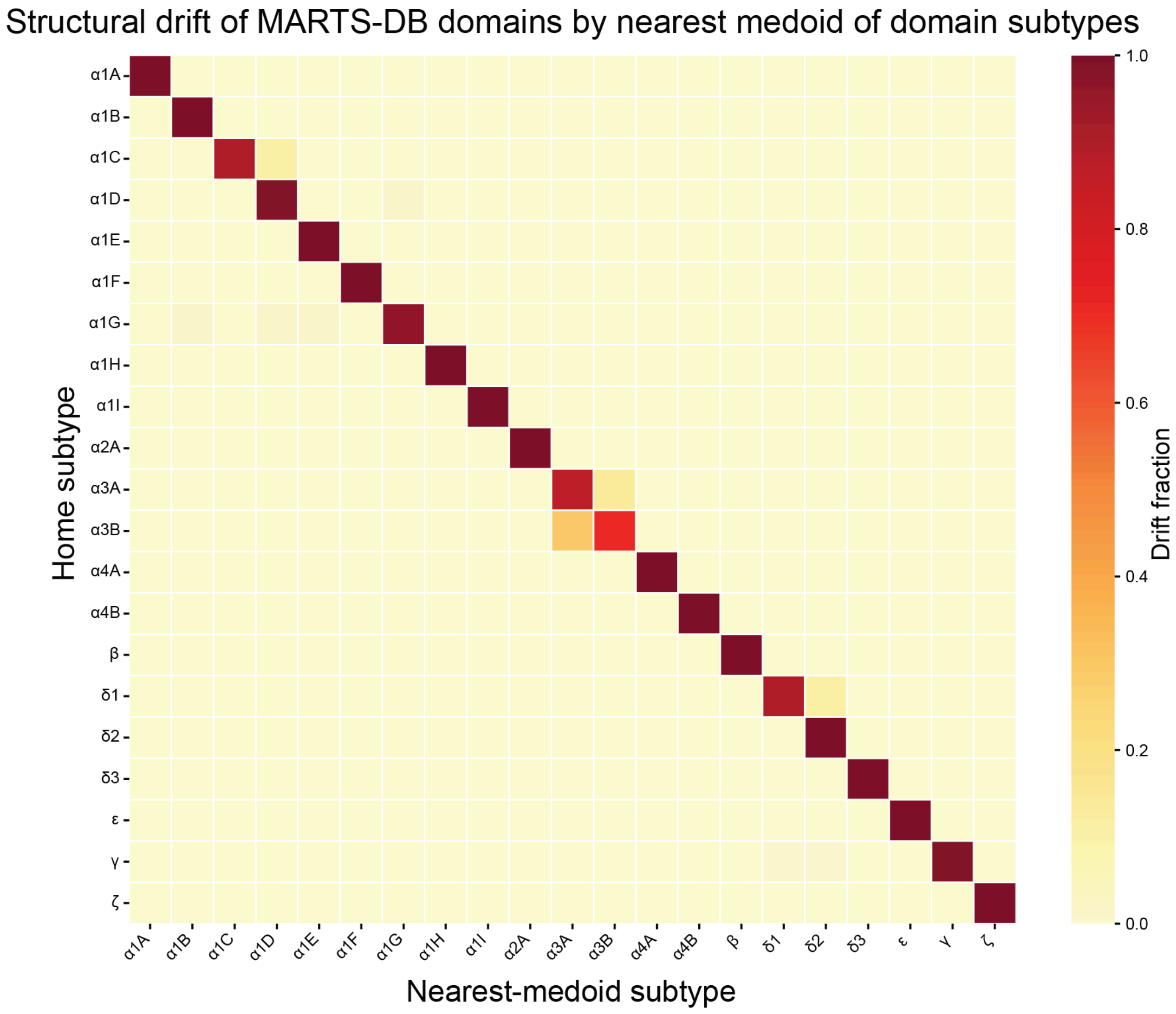


**Fig. S24 | Heatmap of structural drift across the determined domain subtype clusters in MARTS-DB.** Heatmap shows the fraction of transitional-domains from each domain subtype to another. A domain is flagged as transitional if TM-score to its home subtype's medoid is within 0.05 of the closest rival subtype's medoid. Domain subtypes are showing a consistent stability except α1C, α3A, α3B, and δ1, which have a tendency to drift to their sibling subtypes.

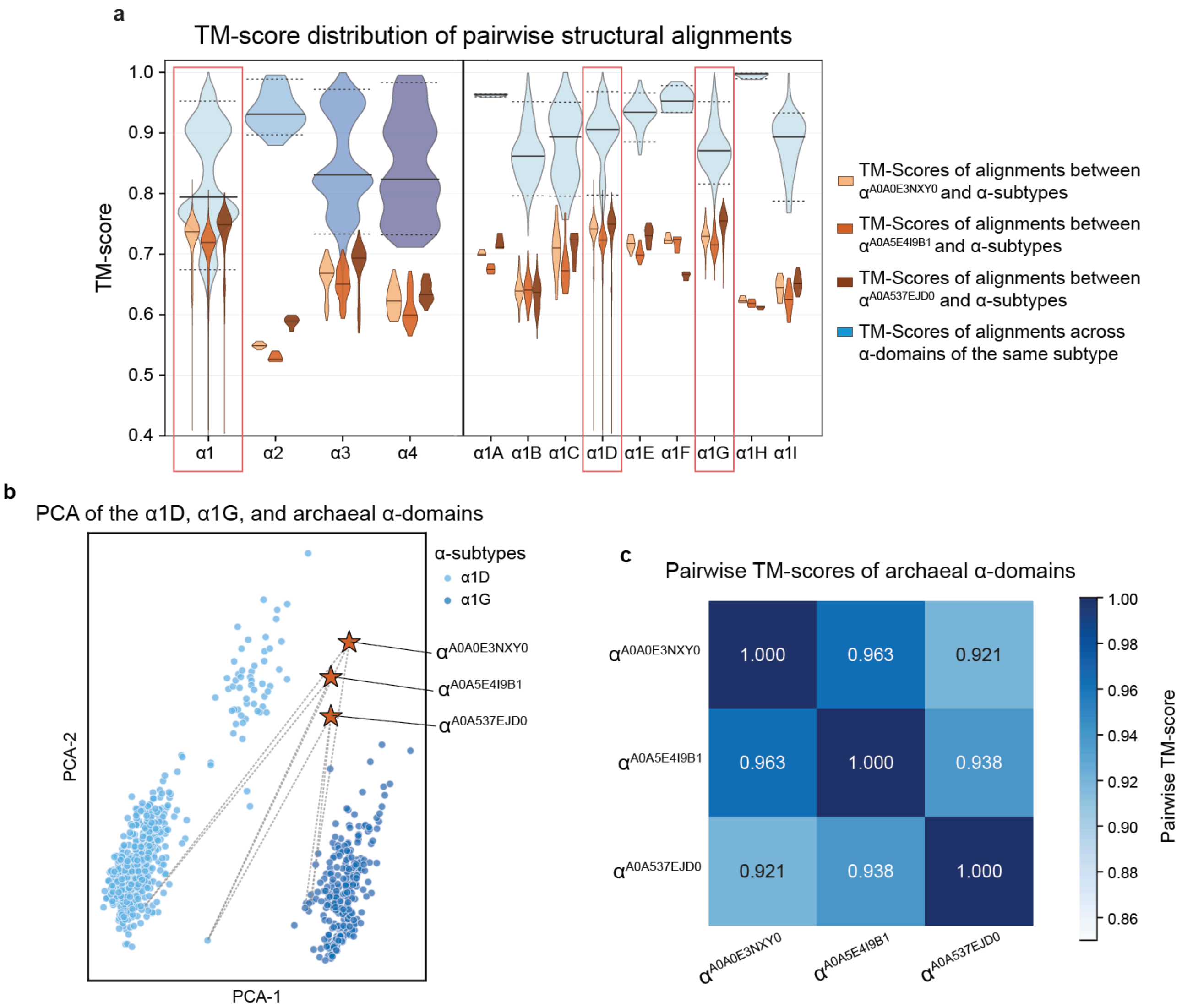


**Fig. S25 | Structural analysis of archaeal α-domains. a**, Structural similarity of archaeal α-domains to MARTS-DB α-subtypes. The violins show distribution of TM-scores from pairwise alignments between archaeal α-domains and MARTS-DB α-domain subtypes (brown), and intra-pairwise alignments across MARTS-DB α-domain subtypes (blue). Horizontal dashed lines represent the lower/upper 5% quartiles, and the solid lines represent the median of the distributions. The red rectangles show the closest domain subtypes to the archaeal domains. **b**, Combined PCA of α1D and α1G domain subtypes (the first two closest domain subtypes to archaeal α-domains) and archaeal α-domains. The archaeal α-domains are illustrated by stars, and the dashed lines connect them to the top three structurally closest α-domains. **c**, Heatmap of pairwise TM-scores of archaeal α-domains. Archaeal α-domains are structurally highly similar.

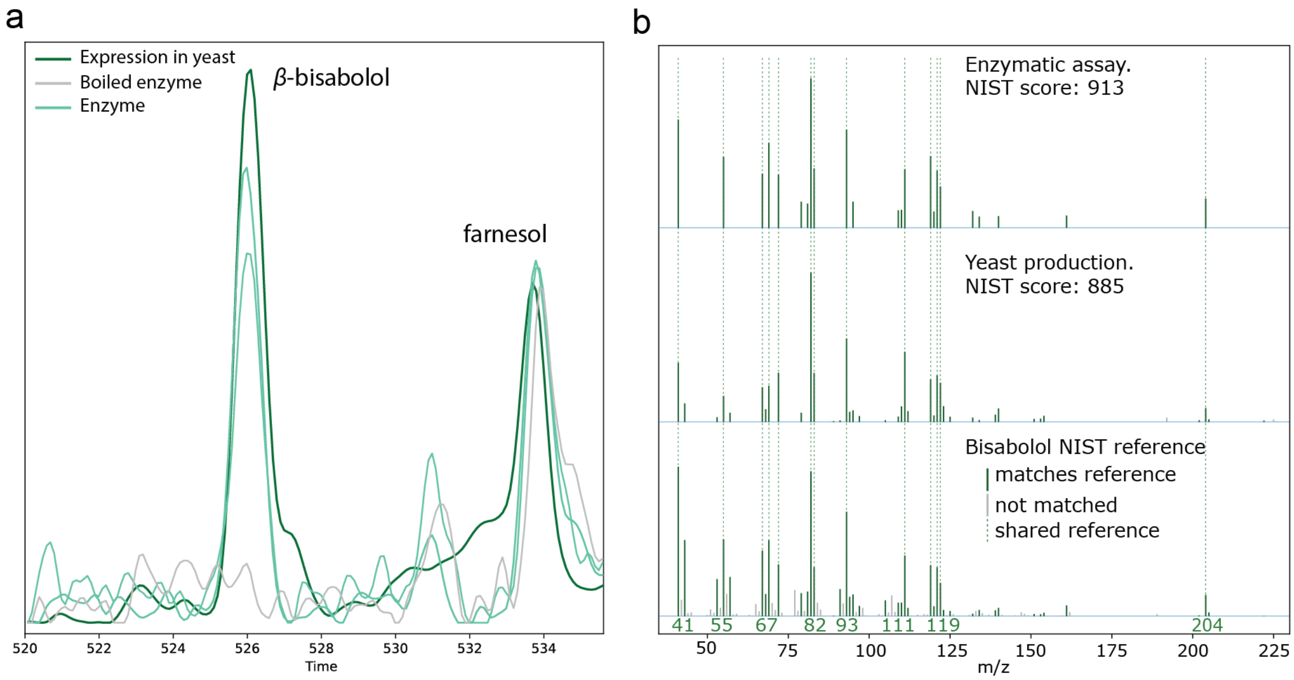


**Fig. S26 | *In vitro* formation of β-bisabolol by A0A5E4I9B1. a**, Overlaid GC-MS chromatograms of pentane extracts from reactions containing purified A0A5E4I9B1 (enzyme), heat-inactivated A0A5E4I9B1 (boiled enzyme) and a β-bisabolol-producing yeast strain used as a positive control. Peaks assigned to β-bisabolol and farnesol are indicated. **b**, Electron-ionization mass spectra of the β-bisabolol peak detected in the enzymatic reaction (top) and yeast reference extract (middle), compared with the National Institute of Standards and Technology β-bisabolol library spectrum (bottom). Green lines indicate ions matching the library spectrum, grey lines indicate unmatched ions and dashed lines indicate ions shared with the reference spectrum.

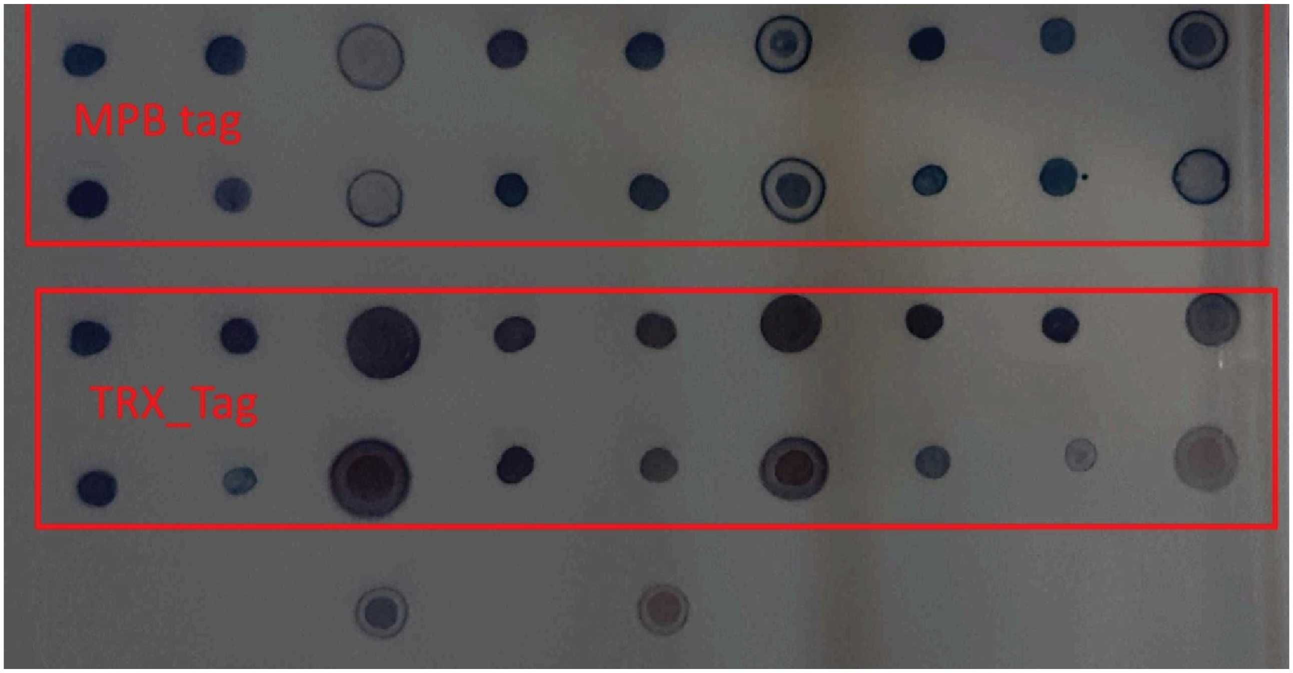


**Fig. S27 | Dot-blot analysis of MBP- and Trx-tagged recombinant proteins.** Protein samples obtained under the indicated expression or purification conditions were spotted onto membranes in duplicate. The upper panel shows samples containing the maltose-binding protein (MBP) tag, and the lower panel shows samples containing the thioredoxin (Trx) tag. Dot-blot signals provide a qualitative assessment of the relative abundance of the corresponding tagged proteins.

### Supplementary Tables

**Table S1**. List of candidate TPS sequences expressed in yeast. Sequences were selected based on phylogenetic analysis of mined TPS-like sequences (**Fig. S10a**). The predicted probability scores correspond to the calibrated prediction scores. All experimentally verified candidates were predicted as TPS with an approx. 100% probability estimate (half-up rounding with six decimal places). The predicted substrate column shows the substrate predictions for FPP and GGPP. EnzymeExplorer was able to predict the correct substrate for four out of six experimentally verified candidates.

| **Accession ID** | **Origin** | **Species of origin** | **InterPro annotations**[3] | **Predicted substrate** | **Observed activity** | **TPS prediction score (%)** | **FPP prediction score (%)** | **GGPP prediction score (%)** |
|---|---|---|---|---|---|---|---|---|
| A0A5A7PUG7 | Plant | *Striga asiatica* | PF01397, PF03936 | - | - | 100.00 | 17.48 | 2.73 |
| GFQG01057826 | Plant | *Amaranthus tricolor* | PF01397, PF03936 | - | - | 100.00 | 2.19 | 22.77 |
| GIMH01053214 | Plant | *Selaginella wallacei* | PF01397, PF03936 | GGPP | **Di** | 100.00 | 0.20 | 94.01 |
| GIRA01039209 | Plant | *Beta vulgaris* | PF01397, PF03936 | - | **Di** | 100.00 | 3.74 | 16.32 |
| A0A353ZAH5 | Bacterium | *Planctomycetaceae bacterium* | SSF48239 | - | | 0.34 | 0.57 | 0.50 |
| A0AAF6BUK7 | Plant | *Marchantia polymorpha* | PF01397, PF03936 | GGPP | **Di** | 100.00 | 0.15 | 95.91 |
| GFFR01069293 | Plant | *Camellia sinensis* | PF01397, PF03936 | FPP | **Sesqui** | 100.00 | 96.75 | 0.07 |
| M2QMG2 | Fungus | *Gelatoporia subvermispora* | PF19086 | FPP | **Sesqui** | 100.00 | 88.59 | 6.23 |
| UPI00031437D0 | Bacterium | *Pseudomonas agarici* | PF19086 | GGPP | **Sesqui** | 100.00 | 16.19 | 59.54 |

**Table S2.** Conserved interactions in the polar region of the active site in the refined AlphaFold 3 model of A0A5E4I9B1.

| **Atom 1** | **Atom 2** | **Distance (Å)** |
|---|---|---|
| Asp63(OD1) | $W_{cat}$ | 3.3 |
| PP(O2a) | $W_{cat}$ | 2.9 |
| Asn221(OD1) | $MG_B$ | 3.9 |
| Glu229(OE1) | $MG_B$ | 1.9 |
| Glu229(OE1) | $MG_A$ | 4.0 |
| Thr225(OG1) | $MG_B$ | 4.0 |
| Asp68(OD2) | Arg24(NE) | 3.0 |
| Asp68(OD1) | Arg24(NH2) | 2.6 |
| Arg24(NH1) | PP(O1a) | 4.0 |
| Arg24(NH2) | PP(O1g) | 4.0 |
| Asp67(OD2) | $MG_A$ | 3.1 |
| Asp67(OD1) | $MG_C$ | 1.8 |
| PP(O1g) | $MG_A$ | 1.8 |
| PP(O2g) | $MG_A$ | 1.9 |
| PP(O2g) | $MG_C$ | 1.9 |
| Asp71(OD2) | $MG_C$ | 2.0 |
| Asp71(OD2) | $MG_A$ | 3.6 |

**Table S3.** Top ten poses for the substrate FPP in A0A5E4I9B1 bound to either the O1*α* or the O2*α* oxygens. EnzyDock energies are total binding energies (kcal/mol).

| **Cluster** | **$E_{tot}$[a]** | **Diphosphate oxygen** |
|---|---|---|
| 1 | 0.27 | O1*α* |
| 2 | 0.62 | O1*α* |
| 3 | 0.94 | O1*α* |
| 4 | 14.36 | O1*α* |
| 5 | 16.36 | O1*α* |
| 6 | 16.61 | O1*α* |
| 7 | 22.84 | O1*α* |
| 8 | 27.96 | O1*α* |
| 9 | 30.22 | O1*α* |
| 10 | 32.39 | O1*α* |
| 1 | 14.50 | O2*α* |
| 2 | 20.38 | O2*α* |
| 3 | 22.11 | O2*α* |
| 4 | 29.37 | O2*α* |
| 5 | 31.07 | O2*α* |
| 6 | 35.78 | O2*α* |
| 7 | 39.13 | O2*α* |
| 8 | 73.88 | O2*α* |
| 9 | 80.10 | O2*α* |
| 10 | 89.00 | O2*α* |

[a] Total force field energy of the ligand.

**Table S4**. List of yeast strains used in this study.

| **Name** | **Genotype** | **Ref.** |
|---|---|---|
| JWY501 | MATα *leu2*–3112::*His3MX6*_P$_{GAL1}$-*ERG19*/P$_{GAL10}$-*ERG8*<br>*ura3–52::URA3*_P$_{GAL1}$-*mvaS(A110G)*/P$_{GAL10}$-<br>*mvaE(CO) his3Δ1::hphMX4*_P$_{GAL1}$-*ERG12*/P$_{GAL10}$-*IDI1*<br>*trp1–289::TRP1*_P$_{GAL1}$-*crtE(X.den)*/P$_{GAL10}$-*ERG20*<br>*yprcδ15::natMX*_P$_{GAL1}$-*crtE(opt)*/P$_{GAL10}$-*crtE*<br>(*ura3–52* prototrophy removed for use of Cas9 system) | Wong *et al.* [52] |
| JWY501_TP0234 | JWY501 *ura3-52::ura3::*P*tdh3*::A0A0E3NXY0::T*ssa1* | This study |
| JWY501_TP0235 | JWY501 ura3-52::ura3::P*tdh3*::A0A5S9IQ85::T*ssa1* | This study |
| JWY501_TP0236 | JWY501 *ura3-52::ura3::*P*tdh3*::A0A653G212::T*ssa1* | This study |
| JWY501_TP0239 | JWY501 ura3-52::ura3::P*tdh3*::A0A8I0Q8N9::T*ssa1* | This study |
| JWY501_TP0240 | JWY501 ura3-52::ura3::P*tdh3*::C4ZEM0::T*ssa1* | This study |
| JWY501_TP0242 | JWY501 ura3-52::ura3::P*tdh3*::A0A8J2KXK4::T*ssa1* | This study |
| JWY501_TP0243 | JWY501 *ura3-52::ura3::*P*tdh3*::D3BLT7::T*ssa1* | This study |
| JWY501_TP0244 | JWY501 ura3-52::ura3::P*tdh3*::A0A1G0GBF6::T*ssa1* | This study |
| JWY501_TP0245 | JWY501 ura3-52::ura3::P*tdh3*::A0A8J2P3A2::T*ssa1* | This study |
| JWY501_TP0246 | JWY501 ura3-52::ura3::P*tdh3*::F0ZD71::T*ssa1* | This study |
| JWY501_TP0247 | JWY501 ura3-52::ura3::P*tdh3*::A0A1I7GSV9::T*ssa1* | This study |
| JWY501_TP0250 | JWY501 ura3-52::ura3::P*tdh3*::A0A2H0W7N1::T*ssa1* | This study |
| JWY501_TP0251 | JWY501 ura3-52::ura3::P*tdh3*::A0A417ATN1::T*ssa1* | This study |
| JWY501_TP0253 | JWY501 ura3-52::ura3::P*tdh3*::A0A2P0VN22::T*ssa1* | This study |
| JWY501_TP0254 | JWY501 ura3-52::ura3::P*tdh3*::A0A450SFA8::T*ssa1* | This study |
| JWY501_TP0255 | JWY501 ura3-52::ura3::P*tdh3*::A0A5E4I9B1::T*ssa1* | This study |
| JWY501_TP0256 | JWY501 ura3-52::ura3::P*tdh3*::A0A537EJD0::T*ssa1* | This study |

**Table S5**. List of plasmids used in this study.

| Plasmid name | Plasmid type | Backbone & selection marker | Description | Ref. |
|---|---|---|---|---|
| pYTK001 | Backbone | pYTK001, Cm | Storage plasmid | Lee et al. [103] |
| pYTK002 | Part type 1 | pYTK001, Cm | ConLS | Lee et al. [103] |
| pYTK009 | Part type 2 | pYTK001, Cm | P*tdh3* | Lee et al. [103] |
| pYTK030 | Part type 2 | pYTK001, Cm | P*gal1* | Lee et al. [103] |
| pYTK047 | Part type 234 | pYTK001, Cm | GFP dropout | Lee et al. [103] |
| pYTK052 | Part type 4 | pYTK001, Cm | T*ssa1* | Lee et al. [103] |
| pYTK067 | Part type 5 | pYTK001, Cm | ConR1 | Lee et al. [103] |
| pYTK074 | Part type 6 | pYTK001, Cm | URA3 | Lee et al. [103] |
| pYTK082 | Part type 7 | pYTK001, m | 2µ origin of replication | Lee et al. [103] |
| pYTK083 | Part type 8 | pYTK083, Amp | Backbone, bacterial module | Lee et al. [103] |
| pYTK096 | Backbone | pYTK096, Amp | Ura3 targeting integration vector | Lee et al. [103] |
| pESC-Ura | Backbone | Backbone, Amp | Ura3 resistance marker | Agilent, cat. #217454 |
| pTP0027 | Backbone | pYTK083, Amp | Transient expression, Ura3 prototrophy | This study |
| pECO_Trx | Backbone | Kan | GoldenGate compatible, Protein expression, KanR, IPTG | This study |
| pTP0098 | Part type 3 | pYTK001, Cm | A0A5A7PUG7 | This study |
| pTP0099 | Part type 3 | pYTK001, Cm | GFQG01057826 | This study |
| pTP0100 | Part type 3 | pYTK001, Cm | GIMH01053214 | This study |
| pTP0101 | Part type 3 | pYTK001, Cm | GIRA01039209 | This study |
| pTP0102 | Part type 3 | pYTK001, Cm | A0A353ZAH5 | This study |
| pTP0103 | Part type 3 | pYTK001, Cm | A0AAF6BUK7 | This study |
| pTP0104 | Part type 3 | pYTK001, Cm | GFFR01069293 | This study |
| pTP0105 | Part type 3 | pYTK001, Cm | M2QMG2 | This study |
| pTP0106 | Part type 3 | pYTK001, Cm | UPI00031437D0 | This study |
| pTP0195 | Transient expression | pTP0027, Amp | P*thd3,* A0A5A7PUG7, T*ssa1* | This study |
| pTP0196 | Transient expression | pTP0027, Amp | P*thd3,* GFQG01057826, T*ssa1* | This study |
| pTP0197 | Transient expression | pTP0027, Amp | P*thd3,* GIMH01053214, T*ssa1* | This study |
| pTP0198 | Transient expression | pTP0027, Amp | P*thd3,* GIRA01039209, T*ssa1* | This study |
| pTP0199 | Transient expression | pTP0027, Amp | P*thd3,* A0A353ZAH5, T*ssa1* | This study |
| pTP0200 | Transient expression | pTP0027, Amp | P*thd3,* A0AAF6BUK7, T*ssa1* | This study |
| pTP0201 | Transient expression | pTP0027, Amp | P*thd3,* GFFR01069293, T*ssa1* | This study |
| pTP0202 | Transient expression | pTP0027, Amp | P*thd3,* M2QMG2, T*ssa1* | This study |
| pTP0203 | Transient expression | pTP0027, Amp | P*thd3,* UPI00031437D0, T*ssa1* | This study |
| pTP0209 | Part type 3 | pYTK001, Cm | A0A0E3NXY0 | This study |
| pTP0210 | Part type 3 | pYTK001, Cm | A0A5S9IQ85 | This study |

| | | | | |
|---|---|---|---|---|
| pTP0211 | Part type 3 | pYTK001, Cm | A0A653G212 | This study |
| pTP0214 | Part type 3 | pYTK001, Cm | A0A8I0Q8N9 | This study |
| pTP0215 | Part type 3 | pYTK001, Cm | C4ZEM0 | This study |
| pTP0217 | Part type 3 | pYTK001, Cm | A0A8J2KXK4 | This study |
| pTP0218 | Part type 3 | pYTK001, Cm | D3BLT7 | This study |
| pTP0219 | Part type 3 | pYTK001, Cm | A0A1G0GBF6 | This study |
| pTP0220 | Part type 3 | pYTK001, Cm | A0A8J2P3A2 | This study |
| pTP0221 | Part type 3 | pYTK001, Cm | F0ZD71 | This study |
| pTP0222 | Part type 3 | pYTK001, Cm | A0A1I7GSV9 | This study |
| pTP0225 | Part type 3 | pYTK001, Cm | A0A2H0W7N1 | This study |
| pTP0226 | Part type 3 | pYTK001, Cm | A0A417ATN1 | This study |
| pTP0228 | Part type 3 | pYTK001, Cm | A0A2P0VN22 | This study |
| pTP0229 | Part type 3 | pYTK001, Cm | A0A450SFA8 | This study |
| pTP0230 | Part type 3 | pYTK001, Cm | A0A5E4I9B1 | This study |
| pTP0231 | Part type 3 | pYTK001, Cm | A0A537EJD0 | This study |
| pTP0234 | Integration vector | pYTK096, amp | P*tdh3*, A0A0E3NXY0, T*ssa1* | This study |
| pTP0235 | Integration vector | pYTK096, amp | P*tdh3*, A0A5S9IQ85, T*ssa1* | This study |
| pTP0236 | Integration vector | pYTK096, amp | P*tdh3*, A0A653G212, T*ssa1* | This study |
| pTP0239 | Integration vector | pYTK096, amp | P*tdh3*, A0A8I0Q8N9, T*ssa1* | This study |
| pTP0240 | Integration vector | pYTK096, amp | P*tdh3*, C4ZEM0, T*ssa1* | This study |
| pTP0242 | Integration vector | pYTK096, amp | P*tdh3*, A0A8J2KXK4, T*ssa1* | This study |
| pTP0243 | Integration vector | pYTK096, amp | P*tdh3*, D3BLT7, T*ssa1* | This study |
| pTP0244 | Integration vector | pYTK096, amp | P*tdh3*, A0A1G0GBF6, T*ssa1* | This study |
| pTP0245 | Integration vector | pYTK096, amp | P*tdh3*, A0A8J2P3A2, T*ssa1* | This study |
| pTP0246 | Integration vector | pYTK096, amp | P*tdh3*, F0ZD71, T*ssa1* | This study |
| pTP0247 | Integration vector | pYTK096, amp | P*tdh3*, A0A1I7GSV9, T*ssa1* | This study |
| pTP0250 | Integration vector | pYTK096, amp | P*tdh3*, A0A2H0W7N1, T*ssa1* | This study |
| pTP0251 | Integration vector | pYTK096, amp | P*tdh3*, A0A417ATN1, T*ssa1* | This study |
| pTP0253 | Integration vector | pYTK096, amp | P*tdh3*, A0A2P0VN22, T*ssa1* | This study |
| pTP0254 | Integration vector | pYTK096, amp | P*tdh3*, A0A450SFA8, T*ssa1* | This study |
| pTP0255 | Integration vector | pYTK096, amp | P*tdh3*, A0A5E4I9B1, T*ssa1* | This study |
| pTP0255µ | Transient expression | pTP0027, Amp | P*gal1*, A0A5E4I9B1, T*ssa1* | This study |
| pTP0255_HisTrx | Protein expression | pECO_Trx (Custom backbone) | His-Trx-tagged A0A5E4I9B1 expression | This study |
| pTP0256 | Integration vector | pYTK096, amp | P*tdh3*, A0A537EJD0, T*ssa1* | This study |

**Table S6.** Comparison of baseline methods and EnzymeExplorer for TPS activity prediction of candidate TPS sequences selected from the dark putatives (**Figs. S10b-c)**. For each candidate, the TPS prediction scores (%) of the methods are given. Details regarding the baselines are described in **Performance evaluation of predictive models**. The rows of experimentally verified sequences are written in bold characters. Even though CLEAN was able to predict TPS activity for each experimentally verified TPS candidate, the 5-fold cross-validation indicated that CLEAN's average precision is significantly lower than BLASTp, Foldseek, and EnzymeExplorer (**Figs. 2b-e, S7**). Further analysis regarding the 5-fold cross-validation showed that CLEAN's 27/1270 positive predictions were false, which were correctly rejected by the three top-performing methods (e.g. UniProt ID Q55EU7, P15482, P58334, O15544, O43008, A0A3G9GNJ4). The lack of precision makes CLEAN unfavourable for screening large sequence databases for "dark putative" TPSs and selection of experimental candidates.

| | CLEAN | Foldseek | BLASTp | pHMM | Pfam | SUPFAM | EnzymeExplorer |
|---|---|---|---|---|---|---|---|
| **A0A450SFA8** | **100.00** | **96.00** | **0.00** | **0.00** | **100.00** | **0.00** | **99.99** |
| **A0A5S9IQ85** | **100.00** | **100.00** | **0.00** | **0.00** | **0.00** | **0.00** | **99.99** |
| **A0A0E3NXY0** | **100.00** | **99.00** | **0.00** | **0.00** | **0.00** | **0.00** | **99.99** |
| **A0A2H0W7N1** | **100.00** | **95.00** | **0.00** | **0.00** | **0.00** | **0.00** | **100.00** |
| **A0A5E4I9B1** | **100.00** | **99.00** | **0.00** | **0.00** | **100.00** | **0.00** | **99.99** |
| A0A1G0GBF6 | 100.00 | 100.00 | 0.00 | 0.00 | 0.00 | 0.00 | 100.00 |
| **A0A537EJD0** | **100.00** | **100.00** | **0.00** | **0.00** | **0.00** | **0.00** | **99.99** |
| A0A653G212 | 100.00 | 100.00 | 0.00 | 0.00 | 0.00 | 0.00 | 99.88 |
| C4ZEM0 | 80.00 | 96.00 | 0.00 | 0.00 | 0.00 | 0.00 | 99.56 |
| A0A1I7GSV9 | 100.00 | 100.00 | 0.00 | 0.00 | 0.00 | 0.00 | 98.74 |
| **A0A2P0VN22** | **100.00** | **0.00** | **0.00** | **0.00** | **0.00** | **0.00** | **95.93** |

**Table S7.** EnzymeExplorer precision and recall at different thresholds of TPS detection calibrated scores.

| Threshold (%) | #positive predictions | Precision (%) | Recall (%) |
|---|---|---|---|
| 50 | 1,263 | 99.05 | 99.60 |
| 80 | 1,247 | 99.76 | 99.04 |
| 90 | 1,243 | 99.76 | 98.73 |
| 95 | 1,240 | 99.92 | 98.65 |
| 99 | 1,236 | 100.00 | 98.41 |
| 99.95 | 1,215 | 100.00 | 96.74 |

**Table S8.** Calibration evaluation by log-loss and Brier scores of raw and calibrated prediction scores by prediction task. For both metrics, lower scores represent better probability estimates.

| Prediction task | log-loss (%) | | Brier score (%) | |
|---|---|---|---|---|
| | raw | calibrated | raw | calibrated |
| TPS | 14.44 | 0.47 | 2.19 | 0.12 |
| GPP | 7.40 | 4.24 | 1.68 | 1.29 |
| FPP | 9.90 | 4.63 | 1.84 | 1.39 |
| GGPP | 8.44 | 3.22 | 1.29 | 0.91 |
| GFPP | 2.62 | 1.72 | 0.30 | 0.28 |
| Copalyl-PP | 2.37 | 1.39 | 0.38 | 0.38 |
| 2,3-epoxysqualene | 0.32 | 0.16 | 0.03 | 0.03 |
| 2xFPP | 0.62 | 0.16 | 0.05 | 0.04 |
| 2xGGPP | 0.82 | 0.31 | 0.08 | 0.09 |

### Supplementary References


77. Bansal, P. *et al.* Rhea, the reaction knowledgebase in 2022. *Nucleic Acids Res.* **50**, D693–D700 (2022).
78. Ashburner, M. *et al.* Gene ontology: tool for the unification of biology. The Gene Ontology Consortium. *Nat. Genet.* **25**, 25–29 (2000).
79. Gene Ontology Consortium. The Gene Ontology knowledgebase in 2026. *Nucleic Acids Res.* **54**, D1779–D1792 (2026).
80. BFD. https://bfd.mmseqs.com/.
81. Pundir, S., Martin, M. J. & O'Donovan, C. UniProt Protein Knowledgebase. *Methods Mol. Biol.* **1558**, 41–55 (2017).
82. Richardson, L. *et al.* MGnify: the microbiome sequence data analysis resource in 2023. *Nucleic Acids Res.* **51**, D753–D759 (2023).
83. Matasci, N. *et al.* Data access for the 1,000 Plants (1KP) project. *Gigascience* **3**, 17 (2014).
84. Goodstein, D. M. *et al.* Phytozome: a comparative platform for green plant genomics. *Nucleic Acids Res.* **40**, D1178–86 (2012).
85. Lesburg, C. A., Zhai, G., Cane, D. E. & Christianson, D. W. Crystal structure of pentalenene synthase: mechanistic insights on terpenoid cyclization reactions in biology. *Science* **277**, 1820–1824 (1997).
86. Tarshis, L. C., Proteau, P. J., Kellogg, B. A., Sacchettini, J. C. & Poulter, C. D. Regulation of product chain length by isoprenyl diphosphate synthases. *Proc. Natl. Acad. Sci. U. S. A.* **93**, 15018–15023 (1996).
87. Starks, C. M., Back, K., Chappell, J. & Noel, J. P. Structural basis for cyclic terpene biosynthesis by tobacco 5-epi-aristolochene synthase. *Science* **277**, 1815–1820 (1997).
88. Thoma, R. *et al.* Insight into steroid scaffold formation from the structure of human oxidosqualene cyclase. *Nature* **432**, 118–122 (2004).
89. DeLano, W. L. & Bromberg, S. PyMOL user's guide. *DeLano Scientific LLC* **629**, (2004).
90. Zhang, C., Shine, M., Pyle, A. M. & Zhang, Y. US-align: universal structure alignments of proteins, nucleic acids, and macromolecular complexes. *Nat. Methods* **19**, 1109–1115 (2022).
91. Barozet, A., Chacón, P. & Cortés, J. Current approaches to flexible loop modeling. *Curr Res Struct Biol* **3**, 187–191 (2021).
92. Virtanen, P. *et al.* SciPy 1.0: fundamental algorithms for scientific computing in Python. *Nat. Methods* **17**, 261–272 (2020).
93. Raffel, C. *et al.* Exploring the limits of transfer learning with a unified text-to-text transformer. *arXiv [cs.LG]* (2019) doi:10.48550/arXiv.1910.10683.
94. van Kempen, M. *et al.* Fast and accurate protein structure search with Foldseek. *Nat. Biotechnol.* **42**, 243–246 (2024).
95. Chicco, D. & Jurman, G. The advantages of the Matthews correlation coefficient (MCC) over F1 score and accuracy in binary classification evaluation. *BMC Genomics* **21**, 6 (2020).
96. Guo, C., Pleiss, G., Sun, Y. & Weinberger, K. Q. On calibration of modern neural networks. *arXiv [cs.LG]* (2017) doi:10.48550/arXiv.1706.04599.
97. Platt, J. Probabilistic Outputs for Support Vector Machines and Comparisons to

Regularized Likelihood Methods. *Adv. Large Margin Classif* **10**, 61–74 (1999).
98. Finn, R. D., Clements, J. & Eddy, S. R. HMMER web server: interactive sequence similarity searching. *Nucleic Acids Res.* **39**, W29–37 (2011).
99. Meier, J. *et al.* Language models enable zero-shot prediction of the effects of mutations on protein function. *bioRxiv* (2021) doi:10.1101/2021.07.09.450648.
100. Katoh, K. & Toh, H. Recent developments in the MAFFT multiple sequence alignment program. *Brief. Bioinform.* **9**, 286–298 (2008).
101. Price, M. N., Dehal, P. S. & Arkin, A. P. FastTree 2--approximately maximum-likelihood trees for large alignments. *PLoS One* **5**, e9490 (2010).
102. Aaron, J. A., Lin, X., Cane, D. E. & Christianson, D. W. Structure of epi-isozizaene synthase from Streptomyces coelicolor A3(2), a platform for new terpenoid cyclization templates. *Biochemistry* **49**, 1787–1797 (2010).
103. Lee, M. E., DeLoache, W. C., Cervantes, B. & Dueber, J. E. A Highly Characterized Yeast Toolkit for Modular, Multipart Assembly. *ACS Synth. Biol.* **4**, 975–986 (2015).
104. Kim, C. Y. *et al.* The chloroalkaloid (-)-acutumine is biosynthesized via a Fe(II)- and 2-oxoglutarate-dependent halogenase in Menispermaceae plants. *Nat. Commun.* **11**, 1867 (2020).
105. Lõoke, M., Kristjuhan, K. & Kristjuhan, A. Extraction of genomic DNA from yeasts for PCR-based applications. *Biotechniques* **50**, 325–328 (2011).
106. Gietz, R. D. & Schiestl, R. H. High-efficiency yeast transformation using the LiAc/SS carrier DNA/PEG method. *Nat. Protoc.* **2**, 31–34 (2007).
107. Schmid, R. *et al.* Integrative analysis of multimodal mass spectrometry data in MZmine 3. *Nat. Biotechnol.* (2023) doi:10.1038/s41587-023-01690-2.
108. Svoboda, A. *et al.* A miniaturized, high-throughput aqueous solvent-centric method for protein solubility screening. *Biochemistry* **65**, 1755–1762 (2026).
109. Robert, X. & Gouet, P. Deciphering key features in protein structures with the new ENDscript server. *Nucleic Acids Res.* **42**, W320–4 (2014).
110. Hwang, W. *et al.* CHARMM at 45: Enhancements in accessibility, functionality, and speed. *J. Phys. Chem. B* **128**, 9976–10042 (2024).
111. Im, W., Lee, M. S. & Brooks, C. L., 3rd. Generalized born model with a simple smoothing function. *J. Comput. Chem.* **24**, 1691–1702 (2003).
112. Huang, J. *et al.* CHARMM36m: an improved force field for folded and intrinsically disordered proteins. *Nat. Methods* **14**, 71–73 (2017).
113. Vanommeslaeghe, K. *et al.* CHARMM general force field: A force field for drug-like molecules compatible with the CHARMM all-atom additive biological force fields. *J. Comput. Chem.* **31**, 671–690 (2010).
114. Raz, K. *et al.* The impression of a nonexisting catalytic effect: The role of CotB2 in guiding the complex biosynthesis of cyclooctat-9-en-7-ol. *J. Am. Chem. Soc.* **142**, 21562–21574 (2020).
115. Lau, A. M., Kandathil, S. M. & Jones, D. T. Merizo: a rapid and accurate protein domain segmentation method using invariant point attention. *Nat. Commun.* **14**, 1–11 (2023).
